# Eco-evolutionary dynamics for finite populations and the noise-induced reversal of selection

**DOI:** 10.1101/2024.02.19.580940

**Authors:** Ananda Shikhara Bhat, Vishwesha Guttal

## Abstract

Theoretical studies from diverse areas of population biology have shown that demographic stochasticity can substantially impact evolutionary dynamics in finite populations, including scenarios where traits that are disfavored by natural selection can nevertheless increase in frequency through the course of evolution. Here, we analytically describe the eco-evolutionary dynamics of finite populations from demographic first principles to investigate how noise-induced effects can alter the evolutionary fate of populations in which total population size may vary stochastically over time. Starting from a generic birth-death process describing a finite population of individuals with discrete traits, we derive a set of stochastic differential equations (SDEs) that recover well-known descriptions of evolutionary dynamics such as the replicator-mutator equation, the Price equation, and Fisher’s fundamental theorem in the infinite population limit. For finite populations, our SDEs reveal how stochasticity can predictably bias evolutionary trajectories to favour certain traits, a phenomenon we call ‘noise-induced biasing’. We show that noise-induced biasing acts through two distinct mechanisms that we call the ‘direct’ and ‘indirect’ mechanisms. While the direct mechanism can be identified with classic bet-hedging theory, the indirect mechanism is a more subtle consequence of frequency and density-dependent demographic stochasticity. Our equations reveal that noise-induced biasing may lead to evolution proceeding in a direction opposite to that predicted by natural selection in the infinite population limit. By extending and generalizing some standard equations of population genetics, we thus describe how demographic stochasticity appears alongside and interacts with the more well-understood forces of natural selection and neutral drift to determine the eco-evolutionary dynamics of finite populations of non-constant size.

## Introduction

Eco-evolutionary population biology has a strong mathematical underpinning and can broadly be captured mathematically via a small number of equations such as the replicator-mutator equation and Price equation (Page and Nowak, 2002; Queller, 2017; Lion, 2018). The Price equation partitions changes in population composition into multiple terms, each of which lends itself to a straightforward interpretation in terms of the high-level evolutionary forces of selection and mutation, thus providing a useful mathematical framework for describing how populations change over time (Frank, 2012). The Price equation also leads to a number of simple yet insightful ‘fundamental theorems’ of population biology and unifies several seemingly disjoint formal structures under a single theoretical banner (Queller, 2017; Lion, 2018; Lehtonen, 2020a; Luque and Baravalle, 2021). However, the replicator-mutator equation, Price equation, and related ‘fundamental theorems’ of evolutionary dynamics are usually formulated in a deterministic setting that neglects stochastic fluctuations due to finite population effects (Page and Nowak, 2002; Queller, 2017; Lion, 2018).

Today, we increasingly recognize that incorporating the finite and stochastic nature of the real world routinely has much stronger consequences than simply ‘adding noise’ to deterministic expectations and can cause qualitative changes in the behavior of diverse biological systems (Horsthemke and Lefever, 1984; Black and McKane, 2012; Boettiger, 2018; Jhawar et al., 2020; Majumder et al., 2021; DeLong and Cressler, 2023; Yamamichi et al., 2023; Wang et al., 2023). In ecology and evolution, stochastic models need not exhibit phenomena predicted by their deterministic analogues (Proulx and Day, 2005; Johansson and Ripa, 2006; Black and McKane, 2012; Débarre and Otto, 2016). They may also exhibit novel phenomena not predicted by deterministic models (Constable et al., 2016; Rogers and McKane, 2015; Joshi and Guttal, 2018; DeLong and Cressler, 2023; Wang et al., 2023).

A striking example of such novel phenomena is the complete ‘reversal’ of evolutionary trajectories (relative to the expectations of infinite population models) that is seen in some finite population eco-evolutionary models (Houchmandzadeh and Vallade, 2012; Constable et al., 2016; McLeod and Day, 2019a; Mazzolini and Grilli, 2023). For example, in public goods games, the production of a costly public good is susceptible to invasion by ‘cheaters’ who use the public good but do not produce it. Due to this, standard (deterministic) evolutionary game theory predicts that producers should eventually go extinct in well-mixed populations. However, in finite, fluctuating populations, producers not only persist but also outcompete non-producers, the exact opposite of infinite population predictions (Constable et al., 2016; McLeod and Day, 2019a). This phenomenon of evolution proceeding in the direction of the classically disfavored type that leads to the ‘reversal’ of the prediction of deterministic natural selection has been dubbed ‘noise-induced selection’ (Week et al., 2021). Such noise-induced effects have been seen in several models in fields as diverse as sex chromosome evolution (Veller et al., 2017; Saunders et al., 2018), cell-cycle dynamics (Wodarz et al., 2017), social evolution (Houchmandzadeh and Vallade, 2012; Chotibut and Nelson, 2015; Constable et al., 2016; McLeod and Day, 2019a), and epidemiology (Kogan et al., 2014; Humplik et al., 2014; Parsons et al., 2018; McLeod and Day, 2019b; Day et al., 2020).

Despite the ubiquity of the phenomenon of qualitative noise-induced effects on evolutionary trajectories, we currently lack a description of how classic equations of evolutionary biology, such as the replicator-mutator equation, Price equation and Fisher’s fundamental theorem, are affected by such demographic stochasticity. Bet-hedging theory, a branch of evolutionary ecology that aims to build general theories that capture the effects of stochasticity on eco-evolutionary dynamics (Seger and Brockmann, 1987; Frank and Slatkin, 1990; Starrfelt and Kokko, 2012), has typically worked with both demographic and environmental stochasticity (Gillespie, 1977; Seger and Brockmann, 1987; Frank and Slatkin, 1990; Olofsson et al., 2009; Childs et al., 2010; Starrfelt and Kokko, 2012), two distinct forms of stochasticity that are qualitatively different in both origin and effect (Lande, 1993; Shoemaker et al., 2020). On the other hand, models of ‘noise-induced’ effects and ‘noise-induced selection’ model stochasticity as arising from inherent probabilistic nature of birth and death of organisms and are thus only concerned with demographic stochasticity (Parsons et al., 2010; Houchmandzadeh and Vallade, 2012; Constable et al., 2016; Parsons et al., 2018; McLeod and Day, 2019a; Day et al., 2020). Due to this, it is often unclear *a priori* under what situations these noise-induced effects become important for evolutionary dynamics or how these effects interact with the more well-understood evolutionary forces of natural selection, mutation, and drift (Shoemaker et al., 2020; Yamamichi et al., 2023). For example, how does noise-induced selection interact with genetic drift, or indeed natural selection? Are ‘noise-induced selection’ and ‘bet-hedging’ essentially the same effect that has been spoken about using different terminology (Parsons et al., 2010), or are there multiple distinct phenomena at play (Wang et al., 2023)?

In this paper, we derive general equations for the dynamics of finite, fluctuating populations evolving in continuous time starting from mechanistic first principles (Fig 1) via a stochastic birth-death process. By starting from ecological individual-level rules for birth and death and systematically describing population-level dynamics (Fig 1), we relax the assumption of constant (effective) population size that appears in classic finite population models of evolution such as the Wright-Fisher or Moran models (Lambert, 2010). Such a mechanistic approach is also thought to be a more fundamental description of eco-evolutionary dynamics (Lambert, 2010; Doebeli et al., 2017). The equations we derive reduce to well-known results such as the replicator-mutator equation and the Price equation in the infinite population limit, thus illustrating consistency with the known formal structures of eco-evolutionary population dynamics (Queller, 2017; Lion, 2018). For finite populations, these same equations also provide a generic description and synthesis of the noise-induced effects of finite population size and their consequences for eco-evolutionary population dynamics.

**Figure 1:**
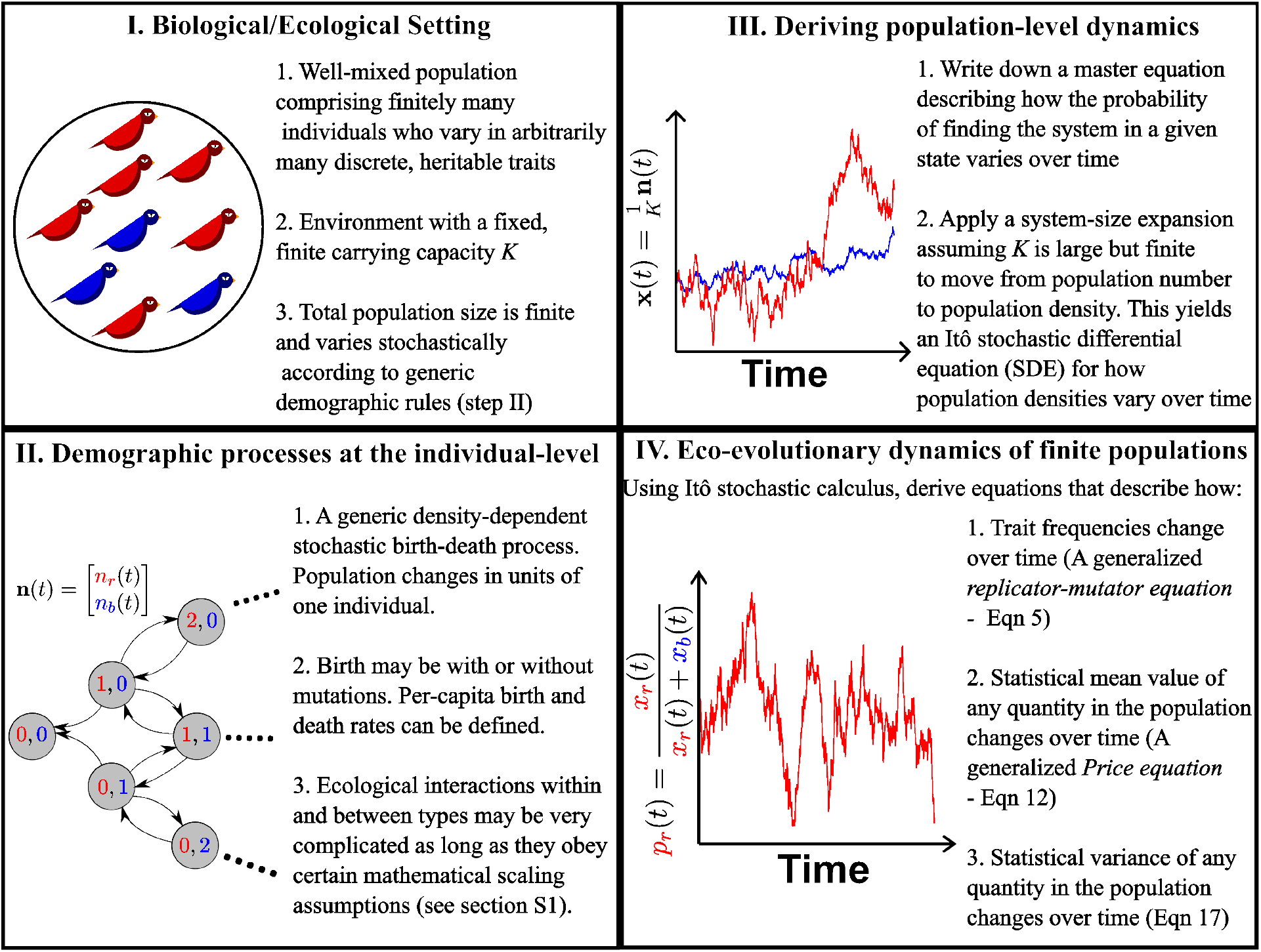
An outline of the approach we adopt in this paper.

Our systematic derivation provides relations between ecological quantities such as the expected population growth rate and the variance in population growth rate and connects them to evolutionary forces such as natural selection and genetic drift in trait frequency space. Using these equations, we synthesize the connections between noise-induced effects on population dynamics, including the ‘Gillespie effect’ of bet-hedging theory (Gillespie, 1977), ‘noise-induced effects’ in ecological population models (Constable et al., 2016; Parsons et al., 2018), ‘drift-induced selection’ (Veller et al., 2017; Saunders et al., 2018), ‘noise-induced selection’ (Week et al., 2021), and the effects of ‘evolutionary noise’ (McLeod and Day, 2019a; McLeod and Day, 2019b).

### A stochastic birth-death process for population dynamics

We consider a well-mixed population that can contain up to *m* different types of individual entities. For example, a gene may have *m* different alleles, individuals within a species may come in one of *m* phenotypes, or a community may have *m* different species; we refer to each distinct variant of an entity as a ‘type’. Unlike many classic stochastic formulations in evolutionary theory (Crow and Kimura, 1970; Lande, 1976; Kimura and Ohta, 1974), we do not assume a fixed or deterministically varying (effective) population size. Instead, we allow the total population size to emerge naturally, and thus fluctuate stochastically, from the stochastic birth and death processes (Fig 1).

#### Description of the process

Given a population that can contain up to *m* different kinds of entities, it can be completely characterized by specifying the number of individuals of each type of entity. Thus, the state of the population at a given time *t* is an *m*-dimensional vector of the form **n** = [*n*_1_(*t*), *n*_2_(*t*), …, *n*_*m*_(*t*)]^T^, where *n*_*i*_(*t*) is the number of individuals of type *i*.

We assume that the birth and death rate of each type in the population depends only on the state of the population (the vector **n**), and thus neglect any potential contributions from a temporally varying external environment. Our model unfolds in continuous time, and we assume that the probability of two or more births (or deaths) occurring at the same instant is negligible. For each type *i* ∈ {1, 2, …, *m*}, we denote the birth rate and the death rate by *b*_*i*_(**n**) and *d*_*i*_(**n**), respectively. We assume that the birth and death rates at the population level scale with the total population size such that *b*_*i*_(**n**) and *b*_*i*_(**n**) are of the order of ∑_*i*_ *n*_*i*_. Further, we assume that there exists a *carrying capacity* or, more generally, a *population size measure* (Czuppon and Traulsen, 2021) *K* > 0 that imposes a bound on population growth rate such that the growth rate of the total population size ∑_*i*_ *n*_*i*_ is expected to be negative whenever ∑_*i*_ *n*_*i*_ > *K* (Box 1).

##### Box 1

**Assumptions on the birth and death rates**

###### Scaling assumptions

Mathematically, we assume that we can find 𝒪(1) functions 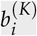 and 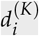 such that we can write

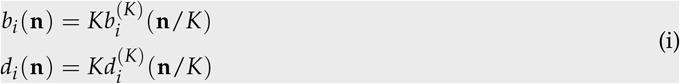

We can now define a notion of population density **x** = **n**/*K* by dividing the population number by the population size measure. We assume the stochastic process scales such that population densities remain well-defined in the infinite population size limit (*K* → ∞). Thus, we consider the limit of infinite population sizes but finite population densities, the usual domain of deterministic equations of population biology such as the Lotka-Volterra equation and logistic equation. We explain the concept of the infinite population size limit in more detail in Supplementary section S1.2.

###### Functional forms and per-capita rates

We assume that the birth and death rate functions have the functional form

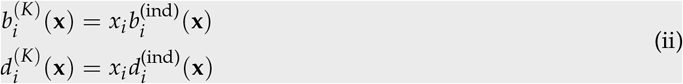

where 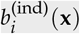 and 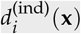 are non-negative functions that respectively describe the per-capita birth and death rate of type *i* individuals. In general, the birth rate of type *i* individuals may contain a component that does not depend purely multiplicatively on the current density *x*_*i*_ of type *i*: For example, when *x*_*i*_ = 0, *i.e*. there are no type *i* individuals in the population, individuals of type *i* may still be born through mutations of other types or immigration from other sources (gene flow). We account for this possibility via an additional ‘influx’ term in the Supplementary (section S1.1). Since such an influx term is not majorly affected by stochasticity (Supplementary section S2), we do not include it in the main text for the sake of conceptual clarity.

We emphasize that these birth and death rates can incorporate complicated interactions, but as we will see, the particular forms of these rate functions do not matter for our purposes as long as the mathematical scaling assumption in Eq. i are met.

Given the per-capita birth rates 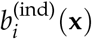 and per-capita death rates 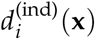 of each type (Box 1), we define the *Malthusian fitness* of the *i*^th^ type as

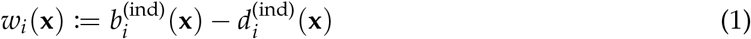

and the *per-capita turnover rate* of the *i*^th^ type as

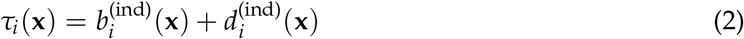

The quantity *w*_*i*_(**x**) describes the per-capita growth rate of type *i* individuals in a population **x**, and *τ*_*i*_(**x**) scribes the total rate of stochastic changes (through both births and deaths) to the density of type *i* individuals. It is notable that both *w*_*i*_ and *τ*_*i*_ depend on the state of the population as a whole (*i.e*. **x**) and not just on the density of the focal type. Thus, in general, both the fitness and the turnover rate in our model may be both density-dependent and frequency-dependent.

### Fundamental equations of eco-evolutionary dynamics

#### Ecological dynamics: Changes in population density

Having described the key demographic processes via a generic birth and death process, we now proceed to understand how the population density vector **x** changes over time.

Recall that the stochastic birth-death process changes in units of 1/*K* in density space. Thus, if *K* is large, each individual contributes a negligible amount to the population density, and the discontinuous jumps due to individual-level births or deaths in units of 1/*K* can be approximated as small, *continuous* changes in population density **x**. In Supplementary section S1, we use a formal version of this intuitive idea via a ‘system size expansion’ (Ethier and Kurtz, 1986, Chapter 11; Van Kampen, 1981, Chapter 10; Black and McKane, 2012; Czuppon and Traulsen, 2021) to derive a continuous description of the stochastic process for population densities. This continuous description takes the form of an Itô stochastic differential equation (SDE) which says that the density of the *i*^th^ type changes according to

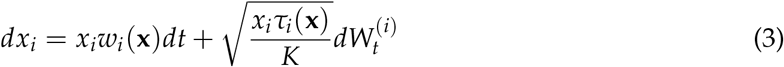

where each 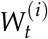 is a one-dimensional Wiener process (standard Brownian motion).

The first and second terms on the RHS of Eq. 3 respectively provide the so-called ‘infinitesimal mean’ and ‘infinitesimal variance’ of the stochastic process *x*_*i*_(*t*) that satisfies Eq. 3 (Karlin and Taylor, 1981; Czuppon and Traulsen, 2021). Informally, the infinitesimal mean and variance can be understood as follows: If we imagine that the population density of type *i* changes from *x*_*i*_ to *x*_*i*_ + *dx*_*i*_ over a very small (infinitesimal) time interval *dt*, we can (informally) view *dx*_*i*_ as a random variable. In that case, the *expected* density change 𝔼[*dx*_*i*_] and the *variance* in the change 𝕍[*dx*_*i*_] are respectively given by:

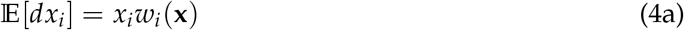

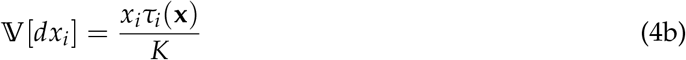

Thus, the Malthusian fitness *w*_*i*_ controls the expected change in population density, whereas the turnover rate *τ*_*i*_ (which is also a measure of the total number of events experienced by type *i* in a given time interval) controls the variance in the change in population density.

Eq. 3 describes the ecological population dynamics. To study the evolutionary dynamics of finite populations, we need to move from population densities to trait frequencies by defining some statistical quantities to describe how traits are distributed in the population (Box 2). We will see that this seemingly innocuous observation has important consequences when population size is non-constant.

##### Box 2

**Statistical measures for population-level quantities**

Given any state **x**(*t*) that describes our population at time *t*, let us first define the total (scaled) population size (*N*_*K*_ (*t*)) and the frequency *p*_*i*_ (*t*) of each type *i* in the population at time *t* as:

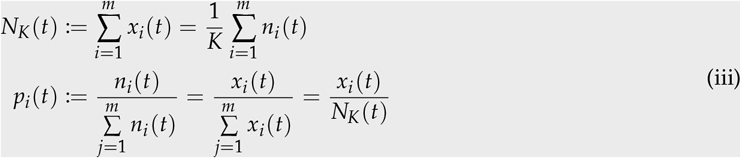

*N*_*K*_ here is an 𝒪(1) quantity and *KN*_*K*_ is the total population size, which is 𝒪(*K*). Since *N*_*K*_ is the sum of *m* stochastically fluctuating quantities, the total population size *KN*_*K*_ also experiences stochastic fluctuations and is thus non-constant in our model. We use the term ‘fluctuating populations’ henceforth to refer to populations of non-constant size that experience stochastic fluctuations in this manner.

Note that the frequency vector is subject to the constraint ∑_*i*_ *p*_*i*_ = 1, and we thus only need to study the system using the *m* variables [*p*_1_, *p*_2_, …, *p*_*m*−1_, *N*_*K*_]. We are often interested in tracking the effects of evolution on quantities described at a population level. To facilitate this, let *f* be any quantity that can be defined at the type-level, such as phenotype or fitness, with a (possibly time-dependent) value *f*_*i*_ ∈ ℝ for the *i*^th^ type. Recall that we defined *m* discrete types in the population on the basis that individuals within each type can be approximated as identical. Now, the statistical mean value of such a quantity in the population [*p*_1_, *p*_2_, …, *p*_*m*_], which we denote by 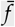, is given by

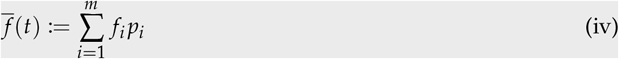

while the statistical covariance of two such quantities *f* and *g* in the population is given by

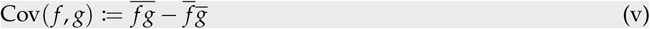

Lastly, the statistical variance of a quantity *f* in the population is given by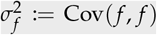. It is important to recognize that these statistical quantities are distinct from and independent of the *probabilistic* expectation, variance, and covariance obtained by integrating over realizations in the underlying probability space. We will denote this latter expectation and variance by 𝔼[·] and 𝕍[·] respectively for clarity.

#### Replicator equation for finite fluctuating populations

We now use Itô calculus to derive equations for the evolutionary dynamics of trait frequencies from Eq. 3, our SDE for population densities. Letting 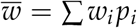 and 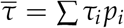 be the average population fitness and the average population turnover respectively, we show in Supplementary section S2 that *p*_*i*_, the frequency of the *i*^th^ type in the population **x**(*t*), changes according to the equation (also see Parsons et al., 2010, Eq. 7; Kuosmanen et al., 2022, Eq. 1):

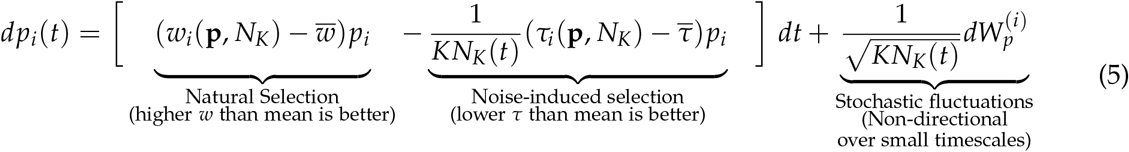

where 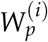 is a stochastic integral term given by

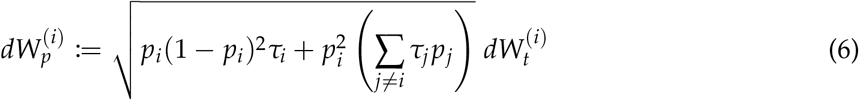

and each 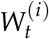 is a one-dimensional Wiener process. An analogous equation has also been derived for stochastic SIR systems (Parsons et al., 2018, Eq. 2.5). The first term of Eq. 5 represents the effect of natural selection for increased (Malthusian) fitness. Eq. 5 when derived with mutation terms (Eq. S29) recovers the replicator-mutator equation (Eq. 6 in Lion, 2018) in the infinite population (*K* → ∞) limit (see section S7 in the supplementary), and without mutation, recovers the standard replicator equation.

Importantly, finite populations experience a directional force dependent on *τ*_*i*_(**x**), the percapita turnover rate of type *i*, that cannot be captured in infinite population models but appears in the second term on the RHS of Eq. 5 (Parsons and Quince, 2007; Parsons et al., 2010; Week et al., 2021; Kuosmanen et al., 2022; Bhat, 2024). This term shows that the effect of turnover rates is structurally identical to that of the differential fitness, but it acts in the opposite direction - a higher relative *τ*_*i*_ leads to a decrease in frequency (notice the minus sign before the second term on the RHS of Eq. 5). For this reason, the effect has been termed ‘noise-induced selection’ (Week et al., 2021), though similar ideas have been known under the names ‘bet-hedging’ and ‘Gillespie effect’ in the evolutionary ecology literature (Gillespie, 1974; Gillespie, 1977; Frank and Slatkin, 1990; Starrfelt and Kokko, 2012; Veller et al., 2017; See Box 4). The same effect has also been noticed in the epidemiology literature, where the term we call ‘turnover’ corresponds to the virulence of the pathogen strain (Kogan et al., 2014; Parsons et al., 2018; Day et al., 2020). Noise-induced selection can be heuristically understood as a stochastic selection for reduced variance in changes in population density (Box 3).

Finally, the last term describes the effects of stochastic fluctuations due to the finite size of the population and shows the 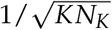 scaling that is typical of demographic stochasticity. Though this last term vanishes upon taking probabilistic expectations (and is hence ‘non-directional’ in the short term), it may bias trait frequency distributions by affecting the amount of time spent in different states, as we illustrate in the next section.

To complete the description of the system, we also require an equation for the total scaled population size *N*_*K*_ = ∑ *x*_*i*_. Upon noting that *dN*_*K*_ = ∑ *dx*_*i*_, using Eq. 3 for *dx*_*i*_, and dividing both sides by *N*_*K*_ we find

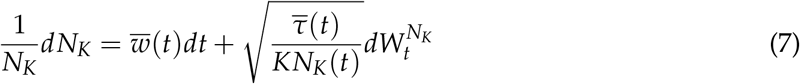

where 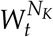 is a one-dimensional Wiener process and we have used the representation of noise terms presented in Supplementary section S5. Thus, fitness affects only the infinitesimal mean and turnover rate affects only the infinitesimal variance of the total population size. Note that the LHS of Eq. 7 is simply the rate of change of log(*N*_*K*_), i.e. the rate of change of the (scaled) population size *N*_*K*_ when viewed on a logarithmic scale.

#### A special case: Two interacting types

To illustrate the way stochasticity affects evolutionary dynamics in finite, fluctuating populations, we consider the simple case of two interacting types (i.e. *m* = 2). Letting *p* = *p*_1_ be the frequency of type 1 individuals in the population, we see from Eq. 5 that our two-type population obeys the equations

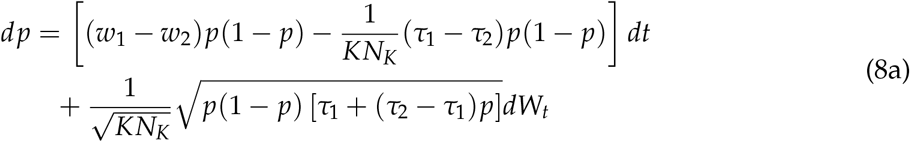

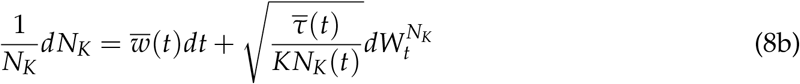

where *W*_*t*_ and 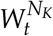 are one-dimensional Wiener processes. We can now identify the (frequency-dependent) selection coefficient *s*(*p, N*_*K*_) := *w*_1_(*p, N*_*K*_) − *w*_2_(*p, N*_*K*_) from classic population genetics. The selection coefficient quantifies the direction and strength of natural selection in the system — a positive (or negative) value of *s* indicates that type 1 individuals are favoured (disfavoured) by natural selection.

Eq. 8a also motivates the definition of an analogous noise-induced selection coefficient *κ*(*p, N*_*K*_) := *τ*_2_(*p, N*_*K*_) − *τ*_1_(*p, N*_*K*_) to quantify the direction and strength of noise-induced selection. If type 1 has a lower turnover rate, *κ*(*p, N*_*K*_) > 0, and thus type 1 is favoured by noise-induced selection.

With this notation, Eq. 8a becomes

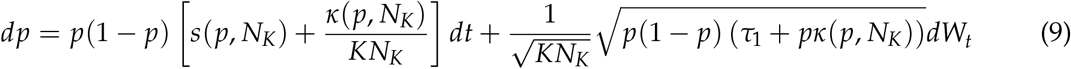

where we see that the selection coefficient *s*(*p, N*_*K*_) affects the infinitesimal mean (*dt* term) of Eq. 9, and the noise-induced selection coefficient *κ*(*p, N*_*K*_) affects both the infinitesimal mean and the infinitesimal variance. Note that fitness only enters into the population dynamics via the selection coefficient *s*, whereas turnover also appears via *τ*_1_ in the second term on the RHS of Eq. 9. In other words, only the relative fitness or the difference *w*_1_ − *w*_2_, but not the absolute value of the fitness *w*_*i*_, matters for the deterministic dynamics. In contrast, the functional form and absolute value of the per-capita turnover rate *does* affect the stochastic dynamics of the system via the second term on the RHS of Eq. 9.

##### Box 3

**A heuristic explanation of noise-induced selection**

One key mechanism through which demographic stochasticity can affect evolutionary dynamics is by biasing evolutionary trajectories towards types with lower turnover rates, even if these types have the same (or even lower) fitness than other types in the population. Here, we explain this mechanism via an intuitive argument that has the same flavour as arguments seen in the bet-hedging literature (Gillespie, 1977; Frank and Slatkin, 1990; Starrfelt and Kokko, 2012).

To illustrate the idea via an example, imagine a system consisting of two types of individuals, 1 and 2, which have equal fitness but unequal turnover rates; without loss of generality, assume *τ*_1_ > *τ*_2_. Let us further assume that both types have the same density *x*_0_. From Eq. (4), we see that, in our example, though the two types of individuals have the *same expected change* in population density, type 1 individuals have a *greater variance* in the changes in density than type 2 individuals.

Since evolution is defined as changes in trait frequencies, we transform variables from population density to trait frequency to see how differential variance affects evolutionary trajectories. This is done via the transformation

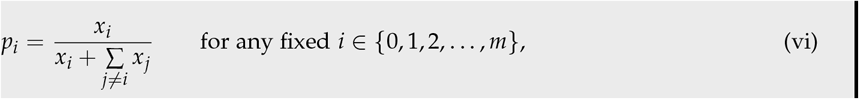

Observe now that frequency (*p*_*i*_) is a *concave* function of density *x*_*i*_ (Eq (vi)). Due to concavity, equivalent changes in density do not correspond to equivalent changes in frequency. Instead, a result mathematically known as Jensen’s inequality and diagrammatically represented in figure 2 applies.

**Figure 2:**
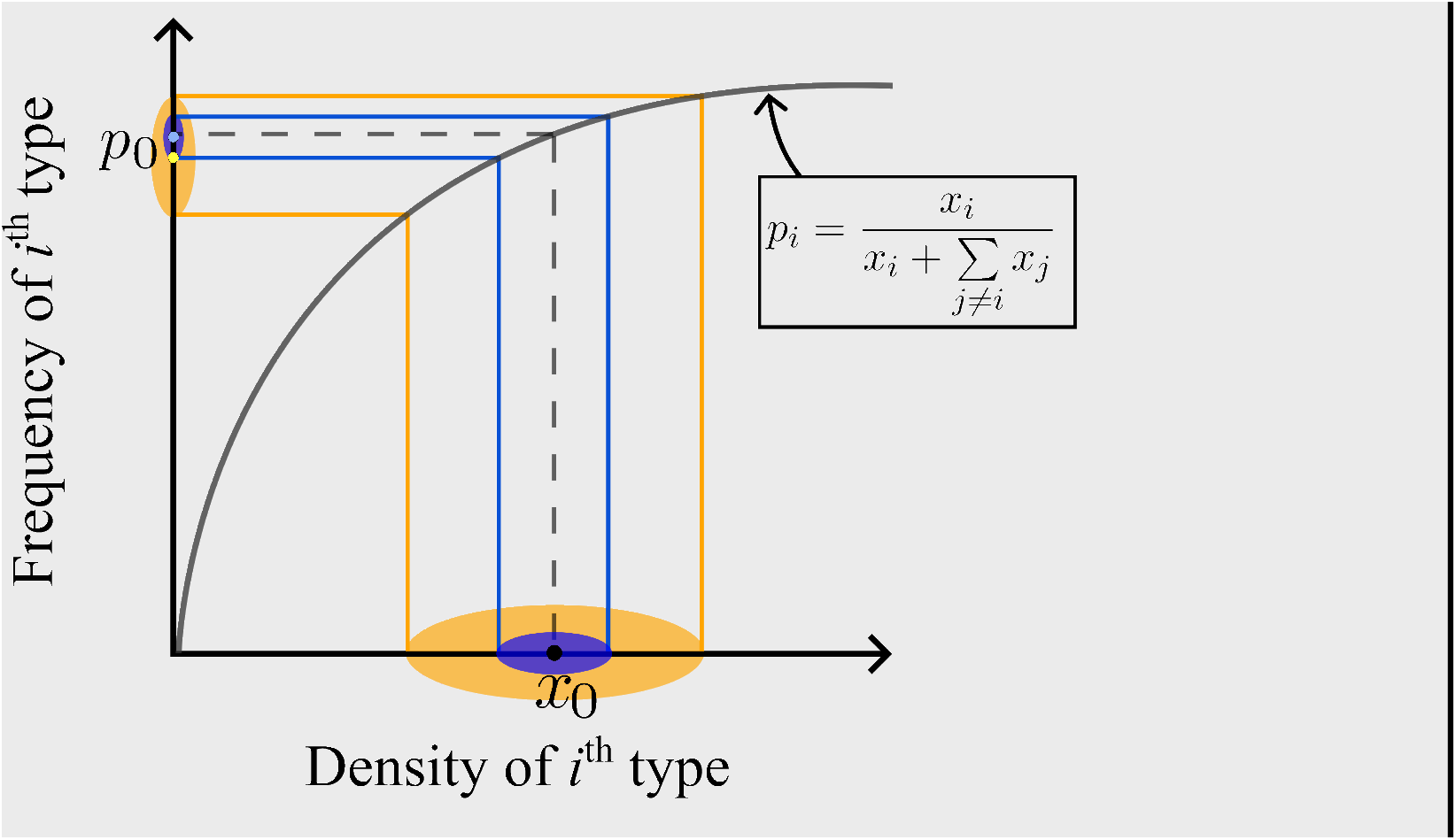
A diagrammatic representation of the consequences of demographic stochasticity when total population size can vary. The grey curve represents the transformation from population densities to trait frequencies via Eq. (vi). The ellipses are representations of possible changes in population composition for two types with the same fitness and same initial density, but different variances (yellow > blue). The center of the ellipse represents the infinitesimal mean of the density process, the major axis captures the infinitesimal variance, and the colored region is thus representative of all possible changes given that an event (birth or death) has occurred. Reductions in density have a stronger effect on frequency than increases in density, and due to this, the expected frequency (centers of ellipses on the y-axis) after an event has occurred is less than the initial frequency *p*_0_ even if the expected density (centers of ellipses on the x-axis) coincides with the initial density *x*_0_. Types with a larger variance in the density process (yellow ellipse in the figure) experience a greater reduction in expected frequency relative to types with a lower variance (blue ellipse). Similar figures, with the X and Y axes being absolute fitness and relative fitness respectively, appear in expositions of bet-hedging (e.g. Frank and Slatkin, 1990; Starrfelt and Kokko, 2012); In our figure, the axes are population density and trait frequency respectively.

An increase in density leads to a relatively smaller increase in frequency, whereas an equivalent decrease in density leads to a larger decrease in frequency. This implies that stochastic reductions in density have a higher cost (decrease in frequency) than the benefits (increase in frequency) conferred by a numerically equivalent increase in density (Fig. 2). Thus, variance in the density process leads to a net cost in frequency space, and all else being equal, a greater variance comes with a greater cost. Types with lower turnover rates (corresponding to lower infinitesimal variance in Eq. (3)) are thus favoured.

The argument we provide here is particular to populations of non-constant size. To see this, assume that the total (scaled) population size ∑_*i*_ *x*_*i*_ is a constant *N* > 0. The transformation in Eq. (vi) then becomes

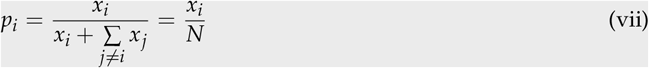

and is now simply a linear re-scaling of *x*_*i*_. The asymmetry between increases in density and decreases in density thus disappears. In other words, the mechanism that we identified above no longer operates for constant populations!

Demographic stochasticity can also affect population dynamics through the second term on the RHS of Eq. 9 due to turnover-dependent stochastic effects (McLeod and Day, 2019a). To study these effects, we will examine the *speed density m* (Karlin and Taylor, 1981; Czuppon and Traulsen, 2021) of the stochastic process described by Eq.9. As we explain in Supplementary section S6, the speed density *m*(*p*_0_) at the point *p*_0_ is a measure of the amount of time the population spends in the ‘immediate neighbourhood’ of the state *p*_0_ (formally, it is proportional to the amount of time spent in the interval (*p* − *ϵ, p* + *ϵ*) in the limit *ϵ* → 0; see Karlin and Taylor, 1981, Chapter 15, Remark 3.2). When a stationary distribution or quasi-stationary distribution (Collet et al., 2013) exists, it is proportional to the speed density (Karlin and Taylor, 1981, Chapter 15, Equation 5.34 along with Chapter 15, Equation 3.10; Collet et al., 2013, theorem 6.4) and the speed density thus describes the trait frequency distribution at (quasi)-stationarity in such cases. In supplementary section S6, we show (Eq. S83) that the speed density *m*(*p*) obeys the equation:

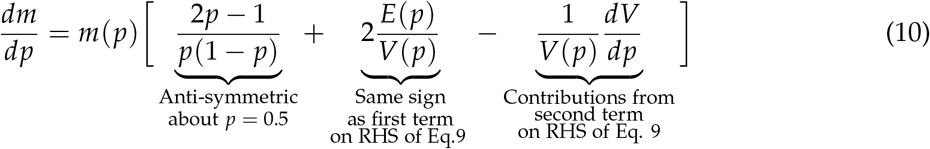

where

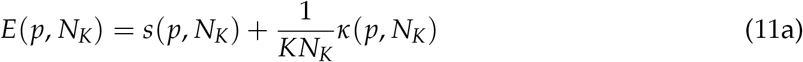

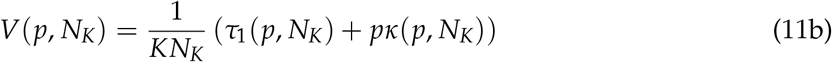

The sign of *dm*/*dp* tells us whether the system spends more time in states in which type 1 is overrepresented (positive meaning that type 1 is favoured), and points at which *dm*/*dp* = 0 tell us about the states which are most likely/least likely to be observed before fixation/extinction has taken place (McLeod and Day, 2019a; Majumder et al., 2021). The first term on the RHS of 10 is anti-symmetric about *p* = 0.5 and thus uninteresting on its own for determining the sign of *dm*/*dp* as a function of *p*.

The second term of Eq. 10 represents the balance between classical selection and noise-induced selection. Since both *s* and *κ* are 𝒪(1) functions, natural selection will tend to dominate *E*(*p*) when the total population size *KN*_*K*_ is large. Additionally, if *s* and *κ* are of similar magnitude (*i.e*. the strength of natural selection is comparable to the strength of the Gillespie effect), natural selection will still dominate the sign of *E*(*p*) since the total population size *KN*_*K*_ must be greater than 1. However, noise-induced selection can qualitatively affect evolutionary dynamics if differences in Malthusian fitnesses are close to zero (*i.e*. natural selection is weak, |*s*| ≪ 1) or if total population size *KN*_*K*_ is small (Parsons et al., 2010; Parsons et al., 2018). We also show this explicitly using an example in Box 5.

Eq. 10 also tells us that noise-induced selection (explained heuristically in Box 3) is not the only way in which demographic stochasticity can bias trait distributions: Instead, the speed density is also profoundly affected by the ‘noise’ terms in Eq. 9, as captured by the last term on the RHS of Eq. 10. In particular, even when the first term on the RHS of Eq. 9 vanishes or acts in the same direction as classical selection, populations may still spend much more time in states where a certain type is overrepresented, in particular possibly ‘reversing’ the prediction of infinite population models, if *dV*/*dp* is non-zero. For example, the system may spend much more time in configurations where type 1 individuals are overrepresented even if *s* + *κ*/*KN*_*K*_ < 0 (meaning that the first term on the RHS of 9 favours type 2 individuals) as long as *dV*/*dp* is sufficiently negative (McLeod and Day, 2019a). Thus, one type is ‘favoured’ through this effect in the sense that we are more likely to observe the population in states where the focal type is overrepresented, an effect that has been ascribed to ‘evolutionary noise’ (McLeod and Day, 2019a; McLeod and Day, 2019b). As an aside, note that 1/*VdV*/*dP* could also equivalently be written as the derivative of log(*V*) with respect to *p*, and thus represents the strength and direction of frequency dependence of log(*V*). Since 𝕍[*dp*] = *p*(1 − *p*)*V*(*p*) from Eq. 9, log(*V*) can be interpreted as being proportional to the logarithm of the variance in the changes in the trait frequency *dp*. This term thus captures the contributions of stochastic fluctuations/’noise’ in the trait frequency changes *dp* and can be interpreted as ‘selecting’ for reduced variance in the change in trait frequencies *dp*, whereas noise-induced selection is a selection for reduced variance in the change in population densities (Box 3). Both these effects can bias the distribution of types observed in finite populations, and we therefore collectively refer to the two effects as *noise-induced biasing*. Since noise-induced selection is directly visible as a deviation in the expected change in frequency 𝔼[*dp*], we call it the ‘direct’ mechanism. Since the term looks mathematically similar to the action of natural selection (compare first and second terms on the RHS of Eq. 5 or Eq. 9), we also use the phrase ‘noise-induced selection’ for the direct mechanism (following Week et al., 2021). In contrast, noise-induced biasing via frequency-dependence of *V* is a more subtle mechanism that affects the distribution of types indirectly by biasing the time spent in different states, and we thus refer to this effect as the *indirect* mechanism of noise-induced biasing (Box 4).

Remarkably, when natural selection does not operate (*s* = 0), there are situations where the speed density, and thus the stationary distribution when it exists, do not depend on the total population size. In particular, if *τ*_1_ and *κ* are such that the ratio *τ*_1_/*κ* is independent of the total population size *KN*_*K*_, then so is the speed density. Intuitively, this is because both noise-induced biasing and drift arise from the stochasticity associated with finite populations. More precisely, when *s* = 0, the total population size *KN*_*K*_ affects the dynamics only through a pre-factor of 1/*KN*_*K*_ that occurs in both *E*(*p*) and *V*(*p*). It therefore disappears in the ratio *E*/*V*. Thus, unlike the classic results regarding natural selection-drift balance, the total population size does *not* affect the relative strengths of noise-induced biasing and genetic drift — instead, it is the details of the demographic processes, as captured by *κ* and *V*, that determine which effect dominates. A similar observation has been made in life-history theory (Shpak, 2005).

##### Box 4

**Two distinct non-neutral effects of demographic stochasticity**

Demographic stochasticity can cause certain types to be systematically overrepresented in the population relative to infinite population expectations, even if the fitness of these focal types is the same as (or lower than) the fitness of other types in the population. Since such biases in the trait distribution are induced purely by stochasticity and do not appear in deterministic models, we call this phenomenon *noise-induced biasing*. Our equations reveal that noise-induced biasing can occur through two distinct mechanisms. In this box, we provide a summary of the connections and delineations between the two mechanisms.

1. The *direct mechanism* selects for reduced variance in changes in population density (Gillespie, 1974; Gillespie, 1977). This mechanism appears in the ‘deterministic’ term (*dt* term) of the replicator equation (Eq. 5) and is detectable as a systematic deviation of the expected trajectory 𝔼[*dp*/*dt*] from the infinite population prediction (Parsons et al., 2010; Parsons et al., 2018). The direct mechanism can be identified with the ‘Gillespie effect’ from the bet-hedging literature (Gillespie, 1974) and is obtained as a balance between natural selection for increased ecological growth rate and a stochastic selection for reduced variance in changes in population densities (see Box 3). Since the direct mechanism looks mathematically very similar to the force of natural selection (Compare first and second terms on RHS of Eq. 5), it has also been called *noise-induced selection* in the literature (Week et al., 2021). Noise-induced selection in this sense is thus a version of classical evolutionary bet-hedging (Frank and Slatkin, 1990; Starrfelt and Kokko, 2012) in an explicitly demographic, dynamical context. Note that unlike in bet-hedging models, *w* and *τ* (and thus *s* and *κ*) are derived from the underlying demographic processes.
2. The *indirect mechanism* appears as an apparent selection for reduced variance in changes in trait frequency (McLeod and Day, 2019a). The effects of demographic stochasticity, in this case, appear in the ‘stochastic’ term (*dW* term) of the replicator equation (Eq. 5) and affect the time spent at various configurations, and thus, indirectly, the probability of observing a polymorphic population in a particular configuration (*p*_1_, *p*_2_, …, *p*_*m*−1_, *N*_*K*_). The indirect mechanism results from frequency-dependence in the variance of changes in trait frequencies and can be thought of as analogous to frequency-dependent ‘viscosity’; populations tend to accumulate in those configurations that lead to slower changes in population composition, and we are thus more likely to observe the population in those configurations that make the rate of change of the population ‘slower’. The strength of indirect noise-induced biasing varies inversely with (the square root of) population size, and the direction of the effect depends on the frequency-dependence of the per-capita turnover rates *τ*_*i*_.

Unlike natural selection, the balance between noise-induced biasing (through either mechanism) and genetic drift in the absence of natural selection does *not* depend on the total population size: Instead, it is determined by the details of the demographic processes occurring in the population: If different types have different turnover rates, the direct mechanism (noise-induced selection) operates, and if some types are associated with lower variance in the change in trait frequencies, the indirect mechanism operates. Note that this observation means that noise-induced biasing, via both direct and indirect mechanisms, need not operate or be a significant force in small populations, depending on demographic details.

#### Price equation for finite fluctuating populations

We show that the statistical population mean 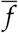 of any type-level quantity *f* (e.g. phenotype, fitness) changes over time according to the equation (see Supplementary section S3)

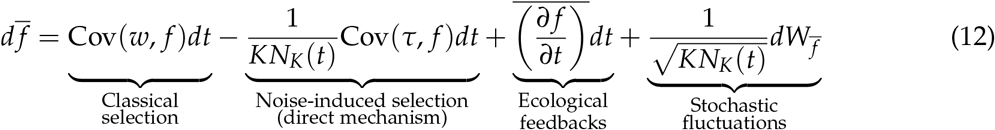

where

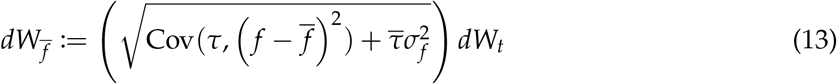

is a stochastic integral term describing un-directed stochastic fluctuations (see Eq. S67 in Supplementary section S5). Here, we use *W*_*t*_ to denote a generic Wiener process whose relation to the Wiener processes in Eq. 5 can be studied using a relation discussed in Supplementary section S5. Equation 12 has previously been derived in the epidemiology literature (Day et al., 2020, Eq. 5.2; see Section S8) and a quantitative trait version of the equation has also been derived using more sophisticated mathematical techniques (Week et al., 2021, Eq. 21b; Bhat, 2024, Eq. 25).

Eq. 12 recovers the Price equation (Eq. 11 in Lion, 2018) in the infinite population (*K* → ∞) limit (see section S7). Each term in Eq. 12 lends itself to a simple biological interpretation. The first term, Cov(*w, f*), is well-understood in the classical Price equation and represents the effect of natural selection. If the trait and the fitness are positively correlated, the mean trait value in the population increases due to the effect of selection. The second term, Cov(*τ, f*)/*KN*_*K*_(*t*) is the effect of noise-induced selection in finite fluctuating populations. Biologically, the Cov(*τ, f*) term (with negative sign) describes a biasing effect due to differential turnover rates between different types; if the trait is positively correlated with turnover rate, this term causes the mean trait value to decrease.

The third term of Eq. 12 is relevant whenever *f*_*i*_ can vary over time and represents feedback effects on the quantity *f*_*i*_ of the *i*^th^ species over short (‘ecological’) time-scales. Such a feedback could be through a changing environment, phenotypic/behavioral plasticity, or any manner of other ‘ecological’ phenomena. This is the term that captures eco-evolutionary feedback loops.

Finally, the last term of Eq. 12 describes the role of stochastic fluctuations. Recall that the square of this term corresponds to the infinitesimal variance of the change in the mean value 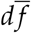 of the quantity *f* in the population. 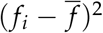 is a measure of the distance of *f*_*i*_ from the population mean 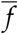. The Cov 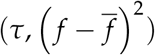 term thus tells us that if turnover *τ*_i_ of the *i*^th^ type covaries positively with the distance of *f*_*i*_ from the population mean (*i.e*. individuals with more extreme *f* have higher turnover rates), the population experiences a greater variance in 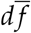, i.e. the change in the mean value of *f* over infinitesimal time intervals. The 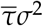 term tells us that even if *τ* and *f* do not covary, there is still some variance in 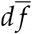, given now by the product of the mean turnover rate 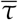 with the standing variation 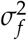 in the quantity *f*. As we shall see in the next section, this is a manifestation of neutral genetic/ecological drift. Just as in the replicator equation, stochastic fluctuations through 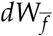 can profoundly affect the time spent at different values of 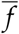 (and the stationary distribution, when it exists) via the indirect mechanism of noise-induced biasing if the term inside the square root of Eq. 13 depends on 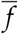. Note that unlike for the replicator equation, the SDE in Eq. 12 is one-dimensional regardless of the number of traits (*m*), and thus the stationary distribution of the mean value 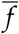 can always be studied the way we studied Eq. 9.

#### Fisher’s fundamental theorem for finite fluctuating populations

Two particularly interesting implications of Eq. 12 are realized upon considering the time evolution of mean fitness and mean turnover rate. First, upon substituting *f* = *w* in Eq. 12 and taking expectations over the underlying probability space, we obtain:

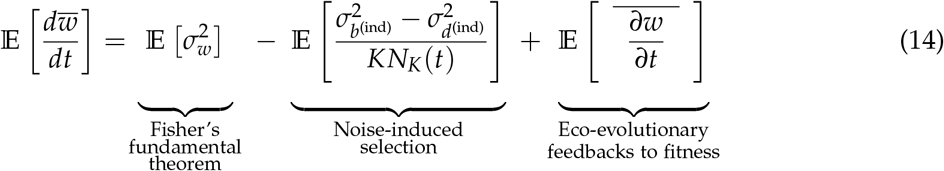

The first term, 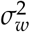, is the subject of Fisher’s fundamental theorem (Frank and Slatkin, 1992; Kokko, 2021), and says that in infinite populations, the rate of change of mean fitness in the population is proportional only to the standing variation in fitness 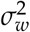 if fitness at the type level (*w*_*i*_) does not change over time. The second term of Eq. 14 is a manifestation of noise-induced selection acting and is particular to finite populations (note that the indirect mechanism does not operate because we are only looking at expectation values). Finally, the last term arises whenever *w*_*i*_ can vary over time (Kokko, 2021), be it through frequency-dependent selection, phenotypic plasticity, varying environments, or other ecological mechanisms, and represents feedback effects on the fitness *w*_*i*_ of the *i*^th^ species over short (‘ecological’) time-scales. Eq. 14 recovers the standard version of Fisher’s fundamental theorem in the infinite population (*K* → ∞) limit (see section S7).

#### The demographic origins of fitness differences induce quantitative corrections to Fisher’s fundamental theorem in finite populations

Since *w* = *b*^(ind)^ − *d*^(ind)^ by definition, Eq. 14 can alternatively also be written as

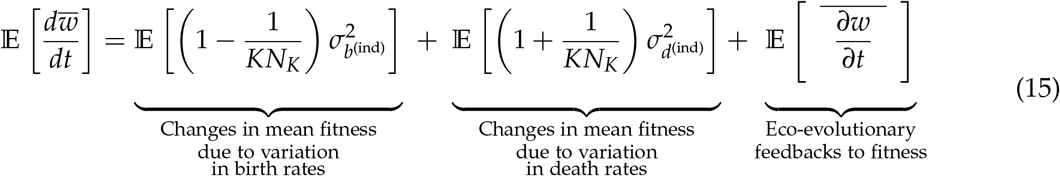

Eq. 15 redescribes variation in fitness in terms of the more fundamental processes of birth and death. Eq. 15 also tells us that variation in death rates leads to a slightly greater rate of increase in mean fitness than an equivalent variation in birth rates. For example, if individuals differ in birth rates alone 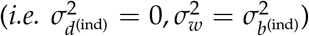, Eq. 15 predicts that the rate of mean fitness in the absence of eco-evolutionary effects is given by 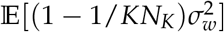. On the other hand, if individuals instead differ in death rates alone, 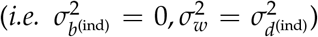, the rate of change of mean fitness in the absence of eco-evolutionary effects is given by 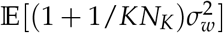, which is a slightly faster rate of change. Note, however, that these are only minor quantitative corrections to Fisher’s fundamental theorem and the two cases exhibit the same qualitative behaviour.

#### An analog of Fisher’s fundamental theorem for the mean turnover rate of the population

Carrying out the same steps in deriving Eq. 14 with *f* = *τ* in Eq. 12 yields a dynamical equation for the evolution of mean turnover rates (Kuosmanen et al., 2022) and reads

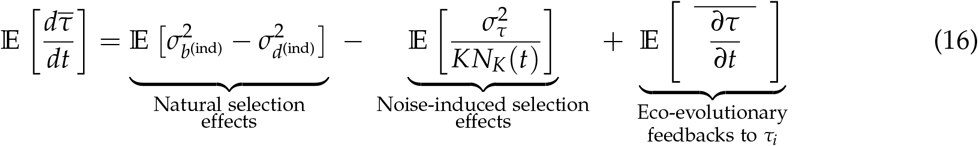

The first term captures the effect of natural selection on mean turnover rates and says that natural selection may either increase or decrease the mean turnover rate depending on the demographic details of the population. More precisely, we predict that natural selection is expected to increase the mean turnover rate in the population if (and only if) the expected variance in the birth rates is greater than the expected variance in the death rates (see also Kuosmanen et al., 2022). The second term of Eq. 16 represents the effect of noise-induced selection and is exactly analogous to the 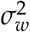 term in Fisher’s fundamental theorem. This term says that noise-induced selection always reduces mean turnover in the population, with the rate of reduction of the mean turnover rate being proportional to the standing variation in turnover rates 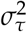. Finally, the last term on the RHS of Eq. 16 quantifies the effect of eco-evolutionary feedback via changes in the turnover of each type over time. In infinitely large populations (*K* → ∞), the second term on the RHS of Eq. 16 disappears; thus, the mean turnover rate 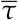 may either decrease or increase in infinitely large populations based on demographic details of (the variance of) birth and death rates in the population (Kuosmanen et al., 2022). In contrast, the noise-induced selection (second term) always reduces the mean turnover rate.

#### The fundamental equation for the population variance via a generalization of an equation for variance of type-level quantities

Eq. 12 is a general equation for the mean value of an arbitrary type level quantity *f* in the population. In many real-life situations, we are interested in not just the population mean, but also the variance of a quantity in the population. In Supplementary section S4, we show that the statistical variance of any type level quantity *f* obeys

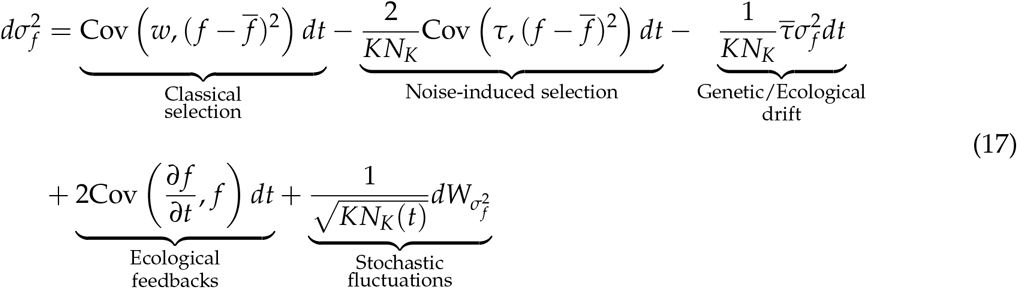

where

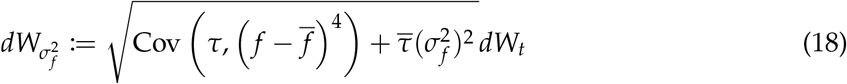

is a stochastic integral term measuring the (non-directional) effect of stochastic fluctuations that vanishes upon taking an expectation over the probability space (see Eq. S69 in Supplementary section S5). As before, we use *W*_*t*_ to denote a generic Wiener process — the *W*_*t*_ that appears in Eq. 17 is not necessarily the same process that appears in either Eq. 5 or Eq. 12. The stochastic dependencies between the various Wiener processes can be studied using a relation discussed in Supplementary section S5. An infinite population (*K* → ∞) version of Eq. 17 appears in Lion, 2018 (see Supplementary section S7) as a dynamic version of earlier, dynamically insufficient equations for the change in trait variation over a single generation (For example, see Eq. 6.14 in Rice, 2004)

Once again, terms of Eq. 17 lend themselves to straightforward biological interpretation. The quantity 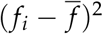 is a measure of the distance of *f*_*i*_ from the population mean value 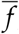, and thus covariance with 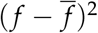 quantifies the type of selection operating in the system: A negative correlation is indicative of stabilizing or directional selection, and a positive correlation is indicative of disruptive (*i.e*. diversifying) selection (Rice, 2004, Chapter 6; Lion, 2018). An extreme case of diversifying selection for fitness occurs if the mean fitness of the population is at a local minimum but 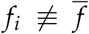 (*i.e*. the population still exhibits some variation in *f*). In this case, if the variation in *f* is associated with a variation in fitness, then Cov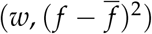 is strongly positive, and the population experiences a sudden explosion in variance, causing the emergence of polymorphism in the population. If Cov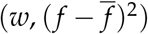 is still positive after the initial emergence of multiple morphs, evolution eventually leads to the emergence of stable coexisting polymorphisms in the population - this phenomenon is a slight generalization of the idea of evolutionary branching that occurs in frameworks such as adaptive dynamics (Doebeli, 2011). The Cov (*∂ f* /*∂t, f*) term represents the effect of eco-evolutionary feedback loops due to changes in *f* at the type level.

Finally, the last term on the RHS of Eq. 17 describes the role of stochastic fluctuations. The square of this term is the infinitesimal (probabilistic) variance of the changes in statistical variance 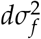 of *f*. Just like in the stochastic replicator and Price equations, this term can affect the time spent at different values of trait variance through the indirect mechanism of noise-induced biasing. Just like the stochastic Price equation, the SDE in Eq. 17 is always one-dimensional, and thus the stationary distribution of the variance 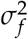 can also always be studied the way we studied Eq. 9.

In the case of one-dimensional quantitative traits, an infinite-dimensional version of Eq. 17 has recently been rigorously derived (Week et al., 2021) using measure-theoretic tools under certain additional assumptions (Week et al., 2021, Eq. (21c); see Appendix S9). Taking expectations over the probability space in Eq. 17 also recovers an equation previously derived and used (Débarre and Otto, 2016) in the context of evolutionary branching in finite populations as a special case (Equation A.23 in Débarre and Otto, 2016 is equivalent to our Eq. 17 for their choice of functional forms upon converting their change in variance to an infinitesimal rate of change *i.e*. derivative).

#### Loss of trait variation in populations experiencing drift

The 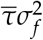 term quantifies the loss of variation due to stochastic extinctions (i.e. demographic stochasticity) and thus represents the classic effect of neutral drift in finite populations. Our equations are agnostic to whether each type *i* is an allele, a phenotype, a morph, or a species, so the drift in question may be either genetic or ecological drift depending on biological context. To see why 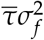 quantifies the loss of trait variation, it is instructive to consider the case in which this is the only force at play. Let us imagine a population of asexual organisms in which each *f*_*i*_ is simply a label or mark arbitrarily assigned to individuals in the population at the start of an experiment/observational study and subsequently passed on to offspring — for example, a neutral genetic tag in a part of the genome that experiences a negligible mutation rate. Since the labels are arbitrary and have no effect whatsoever on the biology of the organisms, each label has the same fitness *w*_*i*_ ≡ *w* and per-capita turnover *τ*_*i*_ ≡ *τ*, and thus 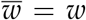 and 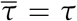. Note that since every type has the same fitness and turnover rate, we have Cov 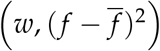 ≡ Cov 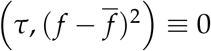. Since the labels do not change over time, we also have Cov (*∂ f* /*∂t, f*) = 0. From Eq. 17, we see that in this case, the variance changes as

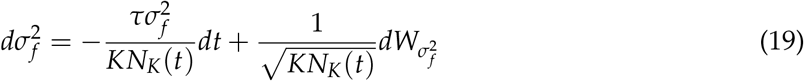

Taking expectations, the second term on the RHS vanishes, and we see that the expected variance in the population obeys

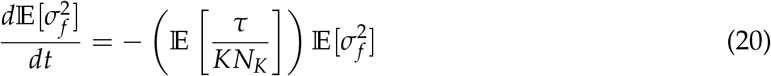

where we have decomposed the expectation of the product on the RHS into a product of expectations, which is admissible since the label *f* is completely arbitrary and thus independent of both 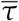 and *N*_*K*_(*t*). Eq. 20 is a simple linear ODE and has the solution

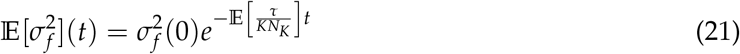

This equation tells us that the expected diversity (variance) of labels in the population decreases exponentially over time. The rate of loss of diversity is 𝔼[*τ*(*KN*_*K*_)^−1^], and thus, populations with a higher turnover rate *τ* and/or a lower population size *KN*_*K*_ lose diversity faster. This is because populations with higher *τ* experience more stochastic events per unit time and are thus more prone to stochastic extinction, while extinction is ‘easier’ in smaller populations because a smaller number of deaths is sufficient to eliminate a label from the population completely. Note that *which* labels/individuals are lost is entirely random (since all labels are arbitrary), but nevertheless, labels tend to be stochastically lost until only a single label remains in the population. Upon rescaling time as *t* → *τt*, equation 21 recovers the continuous time version of the loss of heterozygosity formula for finite populations from population genetics (Ewens, 2004, Eq. 1.5; Crow and Kimura, 1970, sections 7.3 and 8.4).

##### Box 5

**An example: Noise-induced biasing with two competing types**

Consider a population comprised of two competing types of individuals (denoted 1 and 2). For pedagogical clarity, we assume that the birth and death rates of type 1 are simply shifted from those of type 2 by constants *ϵ*_*b*_ and *ϵ*_*d*_ respectively, *i.e*..

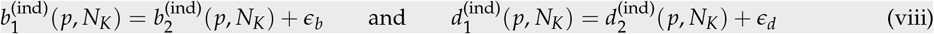

We provide potential biological interpretations of this model in terms of either ecological ‘rate modula-tors’ (Fronhofer et al., 2023) or competing pathogen strains (Parsons et al., 2018) in Supplementary section S10. Using the definitions of the selection coefficient (*s*) and noise-induced selection coefficient (*κ*), we find

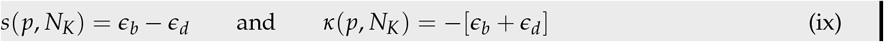

Equation ix shows that all else being equal, reducing the death rate leads to a more favourable evolutionary outcome than increasing the birth rate by the same amount (also see McLeod and Day, 2019a and Raatz and Traulsen, 2023). We now explain the subtle ways in which demographic stochasticity biases evolutionary dynamics.

###### Noise-induced biasing in the absence of natural selection

Let us assume that *ϵ*_*b*_ = *ϵ*_*d*_ = *ϵ*. This corresponds to a faster pace-of-life in type 1 relative to type 2. From Eq. (ix), we see that *s*(*p, N*_*K*_) = 0, and thus the two types have equal fitness. In the absence of natural selection, a given initial frequency remains unchanged over time in infinitely large populations. In finite populations experiencing only neutral genetic drift, we expect the probability of fixation of a type to be proportional to its initial frequency. The effects of noise-induced biasing through the direct mechanism (noise-induced selection) can be observed by looking at the change in the expected frequency 𝔼[*p*], which from Eq. 9 follows:

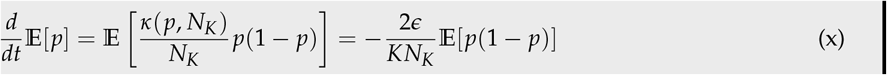

Since the RHS of Eq. (x) is always negative for *p* ∈ (0, 1), we can infer that the proportion of type 1 individuals is expected to decrease to 0 from any initial frequency. Note that unlike in drift, it is always type 2 that is expected to fix. This result is a manifestation of noise-induced selection: All else being equal, a faster pace of life comes with a greater variance in the change of population density within a given time interval since there are simply more stochastic birth/death events taking place, and types with a slower pace of life (type 2) are thus favoured (Parsons and Quince, 2007; Parsons et al., 2010; Wodarz et al., 2017).

To illustrate the indirect mechanism of noise-induced biasing, we need to assume a functional form for the turnover rates *τ*_*i*_. In Supplementary section S10, we obtain an exact expression for the speed density when *τ*_1_ = *bp* + *c* and *τ*_2_ = *bp* + *c* − 2*ϵ* for suitable constants *b* and *c. c* can be viewed as an ‘intrinsic’ turnover rate, and *b* as a frequency-dependent component that may be either positive or negative. Figure 3 plots the speed density for various parameter values, illustrating both the direct (Fig 3A) and indirect (Fig 3B) mechanisms of noise-induced biasing. Note that the direct and indirect mechanisms may operate in isolation or simultaneously and may either supplement (red curve in Fig. 3A and green curve in 3B) or oppose (red curves in 3A and 3B) each other.

**Figure 3:**
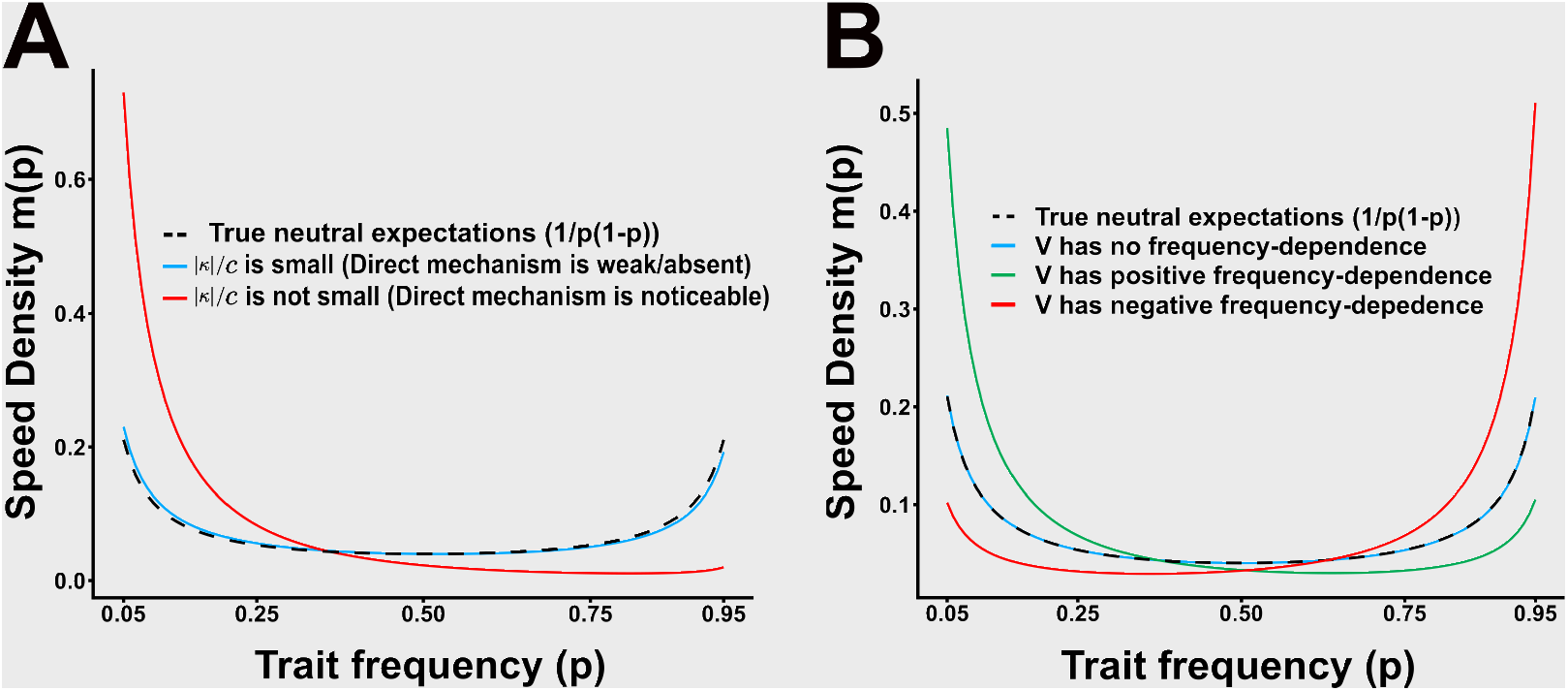
Two distinct noise-induced effects that bias trait distributions. **A**. If the magnitude of the noise-induced selection coefficient *κ* = − 2*ϵ* is large relative to the intrinsic turnover rate *c*, the direct mechanism of noise-induced biasing operates. Parameters are chosen such that *V*(*p*) = *τ*_1_ + *pκ* = (*b* − 2*ϵ*)*p* + *c* is not frequency-dependent (blue: *ϵ* = 0.5, *b* = 1, *c* = 10; red: *ϵ* = 0.5, *b* = 1, *c* = 0.5). **B**. The speed density can also be biased if *V*(*p*) is frequency-dependent. This indirect mechanism of noise-induced biasing favours the type that reduces *V*(*p*). Parameters in this panel are chosen such that |*κ*| is small relative to *c* and thus the strength of the fast mechanism is negligible (blue: *ϵ* = 0.025, *b* = 0.05, *c* = 10; green: *ϵ* = 0.025, *b* = 50, *c* = 10; red: *ϵ* = 0.025, *b* = −8.5, *c* = 10)

###### Noise-induced biasing in the presence of natural selection

Assume now that *ϵ*_*b*_ > *ϵ*_*d*_ > 0. In this case, *s* > 0 and thus natural selection favors type 1 individuals. As before, there are two ways in which demographic stochasticity can bias evolutionary dynamics towards certain types. Noise-induced selection could drive the expected trajectory towards fixation of type 2 despite type 1 being favored by natural selection. In supplementary section S10, we show that this can happen if and only if

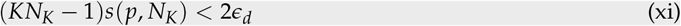

Thus, noise-induced selection can reverse the predictions of natural selection when *s* · (*KN*_*K*_ − 1) is sufficiently small, *i.e*. when *natural selection is weak* (*s* is small), *populations are small* (*KN*_*K*_ is small), or both. Since the strength and direction of the indirect mechanism depend on the functional form of *τ*_*i*_, we do not explicitly study it here. However, we provide some preliminary observations in Supplementary section S10.

## Discussion

The central result of our paper is a set of stochastic dynamical equations for changes in trait frequencies in the population (Eq. 5) that generalizes the replicator equation (or, with mutations, the replicator-mutator equation - Eq. S29) to finite populations of non-constant size evolving in continuous time. From this, we derive a generalization of the Price equation (Eq. 12) and Fisher’s fundamental theorem (Eq. 14) to such populations, as well as an equation for changes in population variance of a type-level quantity (Eq. 17). Our framework recovers well-known equations of population biology, such as the replicator-mutator equation, Price equation, and Fisher’s fundamental theorem, in the infinite population limit (see section S7). Our equations also reveal that demographic stochasticity alone can cause certain types to be more likely to be observed in a population, an effect we term noise-induced biasing. Noise-induced biasing can manifest through two distinct mechanisms (Box 4), one that directly affects the selection term in the replicator equation and another that acts indirectly by affecting the time spent at various states. Several theorists have called for a reformulation of eco-evolutionary dynamics starting from stochastic birth-death processes on the grounds that such a formulation is more fundamental and mechanistic (Metcalf and Pavard, 2007; Lambert, 2010; Doebeli et al., 2017). Our equations provide a starting point for such a reformulation by deriving some fundamental equations for the eco-evolutionary dynamics of finite, stochastically fluctuating populations.

### Finite population effects on eco-evolutionary dynamics

Our equations show that noise-induced effects can bias evolutionary outcomes through two major, qualitatively different mechanisms (Box 4). The direct mechanism manifests in the infinitesimal mean of our SDEs via a ‘noise-induced’ term that is inversely proportional to the population size (the second term on the RHS of Eqs. 5, 12, and 17). The direct mechanism has previously been reported in various specific contexts (Parsons and Quince, 2007; Parsons et al., 2010; Wodarz et al., 2017; Parsons et al., 2018; Kuosmanen et al., 2022). Since the terms capturing these effects in Eqs. 5, 12, and 17 have the same mathematical form as the effect of classic natural selection, the direct mechanism has previously been referred to as noise-induced selection (Week et al., 2021), and we believe this term is appropriate. It has also been the object of study in early models of bet-hedging in finite populations (Gillespie, 1974; Gillespie, 1977; Shpak, 2005), thus explaining why noise-induced selection has previously been associated with the ‘Gillespie effect’ for reduced variance (Parsons and Quince, 2007; Parsons et al., 2010; Parsons et al., 2018). However, note that the variance that is studied in bet-hedging models is typically variance in *offspring numbers* (Gillespie, 1977). The variance in Eq. 4b is *not* variance in offspring numbers, but instead variance in the ecological ‘growth rate’ *dx*_*i*_ (over an infinitesimal time interval), a quantity that has sometimes been called ‘demographic variance’ (Engen et al., 1998; Shpak, 2007). Furthermore, unlike many classic bet-hedging papers such as Gillespie (1974) and Frank and Slatkin (1990), in our framework both *w*_*i*_ and *τ*_*i*_ (and thus the mean and variance of the change in population density) are defined from first principles in terms of birth and death rates (Eqs 1 and 2)

In contrast, the indirect mechanism acts through the infinitesimal variance of our SDEs and thus does not appear in the expected change in trait frequency. This mechanism is a subtle effect of frequency-dependent demographic stochasticity, can be present even when the direct mechanism (i.e. noise-induced selection) is weak or absent (Fig 3B), and is revealed as a systematic bias in the speed density that causes the system to spend disproportionately more time in certain states (McLeod and Day, 2019a; McLeod and Day, 2019b; see Box 4). If a stationary distribution exists, the indirect mechanism will be visible in the stationary distribution but not in the expected trajectory. The indirect mechanism can be thought of as analogous to frequency-dependent ‘viscosity’; populations tend to accumulate in those configurations that lead to slower changes in population composition, and we are thus more likely to observe the population in those configurations that make the rate of change of the population ‘slower’. This observation has previously been referred to as the effect of ‘evolutionary noise’ on evolutionary dynamics (McLeod and Day, 2019a; McLeod and Day, 2019b).

Our results suggest an intriguing requirement for neutral evolution in finite populations. In models of genetic drift, evolution is said to be neutral if the fixation probability of a type in a population is proportional to the initial frequency alone (Ewens, 2004). For populations of non-constant size, we see that neutrality in this sense is not ensured if the trait in question is neutral with respect to fitness *w* alone. Instead, neutral evolution also requires all trait variants to have equal turnover rates, failing which evolution will be ‘quasi-neutral’ and favour those types associated with lower turnover rates (Parsons and Quince, 2007; Parsons et al., 2010; Kuosmanen et al., 2022). In other words, even in (finite) populations with no differential fitness among traits, there exists a directional evolutionary force that may systematically bias the course of evolution. Furthermore, the indirect mechanism of noise-induced biasing means that we may be more likely to observe the population in states in which certain types are overrepresented due to a biasing of the speed density and, when it exists, the stationary distribution (McLeod and Day, 2019a; McLeod and Day, 2019b). Thus, even if all individuals in the population have equal fitness and equal turnover, types associated with lower *V*(*p*) are *still* ‘favoured’ in the sense that we are more likely to observe the population in a configuration at which these types are overrepresented (McLeod and Day, 2019a; McLeod and Day, 2019b) relative to neutral expectations as defined above. However, it may be noted that the strength of noise-induced biasing is likely to be small or even negligible unless the population size is sufficiently small and/or close to having equal mean fitness.

In our model, noise-induced selection is particular to fluctuating populations and does not occur in models with fixed population sizes such as the Wright-Fisher or Moran models (Box 3). Taken alongside other theoretical (Lambert, 2010; Parsons et al., 2010; Abu Awad and Coron, 2018; Kuosmanen et al., 2022; Mazzolini and Grilli, 2023) and empirical (Papkou et al., 2016; Chavhan et al., 2019) studies on evolution in fluctuating populations, this last point suggests that models which assume fixed total population size, such as Wright-Fisher and Moran, may miss out on important evolutionary phenomena that are only seen in finite populations of *non-constant size*. We explain how our framework incorporates the ‘drift-induced selection’ from sex chromosome evolution (Veller et al., 2017; Saunders et al., 2018) as well as some previous studies from social evolution (McLeod and Day, 2019a) and epidemiology (Parsons et al., 2018; Day et al., 2020) in Supplementary section S8. We also explain connections with some other general frameworks of eco-evolutionary dynamics (Rice, 2020; Week et al., 2021; Kuosmanen et al., 2022) in Supplementary section S9.

### Concluding remarks

A steadily growing body of literature has begun to highlight the surprising and counter-intuitive effects of demographic stochasticity in shaping evolutionary outcomes in many ecological scenarios. In this paper, we derive from first principles stochastic dynamical equations for eco-evolutionary dynamics that generalize some standard equations of population biology, thus providing a conceptual synthesis of the findings of these previous studies. The terms of the equations we derive lend themselves to simple biological interpretations and recover standard equations of evolutionary theory in the infinite population limit. To the best of our knowledge, the equations we derive in this paper are the first to showcase how demographic stochasticity generically alters some standard equations of population biology. The utility of the equations we derive thus lies not (necessarily) in their solutions for specific models but instead in their generality and the fact that their terms help us clearly think about the various evolutionary phenomena operating in biological populations (Queller, 2017; Lehtonen, 2018; Lion, 2018; Luque and Baravalle, 2021). The direct and indirect mechanisms of noise-induced biasing have distinct origins, may operate either independently or together, and may push evolution in different directions (see the example in Box 5). It is, therefore, essential that studies explicitly differentiate between these two mechanisms to identify which noise-induced effects are germane to any particular biological population (Box 4). By re-deriving some standard equations of population dynamics for finite populations, we provide a framework with which to approach particular finite population systems and systematically determine which evolutionary forces are important from demographic first principles.

Populations in stochastically fluctuating environments are affected by interactions between two qualitatively different forms of noise — *environmental stochasticity* from fluctuations in environmental factors such as temperature and precipitation, and *demographic stochasticity* due to stochasticity in birth and death rates in finite populations (Lande, 1993; Shoemaker et al., 2020). Though we neglect environmental stochasticity in our current work, populations that experience both environmental and demographic stochasticity often exhibit surprising and counter-intuitive eco-evolutionary dynamics (Gokhale and Hauert, 2016; Chavhan et al., 2021). Studying the interplay of noise-induced biasing with environmental stochasticity may thus present a promising avenue for future work. Since both the strength (Hamilton, 1966; Mallet et al., 2011; Lehtonen, 2020b) and the direction (Chapman et al., 2003; Maklakov and Chapman, 2019) of natural selection may vary in populations structured by classes such as age or sex, extending our model to include population structure could also be fruitful. On the empirical side, developing methods to disentangle different demographic stochastic effects from empirical datasets could be another interesting avenue for future work.

## Acknowledgements

We are grateful to Srikanth Iyer, Kavita Jain, Sébastien Lion, and two anonymous reviewers for their helpful comments on an earlier draft of this manuscript. ASB is supported by a Kishore Yaigyanik Protsahan Yojana (KVPY) fellowship from the Department of Science and Technology of the Government of India (Fellowship ID: SX-1711025).

## Author Contributions

**Ananda Shikhara Bhat:** Conceptualization, Methodology, Formal Analysis, Investigation, Writing - Original Draft, Writing - Review & Editing, Visualization; **Vishwesha Guttal:** Conceptualization, Methodology, Validation, Writing - Review & Editing, Supervision.

## Data and Code Availability

Code for running the simulations and reproducing the figures presented in this manuscript has been supplied as a ZIP file. All code will be made publicly available in a GitHub repository upon acceptance of the manuscript.

## Supplementary Information

### S1 The master equation and the system size expansion

Given a system with *m* different types of individuals and birth and death rate functions *b*_*i*_(**n**) and *d*_*i*_(**n**), we are interested in finding an equation for the rate of change of the conditional probability *P*(**n**, *t*|**n**_0_, 0), the probability of finding the population in a state **n** at time *t*. Henceforth, we omit the conditioning for notational brevity and simply write *P*(**n**, *t*) for this quantity. We assume that the birth and death rates are of the order of the total population size, *i.e*. that *b*_*i*_(**n**) and *d*_*i*_(**n**) are 𝒪(∑_*i*_ *n*_*i*_) functions.

For each *i* ∈ {1, …, *m*}, let us now define two step operators 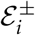 by their action on any function *f* ([*n*_1_, …, *n*_*m*_], *t*) as:

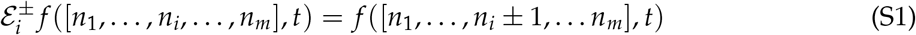

In other words, 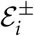 just changes the population through the addition or removal of one type *i* individual. We can now write down an exact equation for the rate of change of *P*(**n**, *t*) by noting that the only direct transitions allowed are those from populations that are exactly one individual away from our focal population. Thus, we have the relation

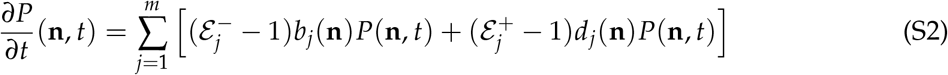

This equation is called the ‘master equation’, and completely characterizes our *m*-dimensional process.

#### S1.1 Scaling assumptions, functional forms of birth and death rates

As mentioned in the main text, we assume that there is a carrying capacity/population size measure *K* > 0 such that the total population size ∑_*i*_ *n*_*i*_ is expected to be 𝒪(*K*). This allows us to move from population numbers **n** to population ‘densities’ **x** = **n**/*K*. Specifically, we assume that we can find 𝒪(1) functions 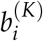 and 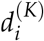 such that we can write

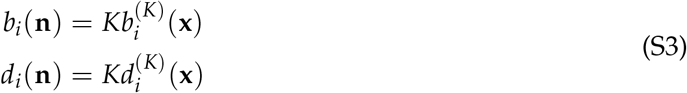

Note that this assumption means that 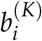 and 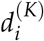 remain well-defined even in the *K* → ∞ limit, since *b*_*i*_/*K* and *d*_*i*_/*K* remain 𝒪(1) by our assumption on the scaling properties of **n**, *b*_*i*_, and *d*_*i*_. Thus, we may still speak of population densities *x* in the infinite population size limit (*K* → ∞). Note that this scaling assumption implies that in the functional forms S4, we assume that 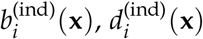, and *Q*_*i*_(**x**) are all 𝒪(1) functions.

In the supplementary, we assume birth and death rates have the slightly more general functional form

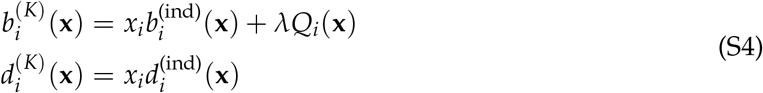

where, as defined in the main text, 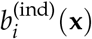 and 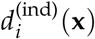 are non-negative functions that describe the per-capita birth and death rate of type *i* individuals, respectively. When there are no type *i* individuals in the population (*x*_*i*_ = 0), individuals of type *i* may still be born through mutations of other types or via immigration from other sources (gene flow). Thus, the birth rate of type *i* may contain terms that do not depend multiplicatively on the density *x*_*i*_ of type *i* individuals. The term *λQ*_*i*_(**x**) accounts for this possibility. The equations in the main text are recovered upon removing this term by setting *λ* = 0.

The term *λQ*_*i*_ in Eq. S4 thus models an influx of type *i* individuals from sources other than the existing pool of type *i* individuals. Here, *λ* ≥ 0 is a constant describing the rate of influx of type *i* individuals from sources other than the existing pool of type *i* individuals, and *Q*_*i*_(**x**) is a non-negative function that describes this additional contribution. For example, if type *i* individuals can arise due to mutations of offspring of other types of individuals during birth, *λ* would represent a mutation rate (typically denoted by *µ*) and *Q*_*i*_ would model the functional form of mutation. A common choice, for example, is *Q*_*i*_(**x**) = ∑_*j*≠ *i*_ *x*_*j*_ (*i.e*. the mutation *j* → *i* occurs at a total rate of *µx*_*j*_). The influx term could also model immigration of type *i* individuals from other populations since such immigration would depend not on the density of individuals *x*_*i*_ in our focal population, but on the density of individuals in the ‘source population’ from which individuals are emigrating into our focal population. In this latter case, *λ* would represent a dispersal rate and *Q*_*i*_ would model the dispersal. Note that no analogous problem exists for the death rate, since the death rate of type *i* individuals must be 0 when *x*_*i*_ is 0 to ensure that we never have negative population densities.

#### S1.2 The infinite population limit

We can now more clearly speak about what we mean by the infinite population limit. Recall that *K* is a population size measure (Czuppon and Traulsen, 2021). Since the growth rate is expected to be negative when ∑_*i*_ *n*_*i*_ > *K*, if a population is initiated with ∑_*i*_ *n*_*i*_(0) being 𝒪(*K*), the total population size ∑_*i*_ *n*_*i*_(*t*) does not have unbounded growth, but instead remains 𝒪(*K*) before the population eventually goes extinct. Thus, in terms of population densities **x**(*t*), the scaled population size ∑_*i*_ *x*_*i*_(*t*) remains 𝒪(1). When we speak of the ‘infinite population limit’ *K* → ∞, we thus take the limit of *K* → ∞ along with initial population size ∑_*i*_ *n*_*i*_(0) → ∞ such that the initial total scaled population size ∑_*i*_ *x*_*i*_(0) remains 𝒪(1). Thus, the limit in question is the standard domain of deterministic population models, namely, a model of a population with infinite population size but finite population density. Note that in the infinite population limit, we cannot speak of **n** or *K* individually but only of the ratio **x** = **n**/*K*, which is guaranteed to remain well-defined by our scaling assumptions. A more thorough, rigorous treatment of the mathematical details behind the rescaling procedure and infinite population limit is presented in Champagnat et al., 2006.

#### S1.3 Step operators and system size expansion

To describe our stochastic process in terms of population densities rather than absolute population sizes, we now define new step operators 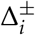 by their action on any real-valued function *f* (**x**, *t*) as

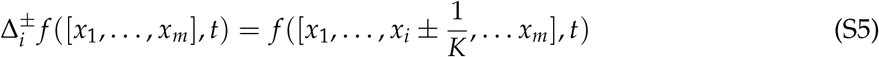

In terms of these new variables, (S2) becomes

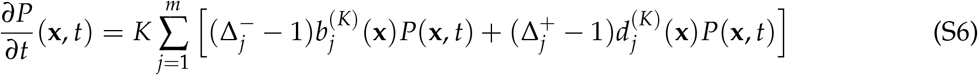

If *K* is large, we can now Taylor expand the action of the step operators as

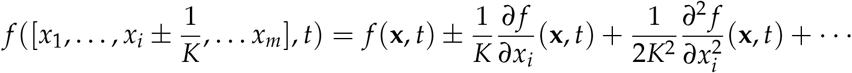

which, after substituting into (S6) and neglecting higher order terms, yields the equation

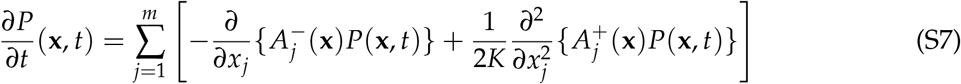

where

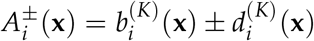

Equation (S7) is an *m*-dimensional version of a ‘Fokker-Planck equation’ or ‘diffusion equation’ for the probability density *P*(*x, t*). For a more detailed discussion on ‘system size approximations’ such as the one we carried out above, we refer the reader to Chapter 11 of Ethier and Kurtz, 1986 for the mathematically rigorous theory and Chapter 10 of Van Kampen, 1981 for a heuristic approach. Pedagogical treatments focused on eco-evolutionary population dynamics can be found in Black and McKane, 2012 and Czuppon and Traulsen, 2021.

##### Itô SDE representation

For our purposes, we will often find it convenient to describe the same process as defined by the Fokker-Planck equation (S7) via an ‘Itô stochastic differential equation’. It is well-known (Øksendal, 1998) that a stochastic process whose probability density function satisfies a Fokker-Planck equation of the form (S7) is equivalent to an *m*-dimensional stochastic process obtained as the solution to the Itô SDE

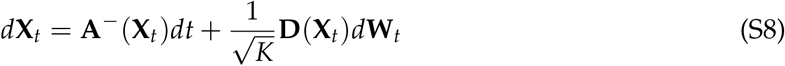

Here, **A**^−^(**X**_*t*_) is an *m*-dimensional vector with *i*^th^ element 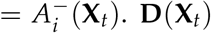. **D**(**X**_*t*_) is an *m* × *m* matrix with *ij*th element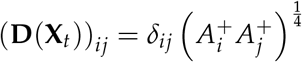, where *δ*_*ij*_ is the Kronecker delta symbol, defined by

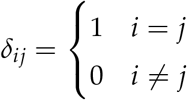

Finally, **W**_*t*_ is the *m*-dimensional Wiener process (standard Brownian motion) and can be thought of as a vector of independent one-dimensional Wiener processes.

### S2 Trait frequency dynamics using Itô’s formula

We first recall the version of the multi-dimensional Itô’s formula that will be relevant to us.

Consider an *m*-dimensional real Itô process **X**_*t*_ given by the solution to

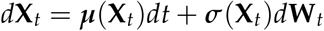

where ***µ*** : ℝ^*m*^ → ℝ^*m*^ is the ‘drift vector’ and ***σ*** : ℝ^*m*^ → ℝ^*m*×*m*^ is the ‘diffusion matrix’. Let *f* : ℝ^*m*^ → ℝ be an arbitrary *C*^2^(ℝ^*m*^) function. Then, Itô’s formula (Øksendal, 1998, Section 4.2) states that the stochastic process *f* (**X**_*t*_) must satisfy:

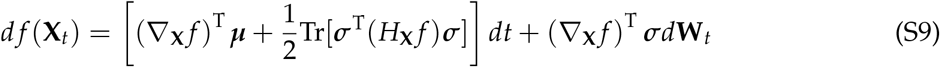

where Tr[·] denotes the trace of a matrix, (·)^T^ denotes the transpose, and we have suppressed the **X**_*t*_ dependence of ***µ*** and ***σ*** to reduce clutter. Here, ∇_**x**_ *f* is the *m*-dimensional *gradient vector* of *f* with respect to **x** and *H*_**x**_ *f* is the *m* × *m Hessian matrix* of *f* with respect to **x**, respectively defined for *f* ([*x*_1_, …, *x*_*m*_]^T^) as:

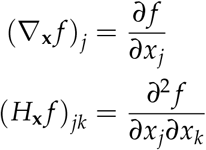

In our case, we have the Itô process given by (S8), which defines how the density of each type of individual changes over time. We thus have ***µ***(**X**_*t*_) = **A**^−^(**X**_*t*_) and 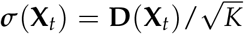. For each fixed *i* ∈ {1, 2, …, *m*}, let us define a scalar function *f*_*i*_ : ℝ^*m*^ → ℝ as

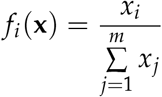

Thus, *f*_*i*_(**X**_*t*_) gives us the frequency of type *i* individuals when the population is described by the vector **X**_*t*_. As an aside, note that we only need to calculate frequencies for *i* ∈ {1, 2, …, *m* − 1} along with *N*_*K*_ = ∑ *x*_*i*_. We can now use Itô’s formula (S9) to describe how *f*_*i*_ changes over time. The *j*^th^ element of the gradient of *f*_*i*_ is given by:

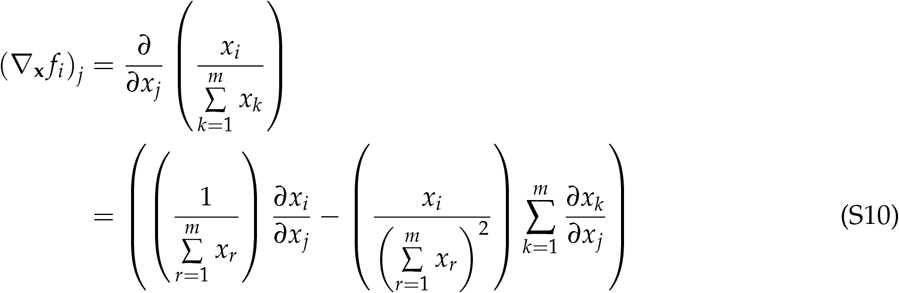

where we have defined the frequency of the *i*^th^ type *p*_*i*_ = *f*_*i*_(**x**). To proceed further, we require the quantity 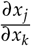 for any pair of types *j, k* ∈ {1, 2, 3, …, *m* − 1, *m*}. Since changes in densities in our system are only being determined by ecological interactions at the individual level, with changes in total population size being an emergent quantity, we can assume that our system obeys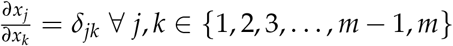. Note that this is *not* true if the total population size is held constant since changes in densities of one type must be accompanied by complementary changes in densities of at least one other type to keep the total density ∑_*i*_ *x*_*i*_ strictly constant.

We can now substitute 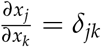 into equation (S10). Upon doing this, we obtain

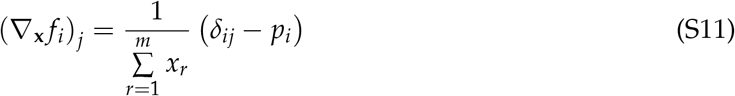

Similarly, we can also calculate the Hessian. The *jk*^th^ element of the Hessian is given by:

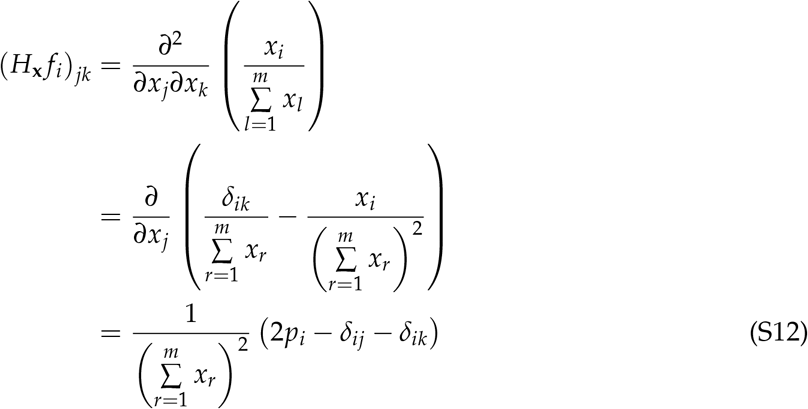

Thus, for the first term of (S9), we have:

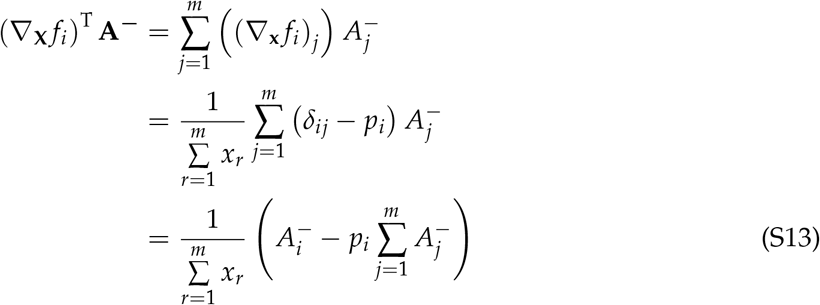

This term describes the effects of selection and influx (mutation/migration) at the infinite population limit. However, the finiteness of the population adds a second directional term to these dynamics, described by the second term that multiplies *dt* in (S9). To calculate it, we first calculate:

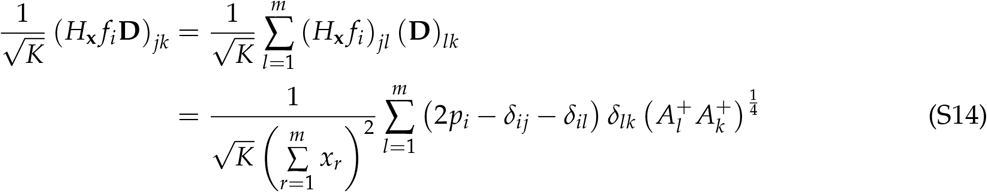

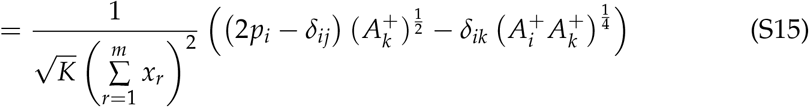

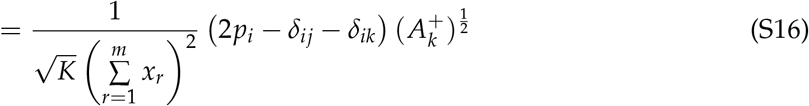

and thus:

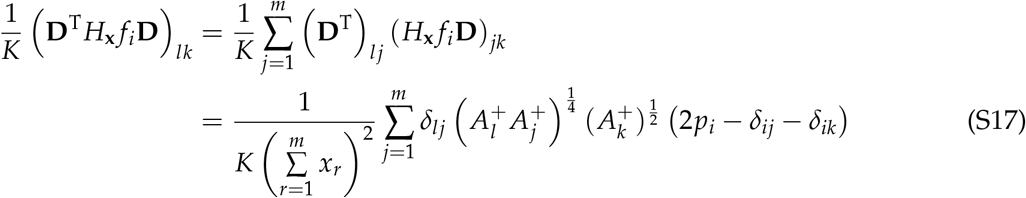

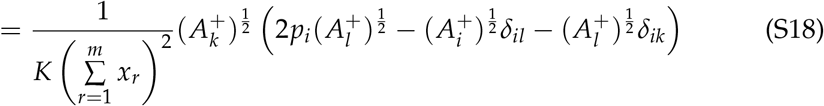

Using this, we see that the trace of this matrix is given by:

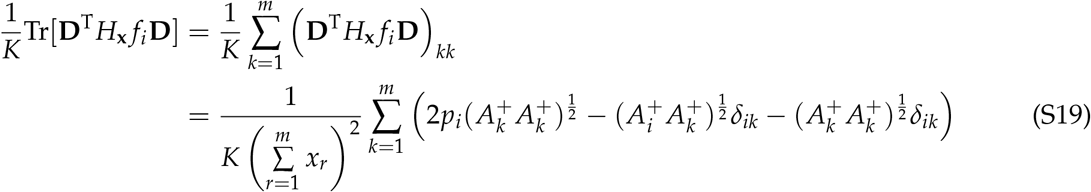

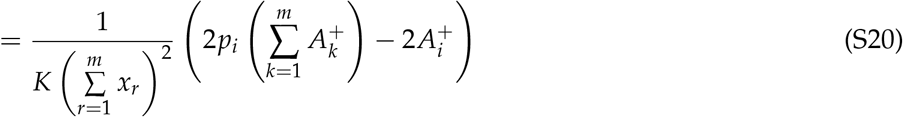

and thus, the second term multiplying *dt* in (S9) is given by:

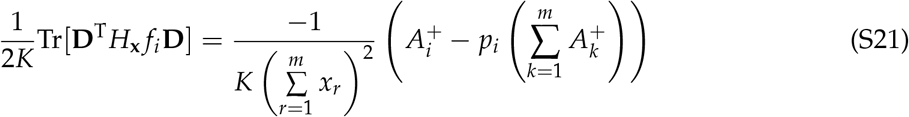

Finally, denoting 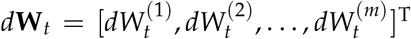 where each 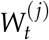 is an independent one dimensional Wiener process, we have:

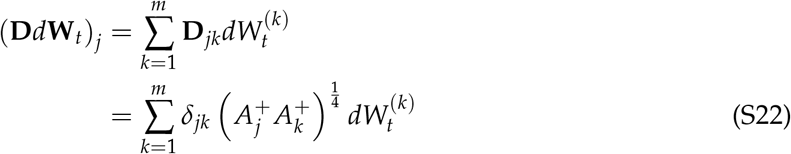

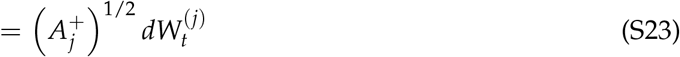

Thus, using (S11), we see that the last term on the RHS of (S9) is given by:

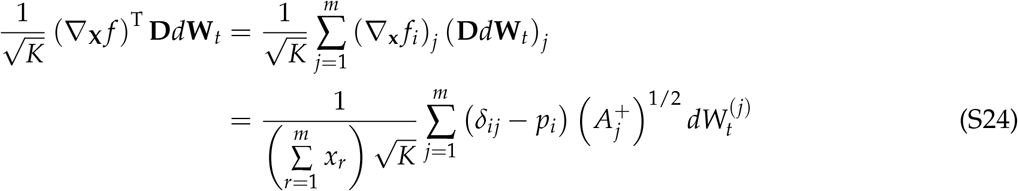

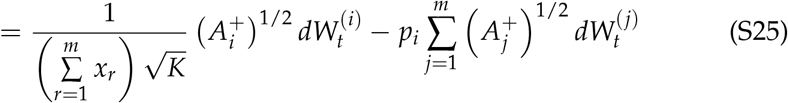

Putting equations (S13), (S21) and (S25) into (S9) and letting 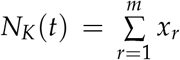 we see that *p*_*i*_ = *f*_*i*_(**X**)_*t*_, the frequency of the *i*^th^ type in the population **X**_*t*_, changes according to the equation:

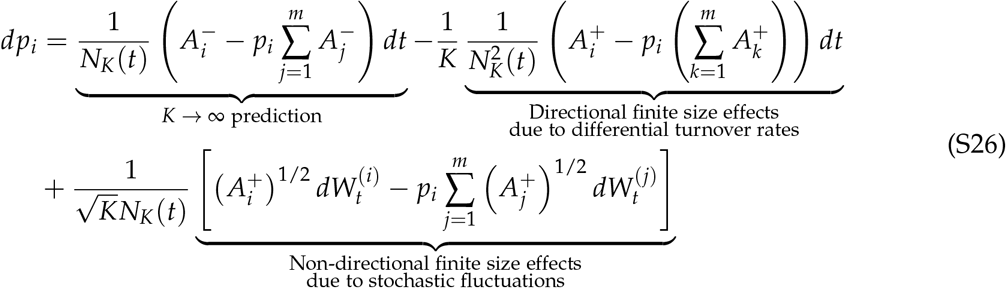

Plugging the functional forms of (S4) and the definitions of *w*_*i*_ and *τ*_*i*_ into the definitions of 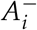 and 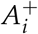, we obtain the relations

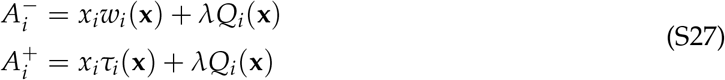

Thus, for the first term of (S26), we have

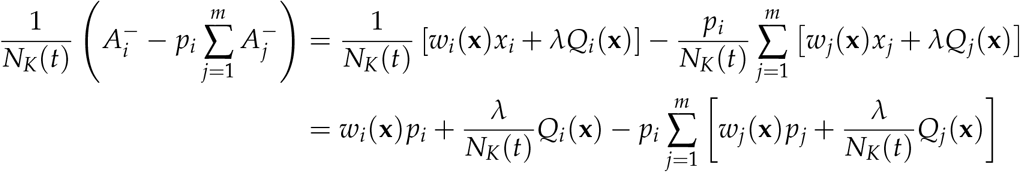

where we have used the definition of *p* _*i*_ from (iii). Now using the definition of mean fitness from (iv) and rearranging terms gives us

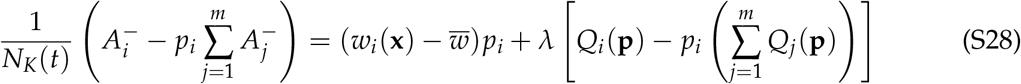

where we have defined *Q*_*j*_(**p**) = *Q*_*j*_(**x**)/*N*_*K*_(*t*). Repeating the exact same calculations for the 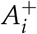 terms in the second term of (S26) now yields equation

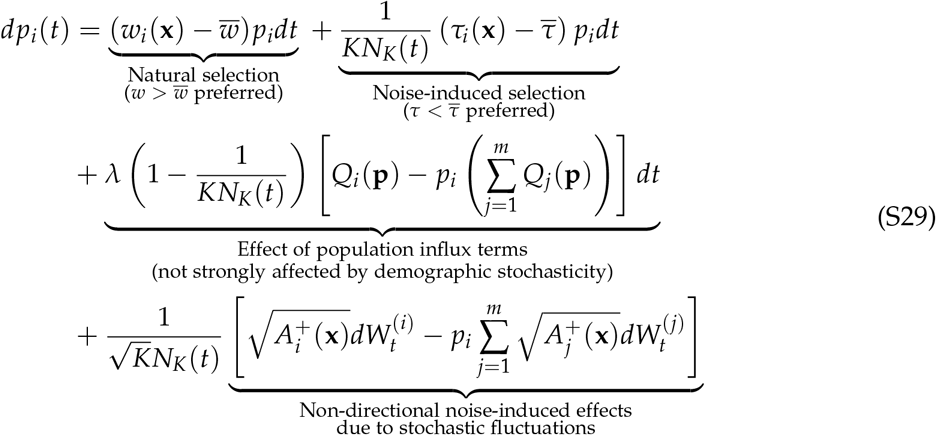

which is the first key result (5) presented in the main text upon setting *λ* = 0.

The third term on the RHS of Eq. S29 represents potential biasing effects due to the influx of individuals of type *i* in a manner that does not depend purely multiplicatively on the current population density *x*_*i*_ of type *i* individuals (for example, through immigration from an external population or mutation of other types during birth). Since 1 − 1/*KN*_*K*_ is typically very close to 1 for medium to large population size (*KN*_*K*_), we see that such influxes of individuals are not strongly affected by demographic stochasticity and thus have qualitatively similar effects in small, large, and infinite populations. This observation justifies our decision to neglect such terms in the main text for the sake of conceptual clarity, keeping the goals of a synthesis in mind.

### S3 A stochastic analog of the Price equation for finite, fluctuating populations

In this section, we will derive an SDE for the rate of change of the population mean value of any type-level quantity in finite, fluctuating populations. Let *f* be any type-level quantity, with value *f*_*i*_(*t*) for the *i*^th^ type. Using the product rule of calculus on the definition (iv) of the statistical mean tells us that we have the relation

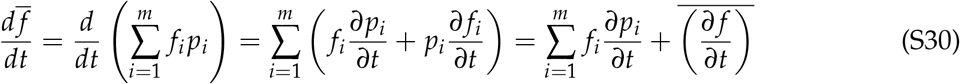

*i.e*.

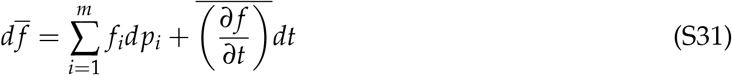

We will further simplify the first term on the RHS of (S31). We do this by using (S29), which gives us a representation of *dp*_*i*_. Using the RHS of (S29), we can conclude that we must have

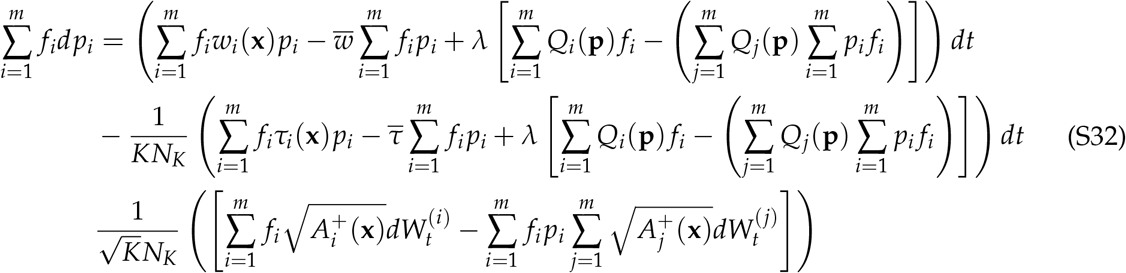

now using the definition of the statistical mean from (iv) in equation (S32), we obtain

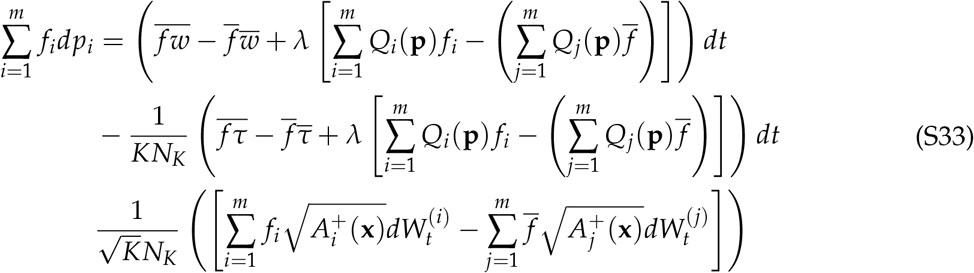

By the definition of the statistical covariance (v), we now obtain

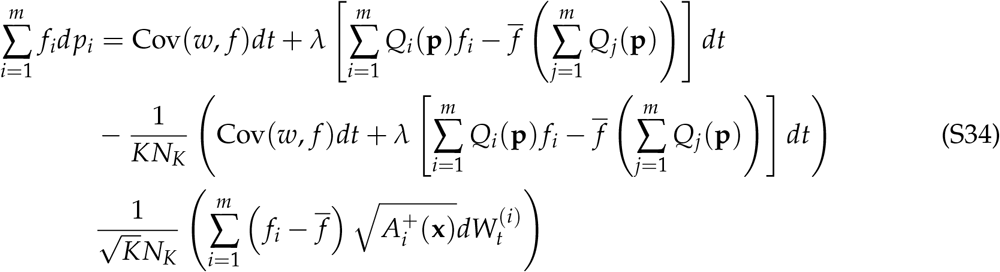

Collecting all terms that capture effects related to mutations/migrations (*i.e*. all terms with a *λ* factor) via defining the term

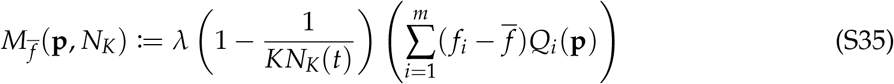

and collecting all stochastic integral terms via defining the term

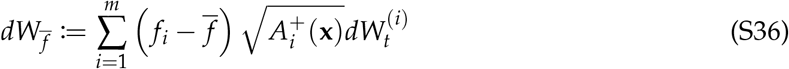

and substituting into equation (S34) now yields

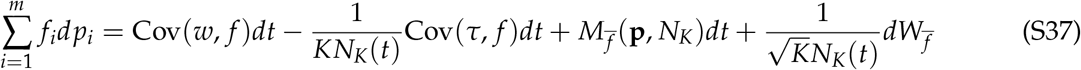

This is the simplified version of the first term on the RHS of equation (S31). Upon substitution, (S31) becomes

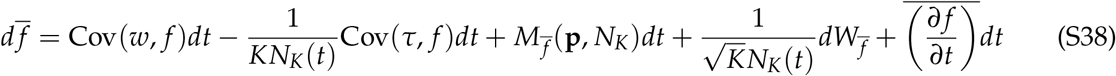

which is precisely equation (12) in the main text once we set *λ* = 0 (i.e. 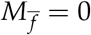).

### S4 A Price-like equation for the variance of a type-level quantity

In this section, we will derive an SDE for the rate of change of the variance of any type-level quantity in finite, fluctuating populations. From the definition (v), we see that the variance of any type level quantity *f* is given by:

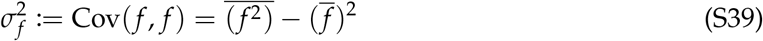

By the product rule, we have

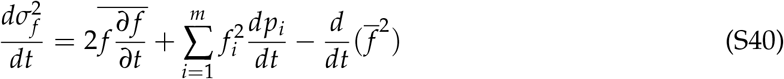

*i.e*.

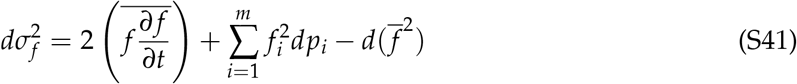

We will evaluate the RHS term by term. The first term is as simplified as can be without more information about *f*. For the second term, we can substitute *dp*_*i*_ from (S29) and then use the exact same steps we carried out in supplementary section S3 to derive equation (S38). Upon doing this, we obtain

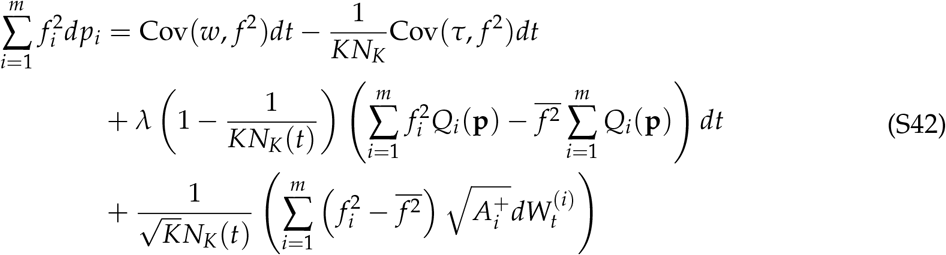

For the third term, we require Itô’s formula. Here, the relevant version of Itô’s formula is the one-dimensional version of (S9). Given a one-dimensional process 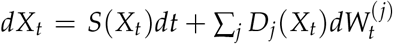 with *S, D*_*j*_ being suitable real functions and each 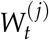 being an independent Wiener process, Itô’s formula says that given any *C*^2^(ℝ) function *g*(*x*), we have the relation:

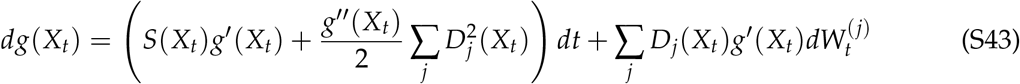

In our case, we have a one-dimensional process for the mean value 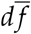 of the type level quantity, and the *C*^2^(ℝ) function *g*(*x*) = *x*^2^. Itô’s formula thus says that the third term of (S41) is given by:

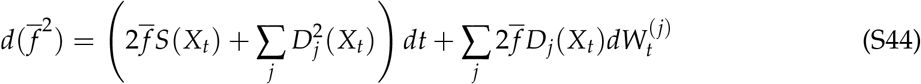

where the relevant functions *S* and *D*_*j*_ can be read off from (S38). Since the 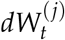 terms are unwieldy, we will denote the contribution of all the 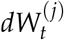 terms collectively by 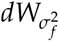 to reduce notational clutter and only explicitly calculate these terms at the end.

We can thus calculate

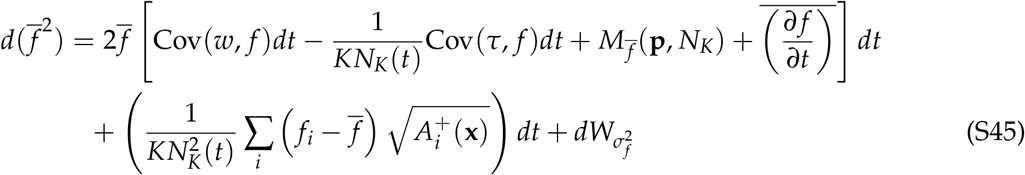

We now observe that the covariance operator is a bilinear form, *i.e*. given any three quantities *X, Y* and *Z* and any constant *a* ≠ 0, we have the relations:

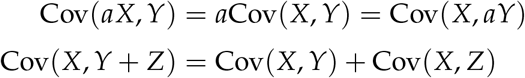

Substituting equations (S42) and (S45) into equation (S41) and using this property of covariances for the Cov(*w*, ·) and Cov(*τ*, ·) terms, we obtain:

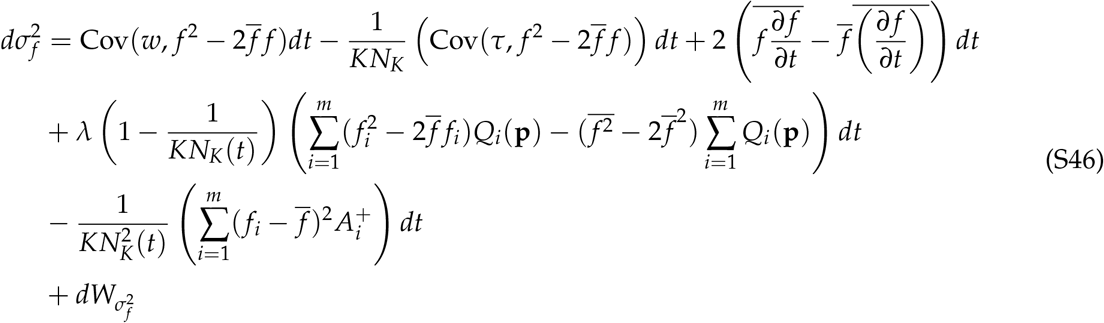

Now, we note that

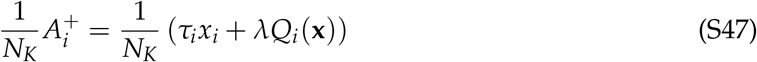

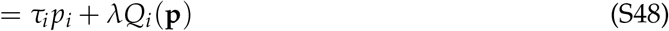

and thus the third line of (S46) is

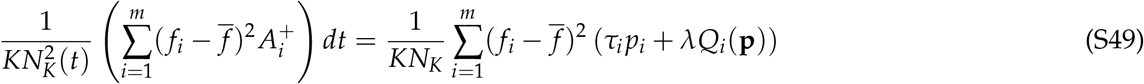

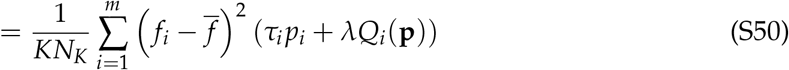

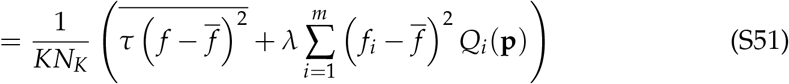

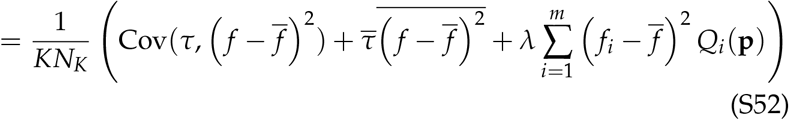

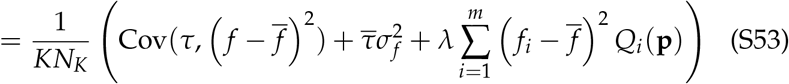

where we have used the definition of statistical covariance in the second to last line and used the definition of statistical variance in the last line. Substituting (S53) into (S46) and using 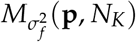 to denote the contributions of all the mutational terms (*i.e*. all terms with a *λ* factor) for notational brevity, we obtain

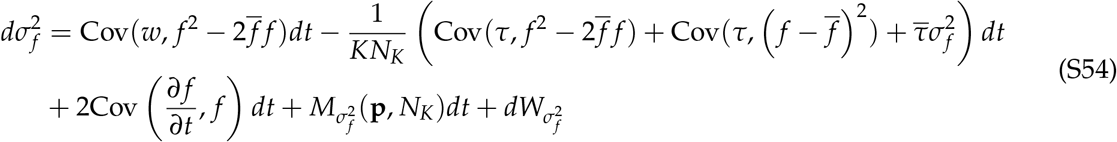

We can now complete the square inside the covariance terms of the first line of the RHS by writing 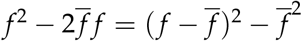 to obtain

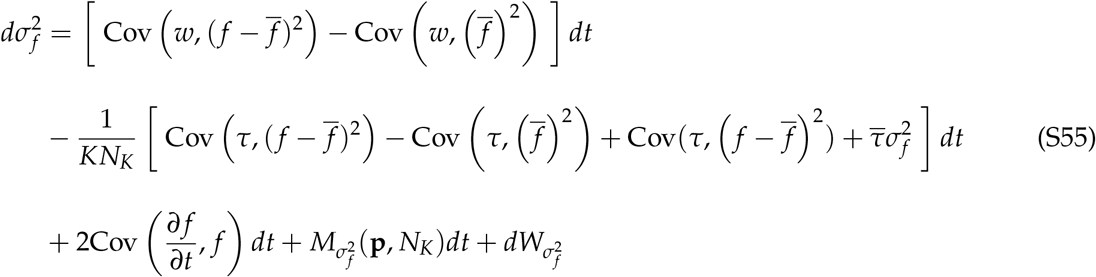

To simplify the covariance terms of the first line of the RHS, we observe that

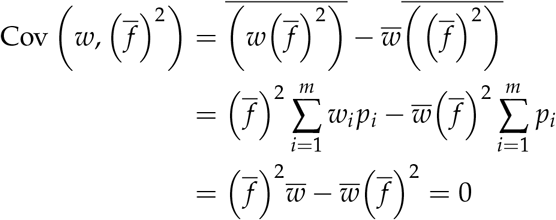

and similarly,

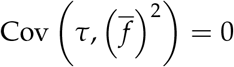

and thus, using this in (S55), we see that the rate of change of the variance of any type-level quantity *f* in the population satisfies:

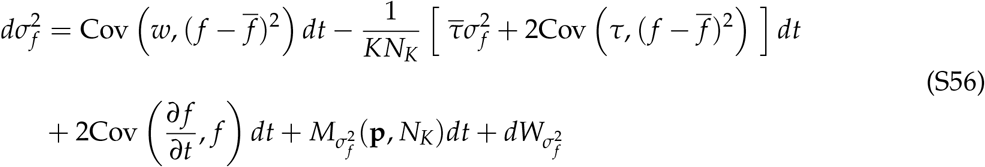

This is precisely equation (17) in the main text. To calculate the mutation term, we substitute (S53) into (S46) to find

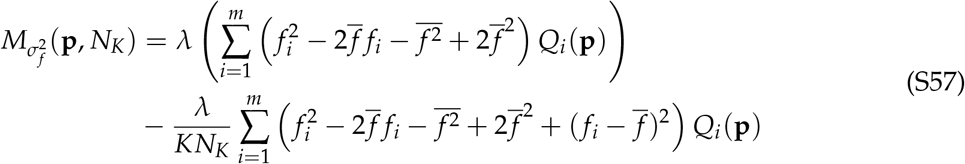

We can further simplify the first term of the RHS as

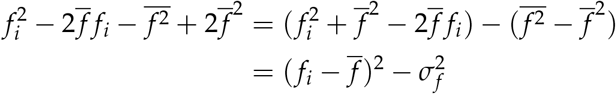

and similarly, the second term as

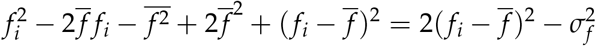

thus, the contributions of influx terms to the change in the variance of *f* are given by

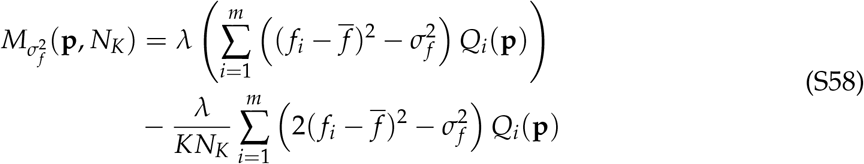

which after slight rearrangement becomes

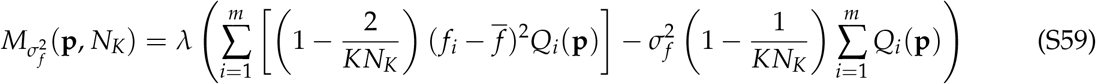

Finally, for the stochastic integral term, we can use equations (S42) and (S44) to calculate:

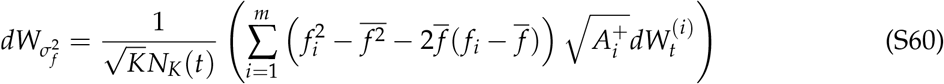

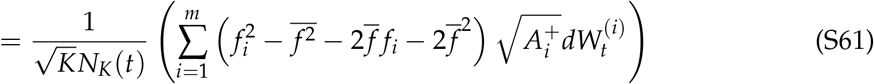

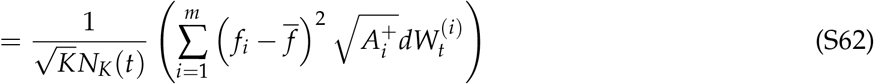

which is equation (18) in the main text upon setting *λ* = 0 (*i.e*.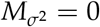).

### S5 A more elegant representation of sums of stochastic integrals against independent Wiener processes

We have arrived at three stochastic differential equations (equations (S29), (S38), and (S60)) that describe the change in the frequency of a type, the population mean value of a type-level quantity, and the population variance of a type-level quantity over time. All three of these equations contain sums of stochastic integrals of several independent functions against independent Wiener processes. In this section, we will present a more elegant representation of these terms as a single stochastic integral.

Let us first recall that given *m* independent one-dimensional Wiener processes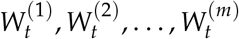, *m* ‘nice’ real functions *g*_1_(*x*), *g*_2_(*x*), …, *g*_*m*_(*x*), and the stochastic process

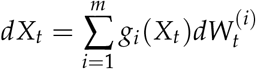

We can always find a *single* one-dimensional Wiener process *W*_*t*_ such that

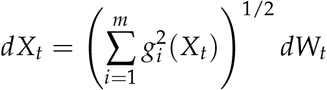

This result is well-known but we were unable to find a reference that explicitly proved it, and so we prove it as a lemma at the end of this supplementary section.

Using this result, we can now calculate the stochastic integral terms of our equations. For equation (7), we can calculate

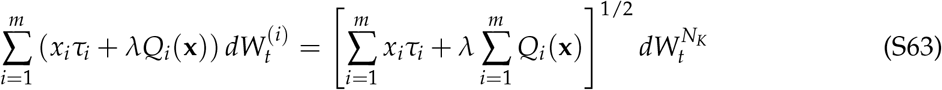

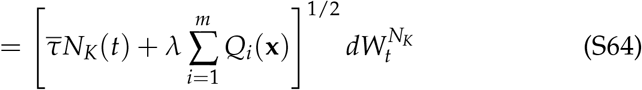

where 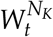 is a one-dimensional Wiener process. For equation (S29), the stochastic analog of the replicator-mutator equation, we find that the noise term can be written as a stochastic integral against a single Wiener process *W*_*t*_ as

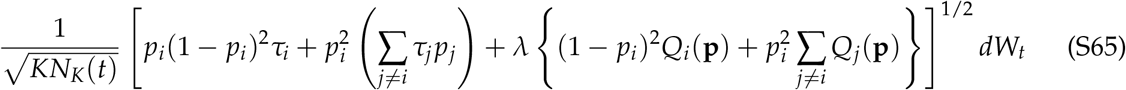

Note that the inclusion of influx terms (*λ* ≠ 0) in Eqs S65 means the stochastic fluctuations do *not* vanish at the boundaries of [0, 1]^*m*^. Studying whether the resultant process is well-behaved (i.e. guaranteed to remain confined in [0, 1]^*m*^ for all times *t* > 0) is beyond the scope of this work.

For equation (S38), the stochastic analog of the Price equation, we have:

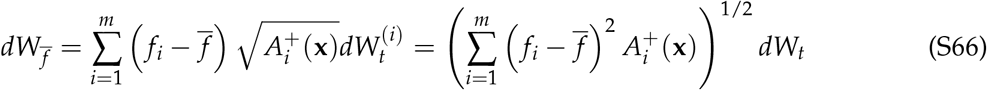

where *W*_*t*_ is now a single one-dimensional Wiener process. This is precisely the term calculated in equation (S53) (barring the 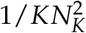 pre-factor), and thus the stochastic term for the mean value is given by:

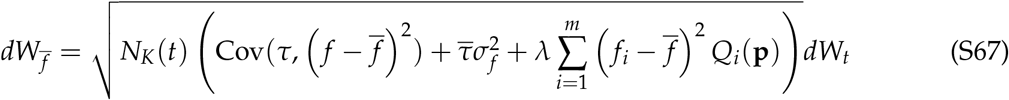

Similarly, for the variance equation (S60), we can write

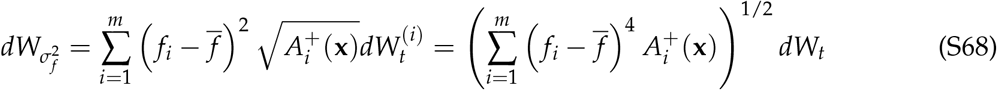

where *W*_*t*_ is now a single one-dimensional Wiener process. A calculation exactly analogous to that done in obtaining (S53) reveals that this term can be written as

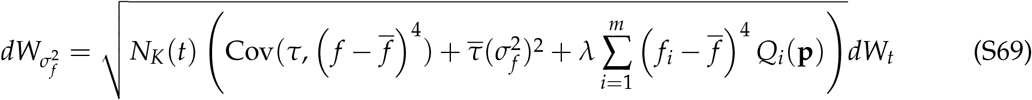

which is the representation used in the main text (with *λ* = 0).

#### Proof of the representation of sums of stochastic integrals with respect to independent Wiener processes

Here, we prove the mathematical result we used above. We stress once again that this is not a new result — we provide the proof here because, while the proof is mathematically straightforward, we were unable to find a suitable citation that explicitly writes down the proof.

##### Lemma.

Let *m* ∈ ℕ. Let 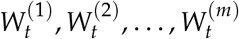 be *m* independent one-dimensional Wiener processes. Let *g*_1_(*x*), *g*_2_(*x*), …, *g*_*m*_(*x*) be *m* ‘nice’ (*L*^2^(ℝ), Lipschitz, *etc*.) real functions. Let

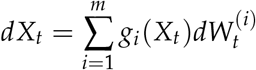

Then, we can always find a *single* one-dimensional Wiener process *W*_*t*_ (on the same probability space) such that

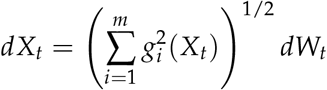

*Proof*. It suffices to prove the *m* = 2 case.

Let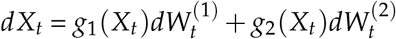. Let us consider the *two*-dimensional process 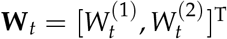 on ℝ^2^. Define a new function *G* : ℝ → ℝ^2^ given by

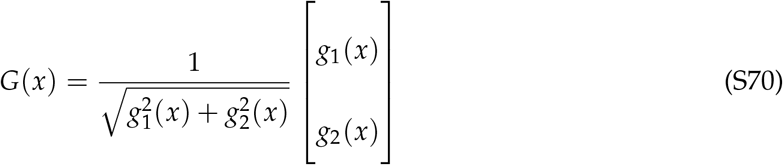

Now, by definition, we have

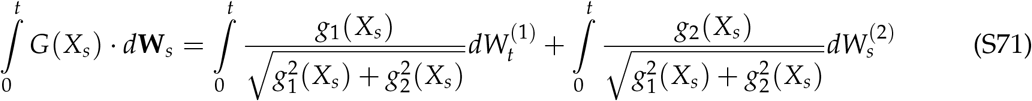

Using the Itô isometry (Karatzas and Shreve, 1998, Chapter 2, Proposition 2.10), we can calculate the quadratic variation of ∫ *G* · *d***W** as

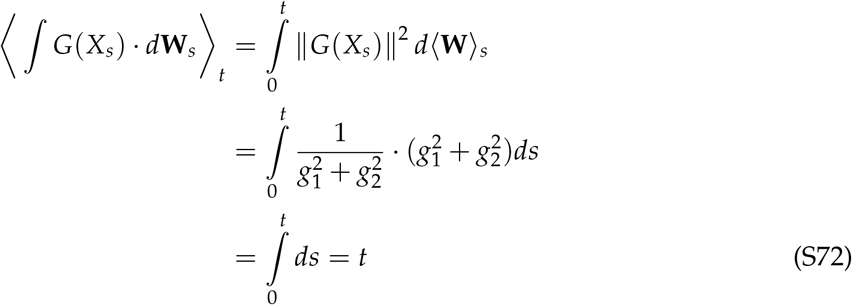

Since ∫ *G* · *d***W** is a stochastic integral, the process 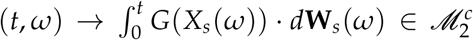 and is thus a continuous martingale. But, by Lévy’s characterization of Brownian motion (Karatzas and Shreve, 1998, Chapter 3, Theorem 3.16), the only continuous martingale *M*_*t*_ that satisfies ⟨*M*⟩_*t*_ = *t* is the Wiener process. Thus, from equation (S72), we are led to conclude that there is a one-dimensional Wiener process *W*_*t*_ on the same probability space such that we can write

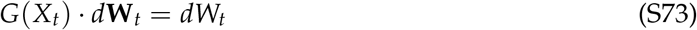

We can now use equation (S71) on the LHS of equation (S73) to write

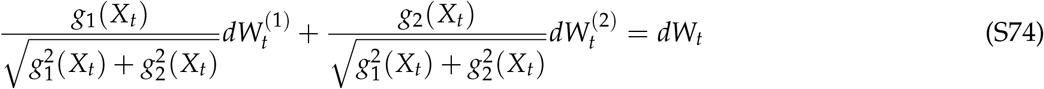

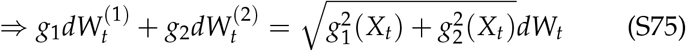

By definition of our original process *X*_*t*_, we can now conclude that

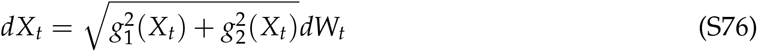

thus completing the proof. □

### S6 The speed density of the stochastic replicator equation for two species

To study the effects of demographic stochasticity on evolutionary dynamics more thoroughly, we use this section to examine the time that the system defined by equation (8) spends at different states. Following McLeod and Day, 2019, we will do this using the speed density. Given any one-dimensional diffusion process *dX*_*t*_ = *µ*(*X*_*t*_)*dt* + *σ*(*X*_*t*_)*dW*_*t*_ defined over an interval [*a, b*] ⊆ ℝ, the *speed density m*(*x*) of the process (Karlin and Taylor, 1981; Etheridge, 2011) is defined as the function

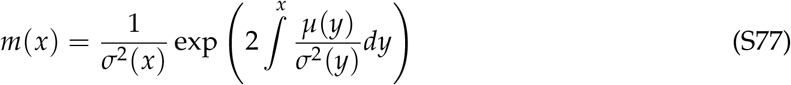

where the lower limit of the integral being left unspecified is meant to denote an indefinite integral evaluated at the point *x* since the choice of the lower limit is arbitrary (Karlin and Taylor, 1981, Chapter 15, Equation 3.10). The speed density is important because it provides information about the long-term behavior of the stochastic process *X*_*t*_. In particular, if there exists a constant 0 < 𝒩 < ∞ such that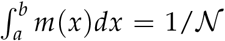, then the stochastic process obtained as the solution to *dX*_*t*_ = *µ*(*X*_*t*_)*dt* + *σ*(*X*_*t*_)*dW*_*t*_ attains a unique stationary state *X*_∞_ as *t* → ∞, and this stationary state has a probability distribution given by (Karlin and Taylor, 1981, Chapter 15, Equation 5.34 along with Chapter 15, Equation 3.10; also see Czuppon and Traulsen, 2021)

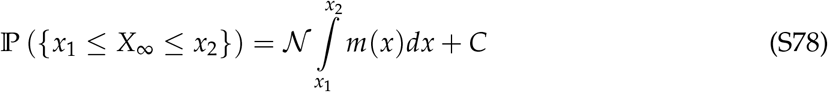

That is to say, the probability density of the stationary state will be given by N *m*(*x*). Regardless of whether such an 𝒩 can be found, the speed density *m*(*x*) always tells us about the time the system spends in the vicinity of the point *x*. More precisely, if we provide an initial condition *x*_0_ ∈ [*a, b*] for the stochastic process obtained as the solution to *dX*_*t*_ = *µ*(*X*_*t*_)*dt* + *σ*(*X*_*t*_)*dW*_*t*_, the expected time taken by this process to exit the interval (*x*_0_ − *ϵ, x*_0_ + *ϵ*) is proportional to *m*(*x*_0_) as *ϵ* → 0 (Karlin and Taylor, 1981, Chapter 15, Remark 3.2). This justified the name ‘speed density’. The quantity *M*(*p*) = ∫^*p*^ *m*(*q*)*dq* is called the speed measure. If a *quasi-stationary distribution* (Collet et al., 2013, definition 2.1) exists, the speed measure describes the quasi-stationary distribution (Collet et al., 2013, section 6.1.1) of the Markov process. A quasi-stationary distribution is known to exist under very general conditions for the kind of processes we study here (Champagnat and Villemonais, 2023).

In our case, we have a stochastic process for the change of type frequencies over time that takes values in [0, 1] and is given by the solution to equation (9). In the rest of this section, we work with equation (8a) and thus do not account for influx terms *λQ*_*i*_. For convenience, let us define

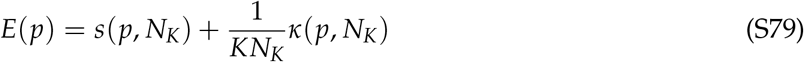

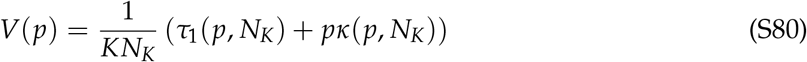

where we have suppressed the *N*_*K*_ dependence of *E* and *V* to reduce clutter. In this notation, equation (9) becomes

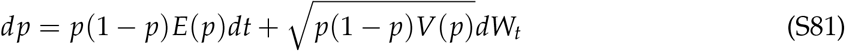

Comparing terms with (S77), we see that the speed density of our process is given by

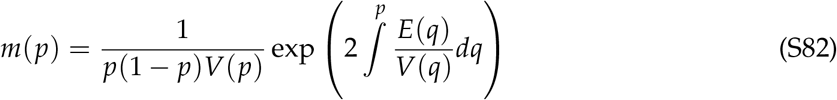

For general functions *E*(*p*) and *V*(*p*), it is very often impossible to analytically calculate or predict the behavior of the complete function defined by (S82). However, since we are primarily interested in which trait frequencies *p* are likely, we can still make analytical progress by examining the derivative *dm*/*dp*. If *dm*/*dp* is a strictly increasing function of *p*, then higher values of frequency *p* are always favored, and species 2 is expected to go extinct more often than species 1. Likewise, if *dm*/*dp* is a strictly decreasing function of *p*, lower frequencies of *p* are favored, and species 1 is expected to go extinct. Lastly, points at which *dm*/*dp* = 0 correspond to extrema of the speed density and can thus be used to find the most likely and least likely values of trait frequency in the system.

We would therefore like to examine the behavior of *dm*/*dp* as a function of p. Differentiating both sides of equation (S82) with respect to *p*, we find

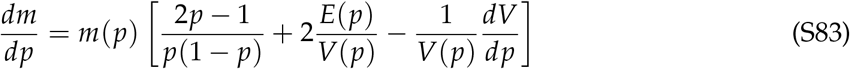

which is Eq. 10 in the main text.

After substituting the functional form of *V*(*p*) from equation (S80), this yields (after some lines of algebra):

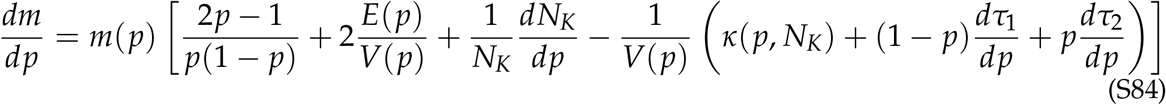

Let us examine each term on the RHS of equation (S84). The first term on the RHS is (2*p* − 1)/*p*(1 − *p*). This expression is (anti)-symmetric about *p* = 0.5 and always drives the system towards the boundaries of [0, 1]. It is thus uninteresting for calculating the sign of *dm*/*dp*.

Since *V*(*p*) must clearly be non-negative in order for equation (S81) to be well-defined, the second term, *E*(*p*)/*V*(*p*), always has the same sign as *E*(*p*). Equation (S84) tells us that the speed density (and thus the stationary distribution, when it exists) also depends on contributions from the *dW*_*t*_ term of equation (S81). We have split this contribution into two separate terms, the third and fourth terms on the RHS of equation (S84), each of which we will examine individually.

The third term on the RHS of (S84) captures the effect of the frequency of species 1 on the per-capita growth rate of the population as a whole. Thus, if species 1 is altruistic, mutualistic, or commensal, then *dN*_*K*_/*dp* will be positive, whereas if the species is spiteful, competitive, or amensal, *dN*_*K*_/*dp* will be negative. The sign of the third term on the RHS of (S84) thus depends on the nature of the ecological interactions that species 1 is involved in — species that increase the per-capita growth rate of the total population are favored, and those that decrease the per-capita growth rate of the total population are disfavored.

The fourth term on the RHS of equation (S84) captures the effects of noise-induced selection acting on differential turnover rates. Since *E*(*p*) also has both a 1/*V*(*p*) factor and a noise-induced selection term, we are better off substituting the functional form of *E*(*p*) from (S79) into equation (S84) and collecting all terms with a 1/*V*(*p*) factor so as to collect all terms corresponding to selection (both classical and noise-induced). Upon doing this, we obtain

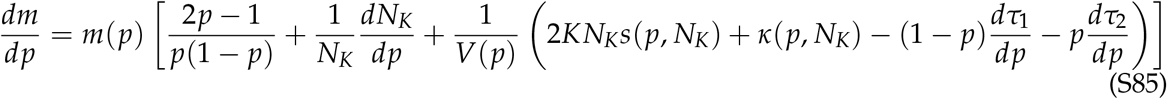

The interpretations of the first two terms on the RHS of (S85) have already been explained above. Since *V*(*p*) is always non-negative, we only need to look at the sign of the expression

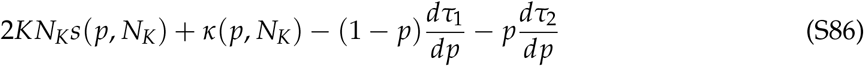

The first term of (S86) is the effect of classical selection and has the same sign as the selection coefficient *s*(*p, N*_*K*_). Notice that since this term is 𝒪(*K*) whereas all other terms in equation (S85) are 𝒪(1), this term dominates the dynamics when *K* is large, again indicating that the effects of natural selection dominate in large populations with non-zero selection coefficient. If instead *Ks*(*p, N*_*K*_) is small, either through a small population size, weak selection (or no selection), or both, the last three terms of (S86) play a stronger role. The second term of (S86) is simply the noise-induced selection coefficient *κ*(*p*), and is thus positive whenever *τ*_1_ < *τ*_2_. This term thus captures the direct mechanism of noise-induced biasing via noise-induced selection and causes the speed density to be biased towards the species with lower per-capita turnover rates. However, when the turnover rates depend on the frequency of traits, the last two terms of (S86) also affect the shape of the speed density. These two terms capture the *indirect* mechanism of noise-induced biasing via frequency-dependent turnover and bias the population towards those states which are associated with lower rates of change of the population as a whole. Note that the total strength of the effect of frequency dependence of a given type is inversely proportional to the current frequency of that type (*dτ*_1_/*dp* is multiplied by 1 − *p* and *dτ*_2_/*dp* is multiplied by *p*). This reflects the intuitive observation that stochastic effects on a focal type should play a stronger role when that type is rare.

### S7 The infinite population limit recovers standard equations of population biology

In this section, we show how our SDEs recover several classic equations of population biology in the infinite population size limit.

#### Replicator-mutator equation

If w take *K* → ∞ in (S29), we obtain an ODE that reads:

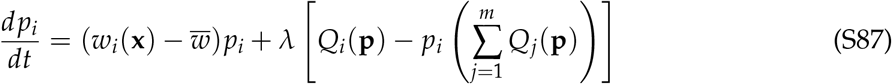

The first term of (S87) describes changes due to faithful (non-mutational) replication, and the second describes changes due to mutation. For this reason, equation (S87) is called the *replicator-mutator equation* in the evolutionary game theory literature, where the individual ‘types’ are interpreted to be pure strategies and the influx rate *λ* is a mutation rate, denoted by *µ*. If in addition, each *w*_*i*_(**x**) is linear in **x**, meaning we can write *w*_*i*_(**x**) = ∑_*j*_ *a*_*ij*_*x*_*j*_ for some set of constants *a*_*ij*_, then we get the replicator-mutator equation for matrix games, and the constants *a*_*ij*_ form the ‘payoff matrix’. As is well-known, the replicator equation (without mutation) for matrix games with *m* pure strategies is equivalent to the generalized Lotka-Volterra equations for a community with *m* − 1 species (Hofbauer and Sigmund, 1998), providing the connection to community ecology. Equation (S87) is also equivalent to Eigen’s *quasispecies equation* from molecular evolution if each ‘type’ is interpreted as a genetic sequence and each *w*_*i*_(**x**) is a constant function (Page and Nowak, 2002).

#### (Dynamical) Price equation

Taking *K* → ∞ in equation (S38) recovers the Price equation as the infinite population limit. Here, we mean the Price equation as formulated in continuous time with time-varying characters (Lion, 2018; Day et al., 2020).

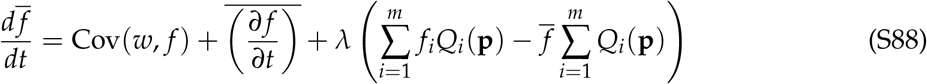

Many authors additionally assume that the quantity *f* does not itself change over time at the type level, meaning that *∂ f*_*i*_/*∂t* ≡ 0 ∀ *i* and the feedback term thus disappears. This yields a somewhat more familiar equation in continuous time (Lion, 2018). Standard texts also usually use a version formulated in discrete time that is more general for single-step changes, but is dynamically insufficient (Frank, 2012; Queller, 2017).

#### Fisher’s fundamental theorem of natural selection

Taking *K* → ∞ in (14) and noting that the process tends to a deterministic process as *K* → ∞, as noted in section S7 (and thus the expectation value in the infinite population case is superfluous), we obtain an ODE:

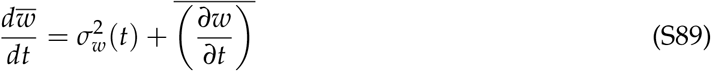

This is Fisher’s fundamental theorem in the presence of ecological feedbacks to fitness (Frank and Slatkin, 1992; Kokko, 2021).

#### The equation for trait variances that appears in Lion, 2018

Taking *K* → ∞ in equation (S60) yields

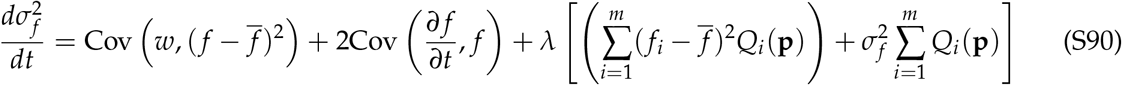

This is precisely equation (14) in Lion, 2018 with influx terms *λQ*_*i*_.

### S8 Noise-induced biasing in various specific contexts

In many social evolution models, cooperators are predicted to go extinct in infinite populations but are actually favored by evolution in finite, fluctuating populations, causing a ‘reversal’ in the direction of evolution predicted by natural selection (Houchmandzadeh and Vallade, 2012; Chotibut and Nelson, 2015; Constable et al., 2016; McLeod and Day, 2019). McLeod and Day, 2019 have recently shown that such reversals can occur in a wide array of social evolution models due to the same effect that we recognize here as noise-induced biasing. Formally, all the models presented in McLeod and Day, 2019 can be recovered in our framework by setting *m* = 2 and *s*(**x**) = −*ϵc*(**x**) for a constant *ϵ* ∈ ℝ and a non-negative function *c*(**x**) in our stochastic replicator-mutator equation (Eq. S29). The function *T*(*p*) in McLeod and Day, 2019 — a quantity that varies in the various models they study — is precisely the mean turnover 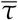 in our framework.

In evolutionary epidemiology, models have shown that reduced virulence is more important than increased transmission rate for pathogen spread in finite, fluctuating populations, especially when the population size is small (Humplik et al., 2014; Parsons et al., 2018; Day et al., 2020). Indeed, if the population is small or selection is weak, slower strains can have higher fixation probabilities than faster strains even if the slower strain has a lower basic reproduction ratio (*R*_0_) than its competitor, causing a complete reversal in the direction of evolution predicted in infinite populations (Parsons et al., 2018). These results have recently been explained in a generic manner using both a replicator-mutator/’stochastic adaptive dynamics’ approach (Parsons et al., 2018) and a two-species Price equation formalism (Day et al., 2020), though both these papers use assumptions and language particular to evolutionary epidemiology. We note that equation (2.5) in Parsons et al., 2018 is exactly equivalent to our stochastic replicator-mutator equation with no mutation (equation (S29) with *λ* = 0) upto a change in notation upon substituting the specific birth and death rate functions chosen in their paper into our equation (S29). Similarly, equation (5.1) in Day et al., 2020 is exactly equivalent to our stochastic Price equation for 2 species (equation (S38) with *m* = 2) if we write out *w* and *τ* in terms of per-capita birth and death rates. Our work can, therefore, be used to recapitulate these results and show that the effects they illustrate are not particular to epidemiological models.

Lastly, noise-induced biases in population dynamics may also appear in the infinitesimal mean as a deviation from the expected trajectory if we project the ecological dynamics onto a ‘slow manifold’ through a separation of timescales argument, a common procedure for reducing the dimension of stochastic dynamical systems (Constable et al., 2013; Parsons and Rogers, 2017). A change of variables via a projection of the dynamics onto a manifold is responsible for the ‘noise-induced effects’ that appear in purely ecological models (i.e. models of population densities) where dynamics are projected onto a manifold describing populations that are at equilibrium over short timescales (Constable et al., 2016; Chotibut and Nelson, 2017; Mazzolini and Grilli, 2023). A change of variables via projection onto a manifold is also at the heart of the stochastic ‘drift-induced selection’ that drives evolutionary transitions between male and female heterogamety (XX/XY to ZW/ZZ and vice versa) in stochastic models of the evolution of chromosomal sex determination systems (Veller et al., 2017; Saunders et al., 2018). In models of sex determination, the projection is onto a manifold describing populations in which the sex ratio is 1:1 (Veller et al., 2017; Saunders et al., 2018). However, note that an additional stochastic term in a projected version of population densities need not lead to selection in the evolutionary sense, namely in terms of changes in trait frequencies (McLeod and Day, 2019).

### S9 Connections with some other general frameworks

Our equations generalize Lion’s (2018) general framework of infinite population deterministic eco-evolutionary dynamics to finite, fluctuating populations — taking *K* → ∞ in Eq. S29, Eq. S38, and Eq. S60 recover equations (6), (11), and (14) in Lion, 2018 respectively. Just like in the deterministic setting (Lion, 2018), equations for trait means (the Price equation, Eq. S38) and trait variances (Eq. S60) can be systematically derived from the equation for changes in trait frequencies (the replicator-mutator equation, Eq. S29) in our framework through repeated application of Itô’s formula. If we assume that the quantity *f* follows a Gaussian distribution, then the mean and variance completely characterize the distribution, and thus, Eq. S29, Eq. S38, and Eq. S60 together specify the complete stochastic dynamics of the system.

Rice has proposed a stochastic version of the Price equation (Rice, 2020 and references cited therein). Like the original Price equation, Rice’s equations are formulated as a general decomposition of the phenotypic change between two given populations. They are thus the true stochastic analog of the original Price equation, whereas our version, Eq. S38, is the analog of Lion’s (2018) version of the Price equation in a continuous time, dynamically sufficient setting. Rice’s derivations also treat fitness as fundamental, whereas we derive suitable notions of fitness and turnover from demographic first principles. As a consequence, the ‘extra’ stochastic term corresponding to noise-induced selection that appears in our equations fundamentally emerges from the stochasticity of the underlying births and deaths of organisms and is thus of ecological/demographic origin, whereas the ‘extra’ stochastic term in Rice’s equations emerges from the stochasticity of fitness alone when viewed as a random variable (Rice, 2020). It thus need not, to the best of our knowledge, correspond to the same effect we identify here.

There are also deep connections between the equations we present in this paper and those of Week et al. (2021). Informally, the equations presented in Week et al. (2021) can be recovered from our framework if we let the number of types *m* go to infinity and replace sums with integrals. Making this intuition precise is rather (mathematically) involved (Bhat, 2024), but we provide a summary of the idea below. We first require an intuitive explanation of how quantitative traits can be modeled in a stochastic birth-death framework.

Populations bearing discrete traits can be characterized by a vector **n** = [*n*_1_, *n*_2_, …, *n*_*m*_] enu-merating the number of individuals of each type. However, this cannot be done for populations bearing quantitative traits since infinitely many distinct types may arise. Instead, we characterize the population as a *finite measure* 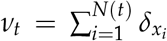 by placing a Dirac delta mass 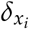 at the loca-tion *x*_*i*_ corresponding to the trait value of the *i*^th^ individual in a population consisting of *N*(*t*) individuals at time *t*. The population dynamics can then be described in terms of a measure-valued stochastic process (or, in physics language, a ‘stochastic field theory’) that tracks how *ν*_*t*_ evolves over time. Since we assume that every type has the same functional form for the birth and death rates, in the discrete case, the set of birth rate functions {*b*_*i*_(**n**)}_*i*=1,2,···, *m*_ (and analogously the death rate functions) can alternatively be viewed as a single function *b* from {1, 2, …, *m*} × [0, ∞)^*m*^ → [0, ∞) whose *i*^th^ component gives us the birth rate of the *i*^th^ individual. This alternative view makes it clear what the analogous birth rate ‘function’ should look like for quantitative traits: The function *b* (and similarly the death rate function) is now replaced by a functional *B* : 𝒯 × ℳ (𝒯) → [0, ∞), where 𝒯 ⊆ R is the trait space describing the set of allowed trait values and ℳ (𝒯) is the space of ‘functions’ (finite measures) on 𝒯. One can then show that under some suitable assumptions, it is possible to carry out an analog of the ‘system size expansion’ we use in the supplementary of this manuscript to approximate the finite measure *ν*_*t*_ as a function *ϕ*(*x, t*) describing the distribution of population densities across the trait space. One can then derive stochastic partial differential equations (SPDEs) for how the distribution of trait frequencies, the mean value of any trait, and the variance of any trait change over time. These SPDEs end up being precisely the quantitative trait analogs of the SDEs we present in the main text (ínformally, the ‘*m* → ∞ limit’ of the *m*-dimensional system of SDEs we present in the main text of this manuscript).

With the broad outline explained above in mind, the connection of our equations with the equations presented in Week et al. (2021) can be made precise. Week et al. (2021) work with an asexual model with Gaussian mutation (their ‘SAGA’ - stochastic asexual Gaussian alleles). In their model, individuals give birth to offspring with mutations, with mutants normally distributed about the parent trait value with some small variance *µ* (their notation). Such mutation is approximated in their framework by the Laplacian of the function describing the distribution of population densities (i.e. as ‘diffusion’ across the trait space). Given a type *x* (an index *i* in the discrete case, now a real number that is also the trait value) in a population *ϕ* (a vector **x** in the discrete case, now a function), the connection between our equations and Week et al. (2021)’s equations can be formally made via the relations (our notation on the left, Week et al. (2021)’s notation on the right):

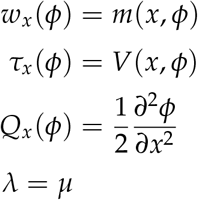

(Compare Eq. 3 in our manuscript with Eq. 20 in Week et al. (2021)). Further, in the equation for the mean value (Their Eq. 21b, our Eq. S38) and variance (Their Eq. 21c, our Eq. S60) of a quantity, they restrict themselves to the special case *f*_*x*_(*ϕ*) = *x*, the quantitative trait analog of what in the discrete trait notation would be *f*_*i*_(**x**) = *i*. In this case, the influx terms of our equations will vanish for Week et al. (2021)’s choice of *Q*_*x*_(*ϕ*). Formally, the influx terms in Eq. S38 and Eq. S60 are all integrals of the form

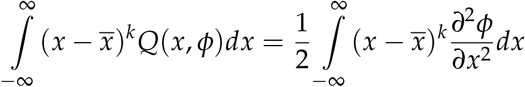

with either *k* = 1 or *k* = 2, and vanish upon using integration by parts and discarding the boundary terms. Thus, the influx terms do not contribute to the dynamics, which is why Eq. 21b in Week et al. (2021) corresponds to our Eq. S38 and Eq. 21c in Week et al. (2021) corresponds to our Eq. S60. A derivation of the quantitative trait analogs of the equations we present in this paper, along with a more precise explanation of the equivalence with Week et al., 2021, appears in Bhat, 2024.

### S10 A more detailed explanation of the example in the main text

In this section, we flesh out the example introduced in Box 5 in more detail. To illustrate when noise-induced selection can be important for population dynamics, we use a simple biologically motivated example in this section. Several abiotic factors such as temperature and pH are known to be ecological ‘rate modulators’ that affect either the birth rate or death rate of organisms, with obvious consequences for evolutionary dynamics (Fronhofer et al., 2023). To see how demographic stochasticity may affect the effect of ecological rate modulators on evolutionary dynamics, consider here two competing phenotypes, which we denote by 1 and 2. Though we stick to this ‘rate modulation’ language henceforth, another potential interpretation of the model we study below comes from epidemiology: In this case, the two types can be thought of as two competing strains of pathogens, a ‘rate modulator’ that affects birth rates can be thought of as affecting transmission rate, and a ‘rate modulator’ that affects death rates can be thought of as affecting virulence (Parsons et al., 2018). We consider the case where type 1 is affected by the ecological rate modulator but type 2 is not. For simplicity, we assume the population is closed with no mutations during birth (*i.e. λ* = 0). Below, we use *p* to denote the frequency of type 1 individuals in the population.

For pedagogical clarity, we assume that rate modulation occurs by simply shifting the birth and/or death rate by a constant. In equations, this can be modelled via the relations:

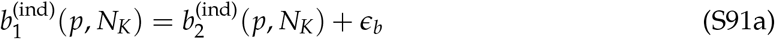

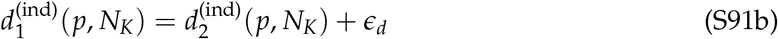

where *ϵ*_*b*_ and *ϵ*_*d*_ are real numbers describing the effect of the ecological rate modulator on the birth and death rates respectively. Using the definitions of *s* and *κ*, we find

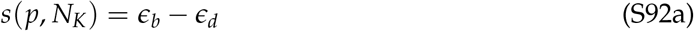

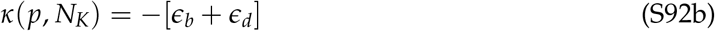

Note that if *ϵ*_*b*_ = 0, *ϵ*_*d*_ < 0, both *s* and *κ* are positive, whereas if *ϵ*_*b*_ > 0, *ϵ*_*d*_ = 0, *s* > 0 but *κ* < 0. In other words, if type 1 has a decreased death rate (virulence in the epidemiological case) but identical birth rate relative to type 2, type 1 is favored by both natural selection and noise-induced selection. On the other hand, if type 1 has an increased birth rate (transmission rate in the epidemiology case) but an identical death rate relative to type 2, type 1 is favored by natural selection but disfavored by noise-induced selection. Thus, all else being equal, reducing the death rate is generically more favorable than increasing the birth rate by an analogous amount, an observation that has been made in finite population models in epidemiology (Parsons et al., 2018), social evolution (McLeod and Day, 2019), life-history evolution (Alexander and Wahl, 2008), and cancer biology (Raatz and Traulsen, 2023).

For the rest of this example, we assume that *ϵ*_*b*_ > 0, *ϵ*_*d*_ > 0, *i.e*. that type 1 has both an increased birth rate and an increased death rate compared to type 2. We may now ask, when is the outcome of evolution different from that expected by infinite population dynamics?

#### Noise-induced biasing in the absence of natural selection

First, consider the situation *ϵ*_*b*_ = *ϵ*_*d*_ = *ϵ*. This corresponds to the two types having the same growth rate, but type 1 having a faster pace of life than type 2. The selection coefficient and noise-induced selection coefficient are

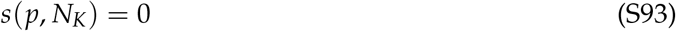

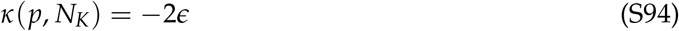

Thus, as expected, natural selection does not operate in the system. In the infinite population limit, natural selection is the only force that affects population dynamics and we thus expect any initial frequency *p*_0_ of type 1 individuals to remain unchanged in the population (to see this, take *K* → ∞ in Eq. 9). Over short timescales, the effects of demographic stochasticity can be observed by looking at the expected change in frequency 𝔼[*dp*]. Using Eq. 9 and substituting the functional forms given by Eq. S92, we find

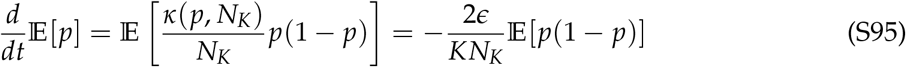

Since the RHS of Eq. S95 is always negative for *p* ∈ (0, 1), we can infer that if the system begins at any initial frequency *p*_0_ ∈ (0, 1), the proportion of type 1 individuals is expected to decrease on average. If *ϵ*_*b*_ = *ϵ*_*d*_, the ecological rate modulator is thus detrimental to the evolutionary fate of type 1 individuals over short time scales in finite populations, despite infinite population models predicting neutrality. This result is a manifestation of the ‘direct’ mechanism of noise-induced biasing via the Gillespie effect (noise-induced selection): All else being equal, a faster pace of life comes with a greater variance in change of population density within a given time interval since there are simply more stochastic birth/death events taking place.

However, the speed density S77 depends not only on the expected change of frequency alone but also on the variance in the change of frequencies. This stochastic effect, captured by the *dW* term in Eq. 9, depends on the functional form of *τ*_1_(*p, N*_*K*_) (and not merely the difference *κ* = *τ*_2_ − *τ*_1_), which we have not yet specified in our model (Eq. S91). For simplicity, let us assume that the turnover rates *τ*_*i*_ depend linearly on *p*. Specifically, let us assume that *τ*_1_(*p, N*_*K*_) = *bp* + *c*, where *b* and *c* are constants. *c* can be viewed as an ‘intrinsic’ turnover rate, and *b* as a frequency-dependent component that may be either positive or negative. We are therefore restricting ourselves to linear frequency dependence of *τ*_1_, but allowing both positive and negative frequency-dependence, with the strength of frequency-dependence controlled by |*b*|. For convenience, let *a* = 2*ϵ*_*d*_ = −*κ*(*p*). Note that since *τ*_1_ is the sum of two rates and *p*(1 − *p*)*V*(*p*) is the infinitesimal variance of the trait frequency SDE, the parameters *a, b*, and *c* must be chosen such that *τ*_1_(*p*) = *bp* + *c* > 0, *V*(*p*) = (*b* − *a*)*p* + *c* > 0 ∀ *p* ∈ [0, 1] for the system to be biologically meaningful. In particular, *τ*_1_(0, *N*_*K*_) and *τ*_1_(1, *N*_*K*_) must be non-negative, and we must thus have *τ*_1_(0, *N*_*K*_) = *c* ≥ 0 and *τ*_1_(1, *N*_*K*_) = *b* + *c* > 0. We must also have *V*(1) > 0, and thus *b* + *c* − *a* ≥ 0.

Since we would like *κ* to still be given by Eq. S94, this automatically specifies *τ*_2_ as *τ*_2_ = *bp* + *c* − 2*ϵ*. Thus, we assume *τ*_1_ and *τ*_2_ change in the same direction (increase or decrease) as the frequency of type 1 individuals increases.

In this case, we can exactly solve for the speed density. Note that we have

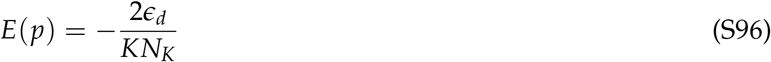

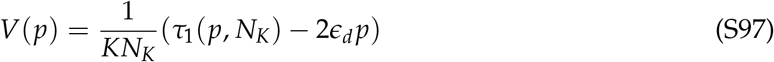

where *a* = 2*ϵ*_*d*_ = −*κ*(*p*). In our new notation, Eq. S96 and S97 become

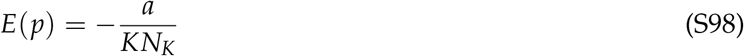

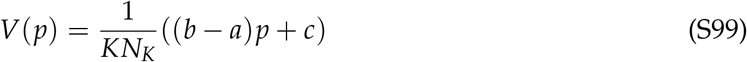

The speed density of the system can be written (from Eq. S82) as

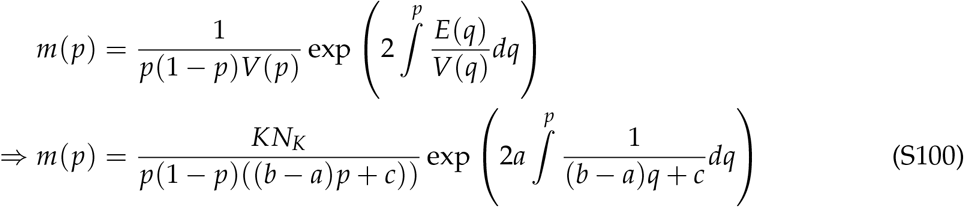

We need to distinguish between two cases based on whether or not *V* is frequency-dependent.

#### Case 1: No frequency dependence in V(p)

If *a* = *b, i.e*. the frequency dependence of *τ*_1_ is positive with strength exactly equal to 2*ϵ*_*d*_, Eq. S100 becomes

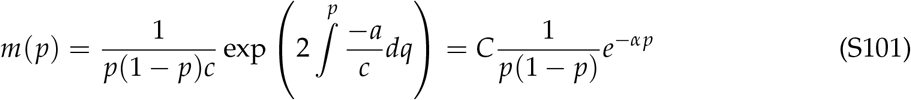

where *α* = 2*a*/*c* > 0 is a positive constant, and we use *C* to denote a constant whose precise value is irrelevant (and thus may change from line to line below — the important thing is that *C* does not depend on *p* and thus plays the role of a normalization constant).

The shape of the distribution given by Eq. S101 can be thought of as the combination of two components: The term *p*(1 − *p*) is symmetric with respect to the transformation *p* → 1 − *p* (i.e. symmetric about *p* = 0.5) and thus does not favour either type of individual, whereas *e*^−*αp*^ is a strictly decreasing function of *p* and thus always favours lower frequencies of type 1 individuals. If *α* is very small, the effect of *e*^−*αp*^ is negligible and the distribution of types is approximately a symmetric ‘U-shaped’ parabola centered at 0.5, with *p* = 0.5 being the least likely frequency. This is the expectation we would have if neutral genetic drift was the only force at play: The distribution is (approximately) symmetric with respect to the transformation *p* → 1 − *p*, with *p* = 0 and *p* = 1 being the most likely states and *p* = 0.5 being the least likely state.

If, instead, *α* is not small, the function *e*^−*αp*^ decays quickly and biases the distribution towards lower values of *p*. In this case, the function is a distorted U-shape, with the minimum point being somewhere in (1/2, 1). The extent of bias towards lower values of *p* increases as *α* increases.

Thus, in the case where 2*ϵ*_*d*_ = *dτ*_1_/*dp*, we can conclude that lower frequencies of type modulators are always associated with a higher speed density, and the biasing is stronger as the ratio of the rate modulation (*ϵ*_*d*_) to the intrinsic frequency-independent turnover rate (*c*) increases. Note that the shape of the speed density (and thus the extent of deviation from neutrality in terms of time spent/likelihood of observation conditioned on non-fixation) does *not* depend on the total population size *KN*_*K*_.

#### Case 2: Frequency dependence in V(p)

Assuming *a*≠ *b*, we can calculate the exponential term in Eq. S100 as

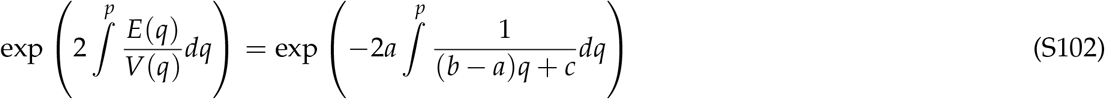

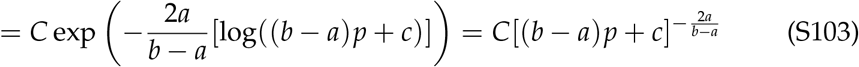

where we once again use *C* to denote a multiplicative constant whose precise value is irrelevant. Thus, the speed density S100 is given by

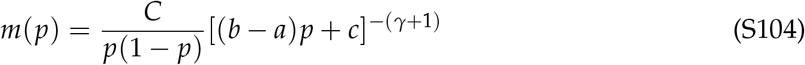

where we have defined *γ* = 2*a*/(*b* − *a*).

Equations S101 and S104 are exact solutions for the speed density, and are the quantities plotted in Figure 3 in Box 5. For convenience, we reproduce the figure below

**Figure S1:**
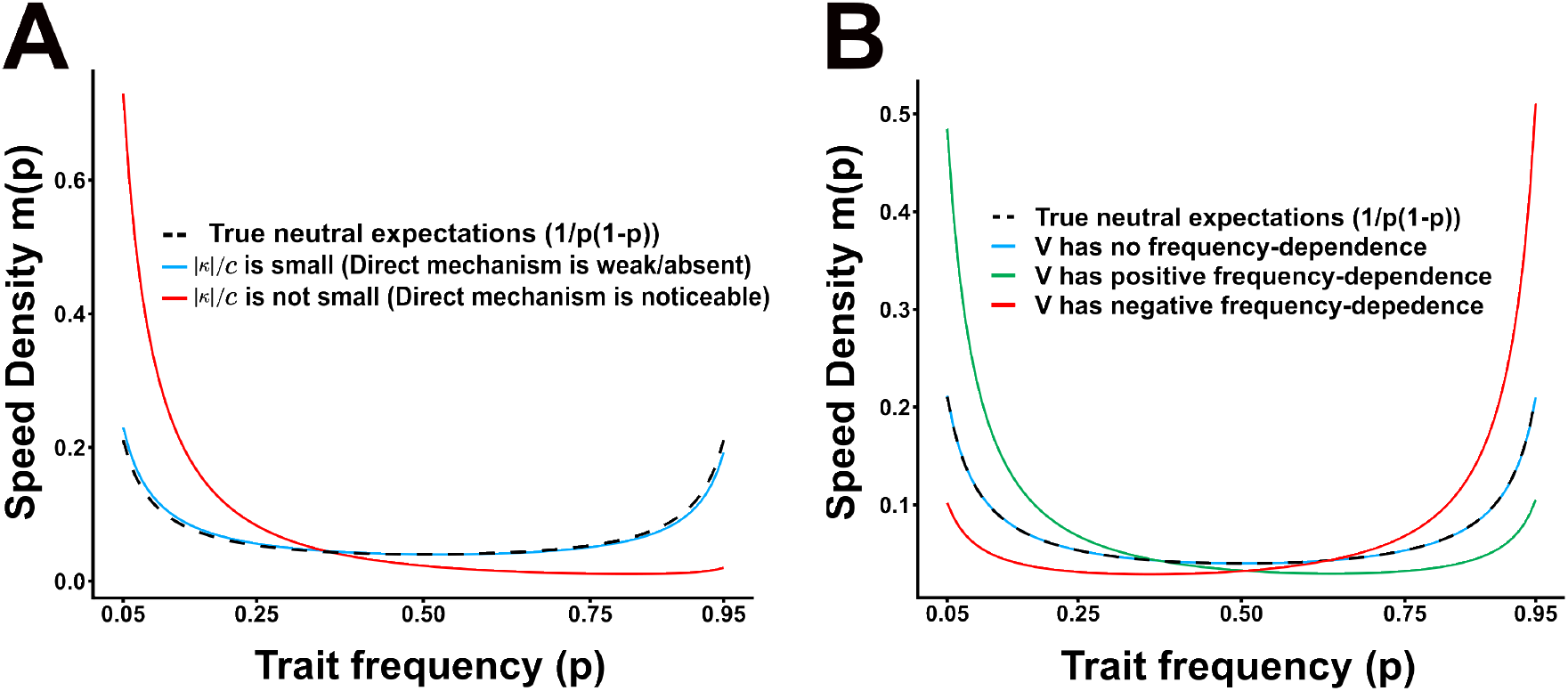
Two distinct noise-induced effects that bias evolutionary dynamics. **A**. If the magnitude of the noise-induced selection coefficient *κ* = − 2*ϵ* is large relative to the intrinsic turnover rate *c*, the direct mechanism of noise-induced biasing operates. Parameters are chosen such that *V*(*p*) = *τ*_1_ + *pκ* = (*b* − 2*ϵ*)*p* + *c* is not frequency-dependent (blue: *ϵ* = 0.5, *b* = 1, *c* = 10; red: *ϵ* = 0.5, *b* = 1, *c* = 0.5). **B**. The speed density can also be biased if *V*(*p*) is frequency-dependent. This indirect mechanism of noise-induced biasing favours the type that reduces *V*(*p*). Parameters in this panel are chosen such that | *κ* | is small relative to *c* and thus the strength of the fast mechanism is negligible (blue: *ϵ* = 0.025, *b* = 0.05, *c* = 10; green: *ϵ* = 0.025, *b* = 50, *c* = 10; red: *ϵ* = 0.025, *b* = −8.5, *c* = 10)

Figure S1 illustrates two distinct ways in which stochasticity alone can bias trait frequency distributions in finite, fluctuating populations. If dynamics are truly neutral (in the sense of the two types being exactly equivalent) and the system begins with *p* = 0.5, then both types are equally likely to increase/decrease. The speed density is thus equal to 1/*p*(1 − *p*) (up to a constant). Noise-induced effects can bias this distribution in two distinct ways (Box 4): (1) The noise-induced selection coefficient *κ* can bias the expected trajectory. This is noise-induced selection and can be identified with the Gillespie effect from bet-hedging theory as a selection for reduced variance in density change *dx*_*i*_. Since *κ* = −2*ϵ* < 0, noise-induced selection always favours the type with the slower pace of life (Fig. S1A). (2) A second noise-induced effect appears as a biasing of the speed density via the last term of Eq. 10. We call this an ‘indirect’ mechanism of bias because it is not observable in terms of expectations 𝔼[·] but is visible in terms of time spent in various states. For our example, we can calculate

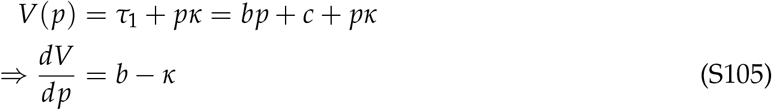

Equation S105 tells us that this indirect mechanism favours type 1 if *b* < *κ*, type 2 if *b* > *κ*, and does not operate if *b* = *κ*. Thus, the indirect mechanism may cause biases in either the same direction or the opposite direction of noise-induced selection based on the details of the frequency-dependence of the per-capita turnover rates (Fig S1B).

#### Noise-induced biasing in the presence of natural selection

Consider now instead a situation in which the rate modulator affects the birth rate more than it does the death rate (*i.e. ϵ*_*b*_ > *ϵ*_*d*_ > 0). In this case, the selection coefficient *s* in Eq. S92 is always positive, and thus natural selection always favors type 1 individuals. As before, noise-induced biases may manifest in two distinct ways. First, noise-induced selection may invert the direction of the expected trajectory 𝔼[*dp*/*dt*]. Noise-induced biasing may also occur through the indirect mechanism. We examine the two possibilities one by one.

Since *s* > 0, we can use Eq. 9 to say the expected trajectory is in the opposite direction of infinite population predictions if *s* + *κ*/*KN*_*K*_ < 0. Using Eq. S92, we see that this is equivalent to

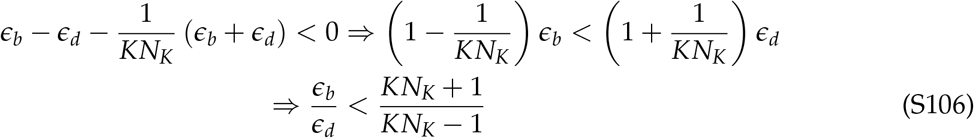

Using inequality S106 in Eq. S92 a, we can arrive at the inequality

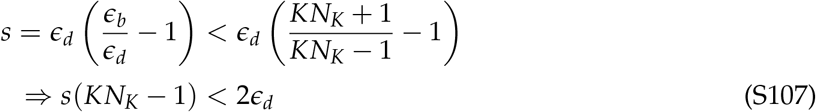

Thus, noise-induced selection can reverse the expected trajectory of evolutionary dynamics when the product *s*(*KN*_*K*_ − 1) is sufficiently small, *i.e*. when either *selection is weak* (*s* is small), *populations are small* (*KN*_*K*_ is small), or both.

We now also examine the contributions of the noise terms to the speed density. We see from Eq. 10 that we can say type 1 is favoured by the indirect mechanism when *dV*/*dp* < 0. Using the definition of *V* from Eq. 11b and substituting the functional forms given by Eq. S92, we see that *dV*/*dp* < 0 is equivalent to

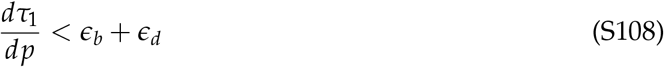

If *τ*_1_ is a constant, *i.e*. the per-capita birth rates 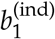 and 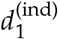 do not depend on population composition, inequality S108 will automatically be satisfied as long as there is some rate modulation in the system (*i.e. ϵ*_*b*_ and *ϵ*_*d*_ are not both 0). If *τ*_1_ is frequency dependent, S108 is satisfied whenever *τ*_1_ exhibits negative frequency dependence, though it may also be satisfied if *τ*_1_ exhibits weak positive frequency dependence. We do not explore the effects of the indirect mechanism further for the sake of conciseness. However, we note that since we already studied the behaviour of *E*(*p*) above, it is now straightforward to determine from Eq. 10 when this latter effect combines with *E*(*p*)/*V*(*p*) to make the RHS of Eq. 10 positive. In Supplementary section S11, we provide an example system in which noise-induced selection can never reverse the expected trajectory 𝔼[*dp*/*dt*], but where the indirect mechanism of noise-induced biasing may nevertheless affect the distribution of types in the population. In Supplementary section S12, we provide an example of a stochastic Lotka-Volterra competition model with both natural selection and mutation in which noise-induced selection can reverse the direction of evolution predicted by natural selection-mutation balance.

### S11 An example in which noise-induced selection can never reverse the direction of evolution over short timescales, but where the indirect mechanism may still operate

Consider a slightly modified version of the example covered in the main text. Consider here two types in which rate modulation decreases the birth rate and increases the death rate of type 1 individuals. In equations, such modulation can be modelled via the relations:

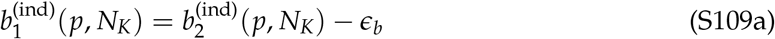

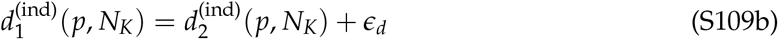

where *ϵ*_*b*_ and *ϵ*_*d*_ are non-negative real numbers describing the effect of the ecological rate modulator on the birth and death rates respectively. Note that in this case, *ϵ*_*b*_ cannot be arbitrarily large: we require 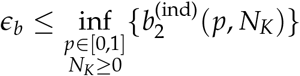 to avoid negative birth rates. As in the main text, we can calculate the selection coefficient and noise-induced selection coefficient to find

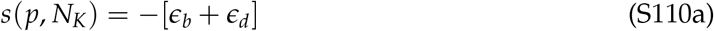

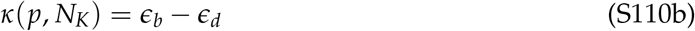

Here, *s* is always negative whenever there is some rate modulation in the system (*i.e. ϵ*_*b*_ and *ϵ*_*d*_ are not both 0), and thus natural selection always favors type 2 over type 1. Note that here, when evolution is neutral with respect to natural selection (*s* = 0), we must have *ϵ*_*b*_ = *ϵ*_*d*_ = 0. In this case, 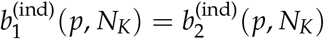 and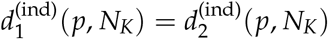, and thus the two types are exactly equivalent in every respect.

We first examine when the sign of 𝔼[*dp*/*dt*] is reversed relative to infinite population expectations, i.e. when noise-induced selection can reverse evolutionary outcomes. Since *s* < 0, we can use Eq. 9 to say the expected trajectory is in the opposite direction of infinite population predictions if *s* + *κ*/*KN*_*K*_ > 0. Using Eq. S110, we see that this is equivalent to

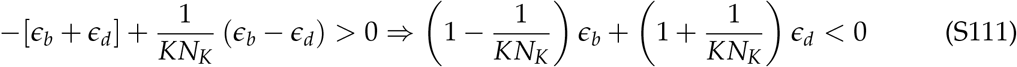

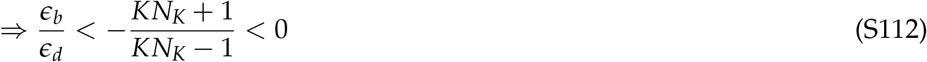

Since *ϵ*_*b*_ and *ϵ*_*d*_ are both non-negative, so is their ratio, and thus inequality S112 can never be satisfied. We therefore conclude that noise-induced selection *cannot* reverse the sign of 𝔼[*dp*/*dt*] relative to infinite population expectations in this case.

However, the speed measure may still be biased due to the indirect mechanism. We see from Eq. 10 that type 1 may be favoured via the indirect mechanism if *dV*/*dp* is sufficiently negative. Using the definition of *V* from Eq. 11b, we see that *dV*/*dp* is negative whenever

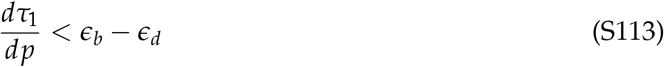

Note, however, that for this system, since *E*(*p*) will always be positive, *dV*/*dp* < 0 is a necessary but not a sufficient condition for deviation from infinite population expectations — we also require *dV*/*dp* to be large enough in magnitude relative to *E*(*p*) to ensure that the RHS of Eq. 10 as a whole becomes positive.

### S12 An example of non-neutral competition where evolution does not proceed in the direction of natural selection due to noise-induced effects

In this section, we provide an example of resource competition with both natural selection and mutation in which noise-induced selection reverses the direction of evolution predicted by natural selection.

Consider a community that contains two types of birds, say type 1 and type 2. These birds compete for limited resources, but in a peculiar manner: Though the two birds feed on different food sources, the trees that type 1 birds use for nesting are the same as those that the type 2 birds rely on for food. Both types are territorial and do not tolerate other individuals of either type on either their nesting or feeding sites. Thus, competition between the two types affects the *birth rate* of type 1 birds (because they can’t find good nesting sites) but the *death rate* of type 2 birds (due to starvation), whereas intratype competition affects the death rate in both cases due to competition for food sources. We also assume that when individuals give birth, they may give birth to offspring of the opposite type (due to mutations) at a rate *λ* > 0. Thus, the influx rate *λ* here is a mutation rate, and we will therefore denote it by *λ* = *µ* to align with standard notational conventions. Let *n*_*i*_ be the number of type *i* individuals (which may vary over time). Assuming trees and birds are both randomly distributed through the landscape and the population dynamics of birds has linear density dependence, the simplest model that can incorporate these features of resource competition is given by:

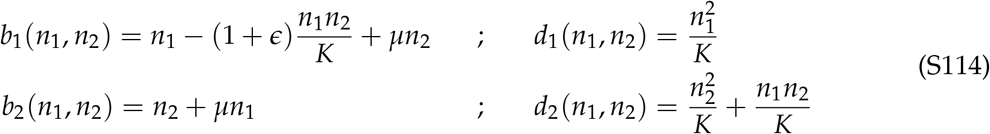

where *K* is a carrying capacity for the habitat, similar to Lotka-Volterra competition, and *ϵ* is a parameter, which as we shall see below, quantifies which type has a competitive advantage. Note that *ϵ* must be chosen such that *b*_1_(*n*_1_, *n*_2_) is always non-negative.

Moving to density space via the change of variables *x*_*i*_ = *n*_*i*_/*K*, letting **x** = [*x*_1_, *x*_2_]^T^, and comparing terms with Eq. S4, we see that the per-capita fitness *w*_*i*_ of each type is:

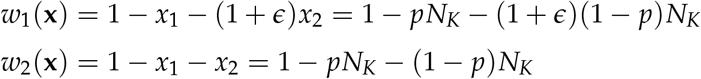

where *N*_*K*_ = *x*_1_ + *x*_2_ is the (scaled) total population size and *p* = *x*_1_/*N*_*K*_ is the frequency of type 1 individuals in the population. In frequency space, we thus see that the selection coefficient *s* := *w*_1_ − *w*_2_ is given by

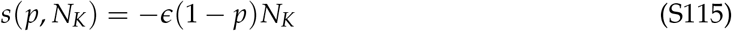

This calculation makes it clear that *ϵ* controls the strength and direction of natural selection operating in the system — when *ϵ* > 0, natural selection favors type 2, whereas when *ϵ* < 0, type 1 is favored. When *ϵ* = 0, the two types of birds have the same fitness and there is no natural selection operating in the system. If we now compute the per-capita turnover rates *τ*_*i*_ of each type, we have

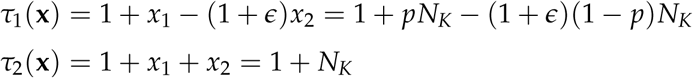

and the noise-induced selection coefficient *κ* := *τ*_2_ − *τ*_1_ is therefore

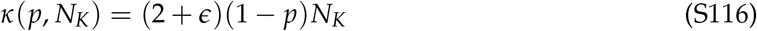

Note that when *ϵ* = 0, *s* vanishes but *κ* does not, meaning that the system exhibits noise-induced selection but no natural selection. Further, whenever *ϵ* > 0 or *ϵ* < −2, *s* and *κ* have opposite signs, *i.e*. natural selection and noise-induced selection act in opposite directions. Here, focusing on the case *ϵ* > 0, we see from Eq. S115 that natural selection favors type 2, whereas Eq. S116 tells us that noise-induced selection favors type 1.

Finally, we also have *Q*_1_(**p**) = (1 − *p*), *Q*_2_(**p**) = *p*. Substituting all these functional forms into Eq. S29 now tells us (after some algebra) that the frequency of type 1 individuals in the population obeys the SDE

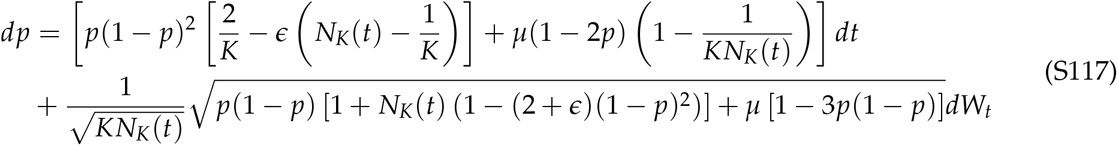

where *W*_*t*_ is a one-dimensional Wiener process. Upon substituting our functional forms of fitness and turnover into Eq. 7, we find that the total scaled total population size *N*_*K*_ obeys the SDE

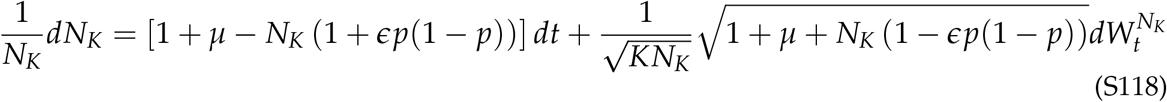

where 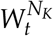 is a one-dimensional Wiener process. We are now in a position to study the behaviour of this system.

#### The infinite population limit

If we let *K* → ∞, the SDE for type frequency given by Eq. S117 reduces to an ODE

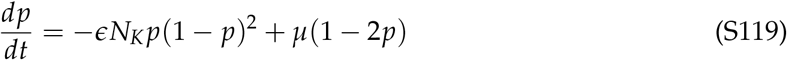

When there is no mutation and no selection in the system (*µ* = *ϵ* = 0), the RHS of Eq. S119 vanishes, and thus, any initial trait frequency *p*_0_ is expected to remain unchanged forever. If we switch off mutation alone (*µ* = 0, *ϵ* ≠ 0), it is easy to check that the type favoured by selection will become fixed in the population. Instead, if we switch off selection alone (*µ* ≠ 0, *ϵ* = 0), mutations drive the population to a state in which both types are equally prevalent (i.e., *p* = 0.5). When both selection and mutation are present in the system, the stable fixed point in the infinite population limit will lie in (0, 1/2) when *ϵ* > 0, and will lie in (1/2, 1) when *ϵ* < 0.

#### Deviations from neutrality due to noise-induced selection in finite, fluctuating populations

The effects of noise-induced selection on the expected dynamics are clearest when there is no natural selection (*ϵ* = 0) and no mutation (*µ* = 0): In this case, the equation for trait frequencies (Eq. S117) becomes

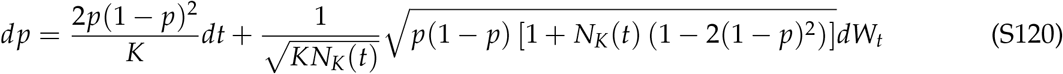

If we now take expectations on both sides, the stochastic integral term vanishes and we obtain an ODE for the expected trait frequency in the population. This ODE takes the form

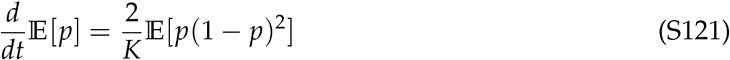

Since the RHS is always positive for *p* ∈ (0, 1), we conclude that the frequency of type 1 birds is always expected to increase until type 1 becomes fixed in the population. Thus, noise-induced selection, in this case, has led to a deviation from evolutionary neutrality in the expected dynamics — in the infinite population case, any initial trait frequency *p*_0_ is expected to remain unchanged forever, whereas for finite, fluctuating populations, assuming *p*_0_ ∉ {0, 1}, the trait frequency of type 1 birds is expected to increase until type 1 eventually fixes in the population. Note that unlike in neutral drift, type 1 is *always* expected to be the type that becomes fixed in the population despite the two types having the same fitness.

#### Reversal of the direction of evolution in finite, fluctuating populations

For the birth-death processes of the type we study in this paper, the entire population will go extinct in finite time with probability 1 (Ethier and Kurtz, 1986). Thus, the true stationary distribution for our system is thus the trivial state *x*_1_ = *x*_2_ = 0, a state at which *p* is undefined. However, the expected time to extinction is often so large as to be biologically meaningless, and in such cases, we can instead speak of the ‘quasi-stationary distribution’ of the stochastic process, obtained by only examining the system before the population goes extinct (Ethier and Kurtz, 1986; Karlin and Taylor, 1981). Thus, we are interested in the behaviour of the system in (*p, N*_*K*_) space conditioned on *N*_*K*_ > 0. To study the behaviour of the trait frequency when the population is far from extinction, we simply use the naive assumption *N*_*K*_ ≡ 1 to arrive at an approximate description of the system. As an aside, we note that better approximations could be made via the so-called ‘weak noise approximation’ or ‘linear noise approximation’ (Van Kampen, 1981; Black and McKane, 2012), though we do not do so here. Under the approximation *N*_*K*_ ≡ 1, the speed density *m*(*p*) is given by (see Supplementary section S6)

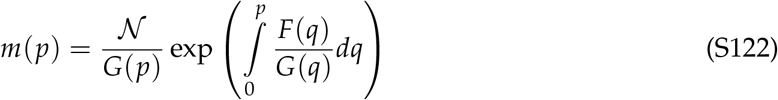

where 𝒩 is a normalization constant and *F* and *G* are functions given by

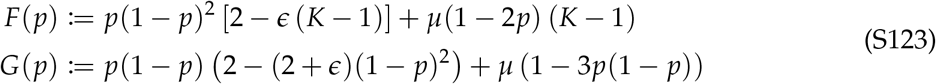

Since the above solution is an approximation, we also conduct exact stochastic individual-based simulations of the complete system defined by Eq. S114 using the Gillespie algorithm. The results of the simulations, as well as the solution predicted by Eq. S122, are plotted for a small *ϵ* > 0 (corresponding to weak selection against type 1) in figure S2.

For low values of *K* and *ϵ*, both the stochastic individual-based simulations and the approximate solution given by Eq. S122 indicate that noise-induced selection causes the distribution of types in the population to be biased in favor of type 1 (rightmost peak in Fig S2A) despite natural selection-mutation balance predicting a polymorphism in which type 2 individuals are over-represented. This bias disappears when *K* and *ϵ* are high, i.e. populations are large and natural selection is strong (Fig S2B).

**Figure S2:**
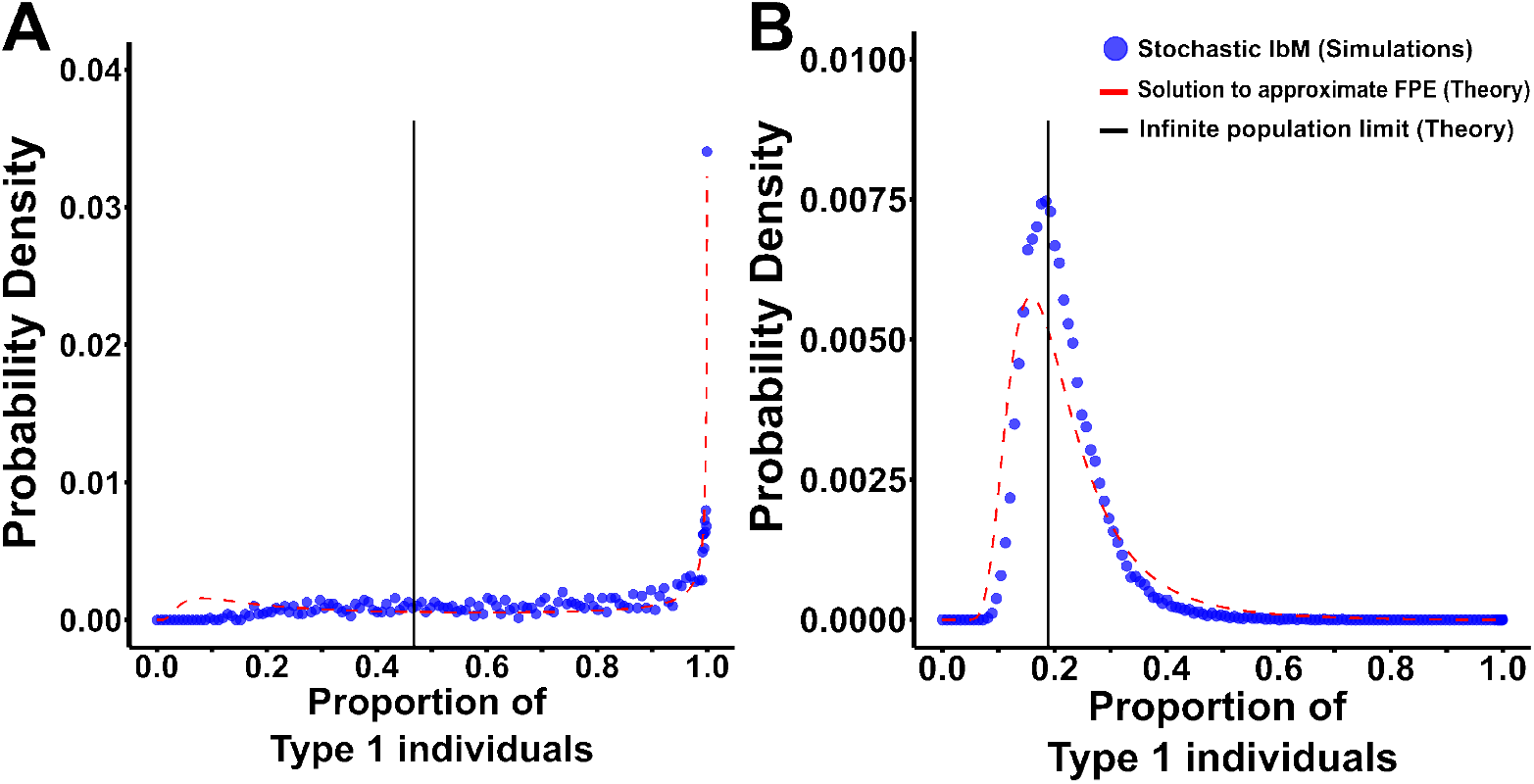
Predictions of our resource-competition model for various parameters. The quasistationary distribution has been plotted for **(A)** *K* = 500, *ϵ* = 0.0005, and **(B)** *K* = 5000, *ϵ* = 0.005. Blue points are from 100 independent Gillespie simulations of the exact birth-death process defined by Eq. S114, each supplied with the initial condition *n*_1_ = *n*_2_ = *K*/2 and allowed to run for 10^5^ timesteps or until the complete population went extinct. The red dotted line is derived from numerically evaluating the RHS of equation Eq. S122. The solid black line is the infinite population limit, obtained by solving equation Eq. S119 under the approximation *N*_*K*_ ≡ 1. For all plots in this figure, *µ* = 0.01.

## Notes

### Competing Interest Statement

The authors have declared no competing interest.

### Summary of Updates

Some terminology has changed ('slow'/'fast' to 'indirect'/'direct' and 'noise-induced selection' to 'noise-induced biasing'). Some text has been moved from the main text to either the Supplementary or dedicated boxes.

## Literature Cited

Abu Awad, D. and Coron, C. (2018). “Effects of demographic stochasticity and life-history strategies on times and probabilities to fixation”. In: Heredity 121.4, pp. 374–386. issn: 1365-2540. doi: 10.1038/s41437-018-0118-6.

Bhat, A. S. (2024). A stochastic field theory for the evolution of quantitative traits in finite populations. 2406.10739.

Black, A. J. and McKane, A. J. (2012). “Stochastic formulation of ecological models and their applications”. In: Trends in Ecology & Evolution 27.6, pp. 337–345. issn: 0169-5347. doi: 10.1016/j.tree.2012.01.014.

Boettiger, C. (2018). “From noise to knowledge: how randomness generates novel phenomena and reveals information”. In: Ecology Letters 21.8, pp. 1255–1267. issn: 1461-0248. doi: 10.1111/ele.13085.

Chapman, T., Arnqvist, G., Bangham, J., and Rowe, L. (2003). “Sexual conflict”. In: Trends in Ecology & Evolution 18.1, pp. 41–47. issn: 0169-5347. doi: 10.1016/S0169-5347(02)00004-6.

Chavhan, Y., Ali, S. I., and Dey, S. (2019). “Larger Numbers Can Impede Adaptation in Asexual Populations despite Entailing Greater Genetic Variation”. In: Evolutionary Biology 46.1, pp. 1–13. issn: 1934-2845. doi: 10.1007/s11692-018-9467-6.

Chavhan, Y., Malusare, S., and Dey, S. (2021). “Interplay of population size and environmental fluctuations: A new explanation for fitness cost rarity in asexuals”. In: Ecology Letters 24.9, pp. 1943–1954. issn: 1461-0248. doi: 10.1111/ele.13831.

Childs, D. Z., Metcalf, C. J. E., and Rees, M. (2010). “Evolutionary bet-hedging in the real world: empirical evidence and challenges revealed by plants”. In: Proceedings of the Royal Society B: Biological Sciences 277.1697, pp. 3055–3064. doi: 10.1098/rspb.2010.0707.

Chotibut, T. and Nelson, D. R. (2015). “Evolutionary dynamics with fluctuating population sizes and strong mutualism”. In: Physical Review E 92.2, p. 022718. doi: 10.1103/PhysRevE.92.022718.

Collet, P., Martínez, S., and San Martín, J. (2013). Quasi-Stationary Distributions: Markov Chains, Diffusions and Dynamical Systems. Probability and Its Applications. Berlin, Heidelberg: Springer Berlin Heidelberg. isbn: 978-3-642-33130-5. doi: 10.1007/978-3-642-33131-2.

Constable, G. W. A., Rogers, T., McKane, A. J., and Tarnita, C. E. (2016). “Demographic noise can reverse the direction of deterministic selection”. In: Proceedings of the National Academy of Sciences 113.32, E4745–E4754. doi: 10.1073/pnas.1603693113.

Crow, J. F. and Kimura, M. (1970). An introduction to population genetics theory. Harper international editions. New York: Harper & Row.

Czuppon, P. and Traulsen, A. (2021). “Understanding evolutionary and ecological dynamics using a continuum limit”. In: Ecology and Evolution 11.11, pp. 5857–5873. issn: 2045-7758. doi: 10.1002/ece3.7205.

Day, T., Parsons, T. L., Lambert, A., and Gandon, S. (2020). “The Price equation and evolutionary epidemiology”. In: Philosophical Transactions of the Royal Society B: Biological Sciences 375.1797, p. 20190357. doi: 10.1098/rstb.2019.0357.

Débarre, F. and Otto, S. P. (2016). “Evolutionary dynamics of a quantitative trait in a finite asexual population”. In: Theoretical Population Biology 108, pp. 75–88. issn: 0040-5809. doi: 10.1016/j.tpb.2015.12.002.

DeLong, J. P. and Cressler, C. E. (2023). “Stochasticity directs adaptive evolution toward nonequilibrium evolutionary attractors”. In: Ecology 104.1, e3873. issn: 1939-9170. doi: 10.1002/ecy.3873.

Doebeli, M. (2011). Adaptive diversification. Princeton, N.J.: Princeton University Press. isbn: 978-0-691-12894-8.

Doebeli, M., Ispolatov, Y., and Simon, B. (2017). “Towards a mechanistic foundation of evolutionary theory”. In: eLife 6. Ed. by Wenying Shou, e23804. issn: 2050-084X. doi: 10.7554/eLife.23804.

Engen, S., Bakke, Ø., and Islam, A. (1998). “Demographic and Environmental Stochasticity-Concepts and Definitions”. In: Biometrics 54.3, pp. 840–846. issn: 0006-341X. doi: 10.2307/2533838.

Ethier, S. N. and Kurtz, T. G. (1986). Markov processes: characterization and convergence. Wiley series in probability and mathematical statistics. New York: Wiley. isbn: 978-0-471-08186-9.

Ewens, W. J. (2004). Mathematical Population Genetics 1: Theoretical Introduction. Springer Science & Business Media. isbn: 978-0-387-20191-7.

Frank, S. A. (2012). “Natural selection. IV. The Price equation*”. In: Journal of Evolutionary Biology 25.6, pp. 1002–1019. issn: 1420-9101. doi: 10.1111/j.1420-9101.2012.02498.x.

Frank, S. A. and Slatkin, M. (1990). “Evolution in a Variable Environment”. In: The American Naturalist 136.2, pp. 244–260. issn: 0003-0147. doi: 10.1086/285094.

Frank, S. A. and Slatkin, M. (1992). “Fisher’s fundamental theorem of natural selection”. In: Trends in Ecology & Evolution 7.3, pp. 92–95. issn: 0169-5347. doi: 10.1016/0169-5347(92)90248-A.

Fronhofer, E. A., Corenblit, D., Deshpande, J. N., Govaert, L., Huneman, P., Viard, F., Jarne, P., and Puijalon, S. (2023). “Eco-evolution from deep time to contemporary dynamics: The role of timescales and rate modulators”. In: Ecology Letters 26.S1, S91–S108. issn: 1461-0248. doi: 10.1111/ele.14222.

Gillespie, J. H. (1974). “Natural selection for within-generation variance in offspring number”. In: Genetics 76.3, pp. 601–606. issn: 1943-2631. doi: 10.1093/genetics/76.3.601.

Gillespie, J. H. (1977). “Natural Selection for Variances in Offspring Numbers: A New Evolutionary Principle”. In: The American Naturalist 111.981, pp. 1010–1014. issn: 0003-0147. doi: 10.1086/283230.

Gokhale, C. S. and Hauert, C. (2016). “Eco-evolutionary dynamics of social dilemmas”. In: Theoretical Population Biology 111, pp. 28–42. issn: 0040-5809. doi: 10.1016/j.tpb.2016.05.005.

Hamilton, W. D. (1966). “The moulding of senescence by natural selection”. In: Journal of Theoretical Biology 12.1, pp. 12–45. issn: 0022-5193. doi: 10.1016/0022-5193(66)90184-6.

Horsthemke, W. and Lefever, R. (1984). Noise-induced transitions: theory and applications in physics, chemistry and biology. Springer series in synergetics. Berlin: Springer-Verlag. isbn: 978-3-540-11359-1.

Houchmandzadeh, B. and Vallade, M. (2012). “Selection for altruism through random drift in variable size populations”. In: BMC Evolutionary Biology 12.1, p. 61. issn: 1471-2148. doi: 10.1186/1471-2148-12-61.

Humplik, J., Hill, A. L., and Nowak, M. A. (2014). “Evolutionary dynamics of infectious diseases in finite populations”. In: Journal of Theoretical Biology 360, pp. 149–162. issn: 0022-5193. doi: 10.1016/j.jtbi.2014.06.039.

Jhawar, J., Morris, R. G., Amith-Kumar, U. R., Danny Raj, M., Rogers, T., Rajendran, H., and Guttal, V. (2020). “Noise-induced schooling of fish”. In: Nature Physics 16.4, pp. 488–493. issn: 1745-2481. doi: 10.1038/s41567-020-0787-y.

Johansson, J. and Ripa, J. (2006). “Will Sympatric Speciation Fail due to Stochastic Competitive Exclusion?”In: The American Naturalist 168.4, pp. 572–578. issn: 0003-0147. doi: 10.1086/507996.

Joshi, J. and Guttal, V. (2018). “Demographic noise and cost of greenbeard can facilitate greenbeard cooperation”. In: Evolution 72.12, pp. 2595–2607. issn: 1558-5646. doi: 10.1111/evo.13615.

Karlin, S. and Taylor, H. M. (1981). A second course in stochastic processes. New York: Academic Press. isbn: 978-0-12-398650-4.

Kimura, M. and Ohta, T. (1974). “Probability of Gene Fixation in an Expanding Finite Population”. In: Proceedings of the National Academy of Sciences 71.9, pp. 3377–3379. doi: 10.1073/pnas.71.9.3377.

Kogan, O., Khasin, M., Meerson, B., Schneider, D., and Myers, C. R. (2014). “Two-strain competition in quasineutral stochastic disease dynamics”. In: Physical Review E 90.4, p. 042149. doi: 10.1103/PhysRevE.90.042149.

Kokko, H. (2021). “The stagnation paradox: the ever-improving but (more or less) stationary population fitness”. In: Proceedings of the Royal Society B: Biological Sciences 288.1963, p. 20212145.doi: 10.1098/rspb.2021.2145.

Kuosmanen, T., Särkkä, S., and Mustonen, V. (2022). Turnover shapes evolution of birth and death rates. doi: 10.1101/2022.07.11.499527.

Lambert, A. (2010). “Population genetics, ecology and the size of populations”. In: Journal of Mathematical Biology 60.3, pp. 469–472. issn: 1432-1416.doi: 10.1007/s00285-009-0286-3.

Lande, R. (1976). “Natural Selection and Random Genetic Drift in Phenotypic Evolution”. In: Evolution 30.2, pp. 314–334. issn: 0014-3820. doi: 10.2307/2407703.

Lande, R. (1993). “Risks of Population Extinction from Demographic and Environmental Stochasticity and Random Catastrophes”. In: The American Naturalist 142.6, pp. 911–927. issn: 0003-0147.

Lehtonen, J. (2018). “The Price Equation, Gradient Dynamics, and Continuous Trait Game Theory”. In: The American Naturalist 191.1, pp. 146–153. issn: 0003-0147. doi: 10.1086/694891.

Lehtonen, J.(2020a). “The Price equation and the unity of social evolution theory”. In: Philosophical Transactions of the Royal Society B: Biological Sciences 375.1797, p. 20190362.doi: 10.1098/rstb.2019.0362.

Lehtonen, J.(2020b). “Longevity and the drift barrier: Bridging the gap between Medawar and Hamilton”. In: Evolution Letters 4.4, pp. 382–393. issn: 2056-3744. doi: 10.1002/evl3.173.

Lion, S. (2018). “Theoretical Approaches in Evolutionary Ecology: Environmental Feedback as a Unifying Perspective”. In: The American Naturalist 191.1, pp. 21–44. issn: 0003-0147.doi: 10.1086/694865.

Luque, V. J. and Baravalle, L. (2021). “The mirror of physics: on how the Price equation can unify evolutionary biology”. In: Synthese 199.5, pp. 12439–12462. issn: 1573-0964. doi: 10.1007/s11229-021-03339-6.

Majumder, S., Das, A., Kushal, A., Sankaran, S., and Guttal, V. (2021). “Finite-size effects, demographic noise, and ecosystem dynamics”. In: The European Physical Journal Special Topics 230.16, pp. 3389–3401. issn: 1951-6401.doi: 10.1140/epjs/s11734-021-00184-z.

Maklakov, A. A. and Chapman, T. (2019). “Evolution of ageing as a tangle of trade-offs: energy versus function”. In: Proceedings of the Royal Society B: Biological Sciences 286.1911, p. 20191604.doi: 10.1098/rspb.2019.1604.

Mallet, M. A., Bouchard, J. M., Kimber, C. M., and Chippindale, A. K. (2011). “Experimental mutation-accumulation on the X chromosome of Drosophila melanogaster reveals stronger selection on males than females”. In: BMC Evolutionary Biology 11.1, p. 156. issn: 1471-2148. doi: 10.1186/1471-2148-11-156.

Mazzolini, A. and Grilli, J. (2023). “Universality of evolutionary trajectories under arbitrary forms of self-limitation and competition”. In: Physical Review E 108.3, p. 034406.doi: 10.1103/PhysRevE.108.034406.

McLeod, D. V. and Day, T. (2019a). “Social evolution under demographic stochasticity”. In: PLOS Computational Biology 15.2, e1006739. issn: 1553-7358.doi: 10.1371/journal.pcbi.1006739.

McLeod, D. V. and Day, T. (2019b). “Why is sterility virulence most common in sexually transmitted infections? Examining the role of epidemiology”. In: Evolution 73.5, pp. 872–882. issn: 0014-3820. doi: 10.1111/evo.13718.

Metcalf, C. J. E. and Pavard, S. (2007). “Why evolutionary biologists should be demographers”. In: Trends in Ecology & Evolution 22.4, pp. 205–212. issn: 0169-5347.doi: 10.1016/j.tree.2006.12.001.

Olofsson, H., Ripa, J., and Jonzén, N. (2009). “Bet-hedging as an evolutionary game: the trade-off between egg size and number”. In: Proceedings of the Royal Society B: Biological Sciences 276.1669, pp. 2963–2969.doi: 10.1098/rspb.2009.0500.

Page, K. M. and Nowak, M. A. (2002). “Unifying Evolutionary Dynamics”. In: Journal of Theoretical Biology 219.1, pp. 93–98. issn: 0022-5193.doi: 10.1006/jtbi.2002.3112.

Papkou, A., Gokhale, C. S., Traulsen, A., and Schulenburg, H. (2016). “Host–parasite coevolution: why changing population size matters”. In: Zoology. SI: Host-Parasite Coevolution 119.4, pp. 330–338. issn: 0944-2006.doi: 10.1016/j.zool.2016.02.001.

Parsons, T. L., Lambert, A., Day, T., and Gandon, S. (2018). “Pathogen evolution in finite populations: slow and steady spreads the best”. In: Journal of The Royal Society Interface 15.147, p. 20180135. doi: 10.1098/rsif.2018.0135.

Parsons, T. L. and Quince, C. (2007). “Fixation in haploid populations exhibiting density dependence II: The quasi-neutral case”. In: Theoretical Population Biology 72.4, pp. 468–479. issn: 0040-5809.doi: 10.1016/j.tpb.2007.04.002.

Parsons, T. L., Quince, C., and Plotkin, J. B. (2010). “Some Consequences of Demographic Stochasticity in Population Genetics”. In: Genetics 185.4, pp. 1345–1354. issn: 1943-2631. doi: 10.1534/genetics.110.115030.

Proulx, S. R. and Day, T. (2005). “What can Invasion Analyses Tell us about Evolution under Stochasticity in Finite Populations?” In: Selection 2.1-2, pp. 2–15. issn: 1585-1931, 1588-287X.doi: 10.1556/select.2.2001.1-2.2.

Queller, D. C. (2017). “Fundamental Theorems of Evolution”. In: The American Naturalist 189.4, pp. 345–353. issn: 0003-0147. doi: 10.1086/690937.

Raatz, M. and Traulsen, A. (2023). “Promoting extinction or minimizing growth? The impact of treatment on trait trajectories in evolving populations”. In: Evolution 77.6, pp. 1408–1421. issn: 0014-3820. doi: 10.1093/evolut/qpad042.

Rice, S. H. (2004). Evolutionary theory: mathematical and conceptual foundations. Sunderland, Mass., USA: Sinauer Associates. isbn: 978-0-87893-702-8.

Rice, S. H. (2020). “Universal rules for the interaction of selection and transmission in evolution”. In: Philosophical Transactions of the Royal Society B: Biological Sciences 375.1797, p. 20190353.doi: 10.1098/rstb.2019.0353.

Rogers, T. and McKane, A. J. (2015). “Modes of competition and the fitness of evolved populations”. In: Physical Review E 92.3, p. 032708. doi: 10.1103/PhysRevE.92.032708.

Saunders, P. A., Neuenschwander, S., and Perrin, N. (2018). “Sex chromosome turnovers and genetic drift: a simulation study”. In: Journal of Evolutionary Biology 31.9, pp. 1413–1419. issn: 1420-9101. doi: 10.1111/jeb.13336.

Seger, J. and Brockmann, H. J. (1987). “What is bet-hedging?” In: Oxford surveys in evolutionary biology. Ed. by P.H. Harvey and L Partridge. Vol. 4. Oxford University Press, pp. 182–211.

Shoemaker, L. G., Sullivan, L. L., Donohue, I., Cabral, J. S., Williams, R. J., Mayfield, M. M., Chase, J. M., Chu, C., Harpole, W. S., Huth, A., HilleRisLambers, J., James, A. R. M., Kraft, N. J. B., May, F., Muthukrishnan, R., Satterlee, S., Taubert, F., Wang, X., Wiegand, T., Yang, Q., and Abbott, K. C. (2020). “Integrating the underlying structure of stochasticity into community ecology”. In: Ecology 101.2, e02922. issn: 1939-9170. doi: 10.1002/ecy.2922.

Shpak, M. (2005). “Evolution of variance in offspring number: The effects of population size and migration”. In: Theory in Biosciences 124.1, pp. 65–85. issn: 1611-7530.doi: 10.1016/j.thbio.2005.05.003.

Shpak, M. (2007). “Selection Against Demographic Stochasticity in Age-Structured Populations”. In: Genetics 177.4, pp. 2181–2194. issn: 1943-2631. doi: 10.1534/genetics.107.080747.

Starrfelt, J. and Kokko, H. (2012). “Bet-hedging—a triple trade-off between means, variances and correlations”. In: Biological Reviews 87.3, pp. 742–755. issn: 1469-185X. doi: 10.1111/j.1469-185X.2012.00225.x.

Van Kampen, N. G. (1981). Stochastic processes in physics and chemistry. Amsterdam, New York, New York: North-Holland. isbn: 978-0-444-86200-6.

Veller, C., Muralidhar, P., Constable, G. W. A., and Nowak, M. A. (2017). “Drift-Induced Selection Between Male and Female Heterogamety”. In: Genetics 207.2, pp. 711–727. issn: 1943-2631. doi: 10.1534/genetics.117.300151.

Wang, G., Su, Q., Wang, L., and Plotkin, J. B. (2023). “Reproductive variance can drive behavioral dynamics”. In: Proceedings of the National Academy of Sciences 120.12, e2216218120.doi: 10.1073/pnas.2216218120.

Week, B., Nuismer, S. L., Harmon, L. J., and Krone, S. M. (2021). “A white noise approach to evolutionary ecology”. In: Journal of Theoretical Biology 521, p. 110660. issn: 0022-5193.doi: 10.1016/j.jtbi.2021.110660.

Wodarz, D., Goel, A., and Komarova, N. L. (2017). “Effect of cell cycle duration on somatic evolutionary dynamics”. In: Evolutionary Applications 10.10, pp. 1121–1129. issn: 1752-4571. doi: 10.1111/eva.12518.

Yamamichi, M., Ellner, S. P., and Hairston Jr., N. G. (2023). “Beyond simple adaptation: Incorporating other evolutionary processes and concepts into eco-evolutionary dynamics”. In: Ecology Letters 26.S1, S16–S21. issn: 1461-0248. doi: 10.1111/ele.14197.

## Literature Cited in Supplementary

Alexander, H. and Wahl, L. (2008). “Fixation Probabilities Depend on Life History: Fecundity, Generation time and Survival in a burst-death model”. In: Evolution 62.7, pp. 1600–1609. issn: 0014-3820. doi: 10.1111/j.1558-5646.2008.00396.x.

Champagnat, N., Ferriére, R., and Méléard, S. (2006). “Unifying evolutionary dynamics: From individual stochastic processes to macroscopic models”. In: Theoretical Population Biology 69.3, pp. 297–321. issn: 0040-5809.doi: 10.1016/j.tpb.2005.10.004.

Champagnat, N. and Villemonais, D. (2023). “General criteria for the study of quasi-stationarity”. In: Electronic Journal of Probability 28, pp. 1–84. doi: 10.1214/22-EJP880.

Chotibut, T. and Nelson, D. R. (2017). “Population Genetics with Fluctuating Population Sizes”. In: Journal of Statistical Physics 167.3, pp. 777–791. issn: 1572-9613. doi: 10.1007/s10955-017-1741-y.

Constable, G. W. A., McKane, A. J., and Rogers, T. (2013). “Stochastic dynamics on slow manifolds”. In: Journal of Physics A: Mathematical and Theoretical 46.29, p. 295002. issn: 1751-8121.doi: 10.1088/1751-8113/46/29/295002.

Day, T., Parsons, T. L., Lambert, A., and Gandon, S. (2020). “The Price equation and evolutionary epidemiology”. In: Philosophical Transactions of the Royal Society B: Biological Sciences 375.1797, p. 20190357.doi: 10.1098/rstb.2019.0357.

Etheridge, A. (2011). Some mathematical models from population genetics: École d’été de Probabilités de Saint-Flour XXXIX-2009. Lecture Notes in Mathematics. Berlin: Springer-Verlag. isbn: 978-3-642-16631-0.

Ethier, S. N. and Kurtz, T. G. (1986). Markov processes: characterization and convergence. Wiley series in probability and mathematical statistics. New York: Wiley.isbn: 978-0-471-08186-9.

Frank, S. A. (2012). “Natural selection. IV. The Price equation*”. In: Journal of Evolutionary Biology 25.6, pp. 1002–1019. issn: 1420-9101.doi: 10.1111/j.1420-9101.2012.02498.x.

Frank, S. A. and Slatkin, M. (1992). “Fisher’s fundamental theorem of natural selection”. In: Trends in Ecology & Evolution 7.3, pp. 92–95. issn: 0169-5347.doi: 10.1016/0169-5347(92)90248-A.

Fronhofer, E. A., Corenblit, D., Deshpande, J. N., Govaert, L., Huneman, P., Viard, F., Jarne, P., and Puijalon, S. (2023). “Eco-evolution from deep time to contemporary dynamics: The role of timescales and rate modulators”. In: Ecology Letters 26.S1, S91–S108. issn: 1461-0248.doi: 10.1111/ele.14222.

Hofbauer, J. and Sigmund, K. (1998). Evolutionary Games and Population Dynamics. Cambridge: Cambridge University Press. isbn: 978-0-521-62570-8.doi: 10.1017/CBO9781139173179.

Karatzas, I. and Shreve, S. E. (1998). Brownian motion and stochastic calculus. 2nd ed.Graduate texts in mathematics. New York: Springer-Verlag. isbn: 978-0-387-97655-6.

McLeod, D. V. and Day, T. (2019). “Social evolution under demographic stochasticity”. In: PLOS Computational Biology 15.2, e1006739. issn: 1553-7358.doi: 10.1371/journal.pcbi.1006739.

Øksendal, B. K. (1998). Stochastic differential equations: an introduction with applications. Berlin; New York: Springer. isbn: 978-3-662-03620-4.

Parsons, T. L. and Rogers, T. (2017). “Dimension reduction for stochastic dynamical systems forced onto a manifold by large drift: a constructive approach with examples from theoretical biology”. In: Journal of Physics A: Mathematical and Theoretical 50.41, p. 415601. issn: 1751-8121.doi: 10.1088/1751-8121/aa86c7.

Van Kampen, N. G. (1981). Stochastic processes in physics and chemistry.Amsterdam, New York, New York: North-Holland. isbn: 978-0-444-86200-6.

